# Complex cooperativity in DNA origami revealed *via* design dependent defectivity

**DOI:** 10.1101/2025.04.23.650292

**Authors:** Jacob Majikes, Amna Hasni, Shankar Haridas, Joey Robertson, Adam Pintar, Michael Zwolak, J. Alexander Liddle

**Affiliations:** Theiss Research, La Jolla, CA, USA; Microsystems and Nanotechnology Division, National Institute of Standards and Technology, Gaithersburg, MD, USA; Biomedical Engineering, Johns Hopkins University, Baltimore, MD, USA; University of Maryland, College Park, MD, USA

## Abstract

DNA origami has become a ubiquitous platform for nanostructure fabrication because it enables straightforward design of structures that self-assemble with high yield. The interactions between the cooperative effects involved in its assembly are currently not well understood. Fortunately, the nearly infinite number of choices available to the origami designer provide a rich environment in which to explore cooperativity in a complex system. The DNA domains comprising origami have predictable energetics and the sources of cooperativity are conceptually straightforward. While the assembly of these systems is difficult to predict because of the large number of cooperative interactions, it can be measured. We are thus able to probe cooperativity by using design variations and measuring their effect on assembly yield. We employ an accelerated assembly protocol that increases the sensitivity of structural perfection, or lack thereof, to design variation, and apply this approach to survey a broad set of design features. Using the resulting dataset, we develop metrics to correlate thermal stability, beneficial cooperativity from short folds, and detrimental cooperativity from long folds, with defectivity. Surprisingly, these metrics can be combined to create a single parameter with a clear correlation to yield that serves as a useful starting place for a predictive understanding of the interplay between cooperativity and design. In doing so, we also identify qualitative trends that provide useful insight into design best practice.

## Introduction

Cooperative effects are ubiquitous in biomacromolecular systems and drive emergent properties associated with complex systems like cell signaling^1^. Molecular nanofabrication employs networks of cooperative effects to steer self-assembling systems towards a single, final state. In short, cooperativity is nature’s answer to Levinthal’s paradox^2^. However, a full mechanistic understanding of how assembly events interact and progress would facilitate the rational design of systems capable of accessing multiple metastable states in response to complex environmental cues. Nucleic acid nanofabrication is an ideal platform for investigating these concepts.

Cooperativity is the propensity for identical, or similar, components in a system to modulate each other’s behavior. The archetypal example of cooperativity is the binding of multiple oxygen molecules by hemoglobin, where each binding event increases the binding strength of the subsequent one^3^, for which the well-known Hill parameter was originally developed. Unfortunately, this parameter, and other common metrics of cooperativity, are properties of a dose response curve and are not readily applicable to complex self-assembling systems. This concern is in many ways definitional; to quantify cooperativity one must identify what the system is cooperative in reference to. Clearer definitions with better specified reference states are easier to quantify, but harder to compare or combine across differing physical sources of cooperativity. For complex systems with multiple sources of cooperativity this necessitates iterative research cycles as individual sources are identified, their quantification devised, and their convolution evaluated against whole system behavior. Complex cooperative systems suffer from the classic curse of dimensionality.

DNA origami combines top-down design with bottom-up assembly and DNA hybridization’s well-understood energetics. The assembly of a single origami structure involves over 600 reversible hybridization reactions between domains spread across more than 200 multivalent oligomer strands (staples) and a primary continuous template strand (scaffold), all of which exhibiting a variety of cooperative energetic contributions, as depicted in Fig 1. The resulting myriad reaction pathways make *a priori* modeling exceptionally difficult, but also lead to robust assembly, with quoted yield values often in the range of 90 % – a surprisingly high fraction for such a complex series of reactions. Though modeling efforts continue, we cannot yet predict the effect of design on yield^4,5^ nor are current algorithmic design tools fully “yield aware”^6–9^. This is a highly relevant topic for industrial applications^10–12^, as the size of origami, 4·5.10^6^ g.mol^-1^ (4.5 MDa), precludes simple purification^11^ and limits potential application areas^13^.

**Fig 1.**
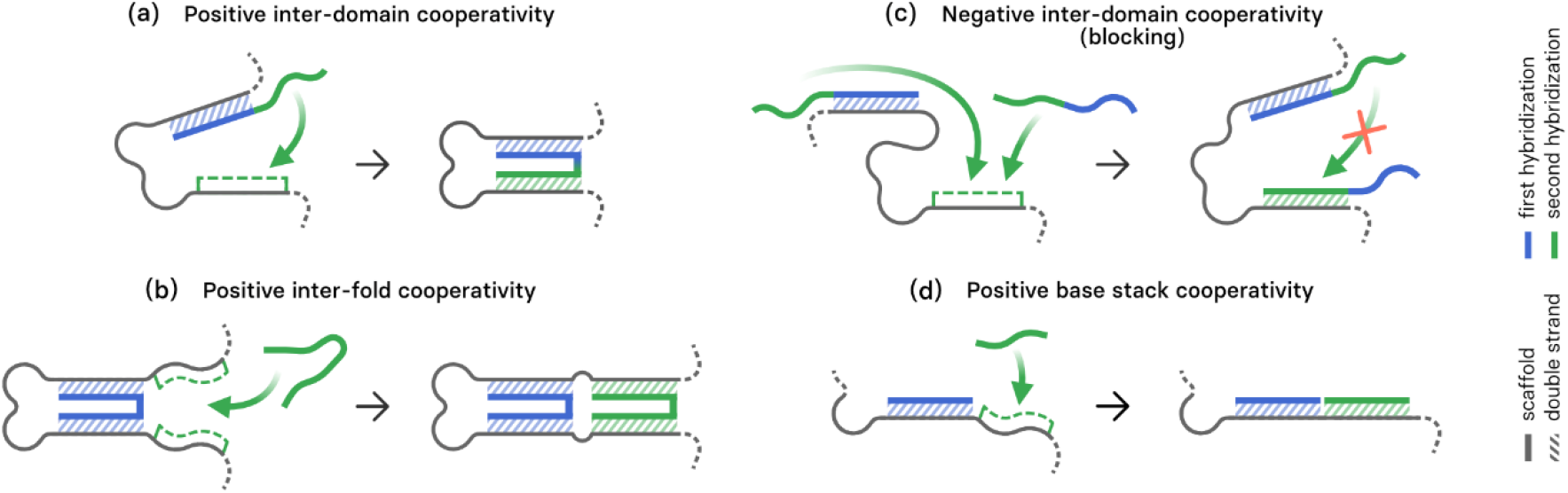
Schematic of the primary forms of cooperativity in DNA origami discussed in the literature. a) positive inter-domain cooperativity: at short scaffold distances, when a single domain on a staple binds, neighboring domains on that staple experience positive cooperativity as their effective local concentration is higher than their competitors in solution. This is reduced by loop entropy at longer distances, b) inter-fold cooperativity: at any fold distance, the closing of one fold reduces the possible conformations available to the entire remaining scaffold, stabilizing all other non-competing folds but most heavily affecting close neighbors c) negative inter-domain cooperativity, or blocking: at very long scaffold distances, the second domain on a staple can become less likely to bind than a competing domain on another copy in solution as the effective local concentration for the bound staple (quantified by loop entropy or j-factor) is lower than the solution concentration of the competitors. d) base stacking cooperativity: when an immediate neighboring domain has already bound there is an energetic bonus for π-π stacking between helix ends.

The four sources cooperativity we define in Fig 1 originate from a variety of observations from studies which varied a single design feature or measured a subset of an origami. These include the presence of a ‘nucleating’ domain on each staple^14,15^ (promoting inter-domain cooperativity by providing energetic heterogeneity^16,17^), the loop distance distribution^18–21^ (affecting inter-fold and inter-domain cooperativity and nucleation of folding^5,22^), high guanine and cytosine (GC) content at long loop distance folds (encouraging negative inter-domain cooperativity induced defects^23,24^), and the presence of high GC content at crossovers (creating a kinetic road bump through negative base stacking cooperativity^25^). Theoretical works have also indicated that the distribution of GC content, as bounded by rotations of a single scaffold sequence, are unlikely to affect assembly yield^22^. To understand full origami behavior, these sources of cooperativity must be quantified and weighed against each other. This is made more difficult still by their operation at different layers of conceptual abstraction, e.g., the cooperative events for inter-domain cooperativity are domain hybridization events while for inter-fold cooperativity are staple:scaffold folding events. However, doing so is a prerequisite to emulate aspects of protein behavior or to perform engineered chemo-mechanical work.

These proposals are consistent growing body of research on the mechanisms of DNA origami assembly. The current consensus is that origami fold *via* nucleation and growth^18,19,21,26^, in which loop entropy and base stacking contributions play a role^5,22^. Notable works have shown multiple paths for nucleation and growth in individual structures^19,26^ which explain other, less direct measurements of assembly^20,27^. Oligomer inclusion studies have indicated assembly in two to three distinct stages, for at least one 3D design^28^. While theoretical advances improve our ability to understand and predict the cooperativity of origami assembly^4,5^, we have yet to extend this understanding to evaluation of designs, or to relate general design rules to cooperative effects.

Additionally, one might anticipate that the relative contributions of each source of cooperativity might scale differently with origami size. For example, one might assume folds on the outer edge of an origami must always be short folds, while the longest folds will always be across some seam inside the center of the structure. One might then reasonably assume a geometric progression of fold size distribution like that of perimeter versus area. Across whole origami, we expect positive inter-domain and base stacking cooperativity, Fig 1 a & d, not to scale with size, while negative inter-domain cooperativity and positive inter-fold cooperativity, Fig 1 b & c, should, with the former scaling faster. This suggests higher defectivity for larger origami.

In origami the sources of cooperativity are mediated by where events occur as topology changes progress during assembly. This topology progression is controlled by origami design, and the origami design space is notoriously vast, where any one shape can be designed a nearly infinite number of ways^29,30^. A crude estimation of this space would consider changing the scaffold routing (including the direction of the helices through the structure, number of seams, and seam locations), rotation of the sequence through the routing, and the length and distribution of binding domains on the staples (*i*.*e*., staple motif), as shown in Fig. 2. Such an estimate, assuming only four choices for each of the above design variables, would suggest at least 4^5^, or 1 024, possible variants for a single 2D shape.

**Fig. 2.**
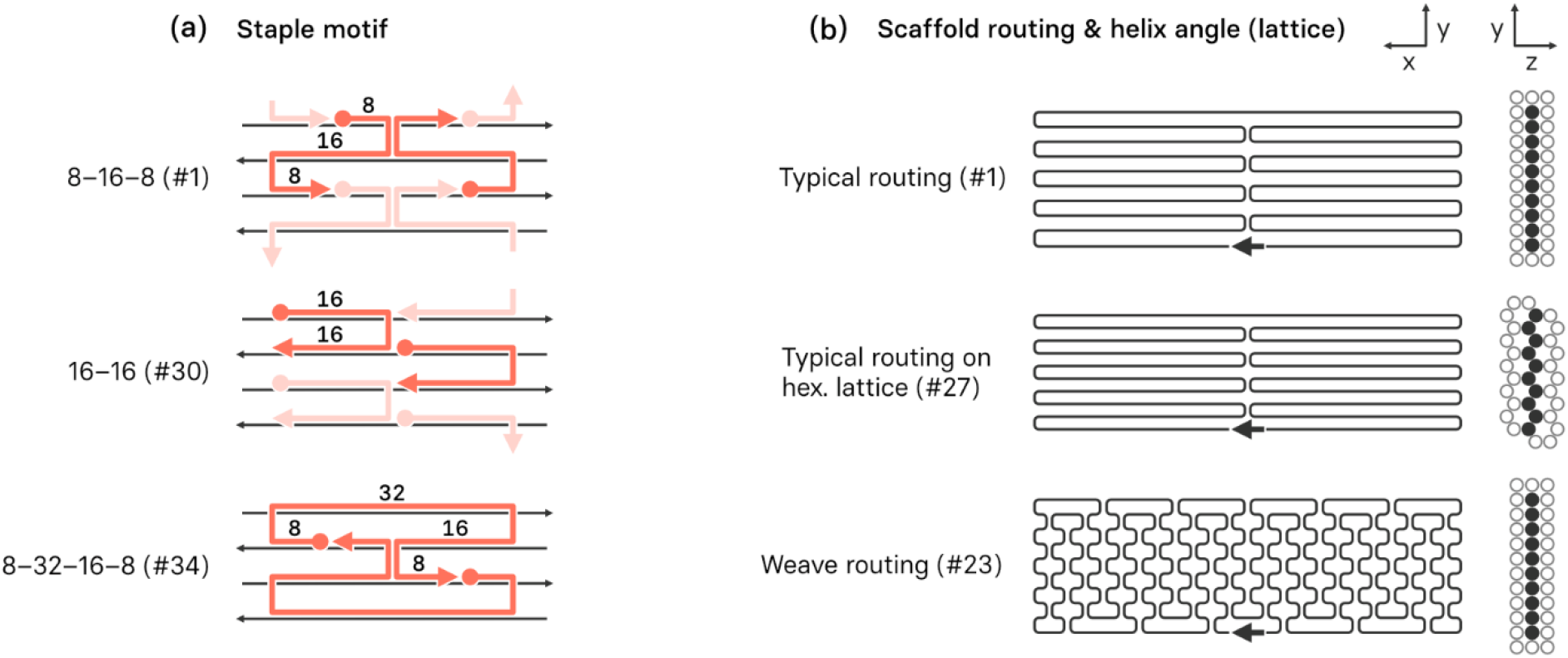
DNA origami design schematics emphasizing simple changes to design that dramatically alter the distribution of cooperative events. a) Examples of staple motifs showing the tiling of that motif. Motifs are named by the length and number of their domains. Note that changing the motif from 8-16-8 to 16-16 will convert the double to single crossovers, but not change the number or position of topological links on the scaffold. By contrast, changing from 8-16-8 to 8-32-16-8 will remove a fraction of topological links as the two staple tiling unit repeats vertically. b) Examples of scaffold routings and inter-helix angles. Changing the angle of helix connections, as between lattices, changes the location of viable crossovers and thus the possible staple motifs. Similarly, using scaffold, rather than staple, crossovers as in the weave routing will alter the folding pathways available for assembly. The design numbers (#1, #23, #27, #30, & #34) refer to specific designs (see Fig. 3). Note: the arrow in routing plots indicates the start of the scaffold sequence, as well as the position where any excess, unused, scaffold enters and exits the structure. In this study this ‘goatee’ of excess scaffold was approximately 209 bases in length.

This multiplicity of designs provides an exciting space for exploration, but unfortunately no change to design will vary any source of cooperativity independently. This, and the lack of metrics which quantify sources of cooperativity across whole designs, formulates a central challenge and gordian knot of this area of research. To confirm that all relevant sources of cooperativity are identified, one must deconvolve them from behavior of the whole origami, in order to deconvolve them one must devise quantitative metrics for each source, and to devise quantitative metrics for them one must first identify the relevant sources of cooperativity. *Identifying, devising, and deconvolving the sources of cooperativity present a trifold problem that is, at its core, an example of the curse of dimensionality that plagues any complex system*.

Yield is the most relevant measurand to leverage against this challenge as some formulation of it was used by the majority of the previously cited works that identified sources of origami cooperativity. However, quantitative yield measurements sufficient for deconvolving cooperative effects pose a much greater challenge than those necessary for more qualitative work. Yield is typically defined as the ratio of product to reactant: [product]/[reactant], but for an origami, the definition of “product” is ambiguous. For example, must the origami be perfect, in the sense that every designed staple is incorporated, or is it sufficient for it to assume the designed shape or have the desired functionality? If perfect staple incorporation is required, then one’s measurement must resolve a 0.17 % change in mass. In contrast, measuring geometric fidelity between an image and a structure’s design presents dramatic challenges to calibration and replication between laboratories.

At present, yield measurements have an inverse relationship between their resolution and both their cost and throughput. They can be categorized from low to high resolution: migratory yield (*e*.*g*., gel electrophoresis and chromatography)^14,31,32^, imaging yield (*e*.*g*., Atomic Force Microscopy, AFM, and Transmission Electron Microscopy, TEM)^19,26,33–36^, and oligomer inclusion yield (*e*.*g*., DNA-PAINT)^28,36–38^. For the purposes of our study, we select AFM imaging yield as the method with the optimum combination of resolution, *i*.*e*., the ability to determine a coarse-grained level of fidelity of an assembled 2D origami to its design, with a throughput high enough to provide adequate statistical confidence across many designs. To aid plotting and interpretation in this work, we chose to use the portion of defective structures rather than the yield of well folded ones, and will refer to yield in the context of defectivity through the rest of the manuscript.

To address these questions and explore the role of cooperativity in DNA origami assembly, we introduce an accelerated annealing protocol to facilitate more efficient extraction of data which dramatically improves the efficiency of time spent imaging each design. This allowed us to perform one of the broadest explorations to date of design variations within a fixed geometry. This provided the data needed to begin deconvolving cooperative contributions in this complex system. In so doing we have identified the most relevant contributors, and shed light on why the human aesthetic preference for symmetry and shorter seams is synonymous with high quality DNA origami designs.

## Results & Discussion

### Given the multiplicity of cooperative effects and their interconnectedness, it is essential to survey a wide variety of designs rather than explore the effects of incremental variations of a single design parameter, e.g., seam location

See Fig. 3 for a summary of the designs and SI Sections 4 & 17 for more detailed information about each. All the designs except the notched rectangle had a nearly identical aspect ratio, number of bases engaged in dsDNA, and had 2 unpaired thymine nucleotides of ssDNA at the end of each helix to minimize base stacking, where ssDNA had to loop between edge helices 5 unpaired thymine nucleotides were used. In some designs these ssDNA extensions were connected, which could explain the limited origami stacking observed in AFM images, as looped ssDNA may be less efficient per base at preventing favorable stacking effects^39^.

**Fig. 3:**
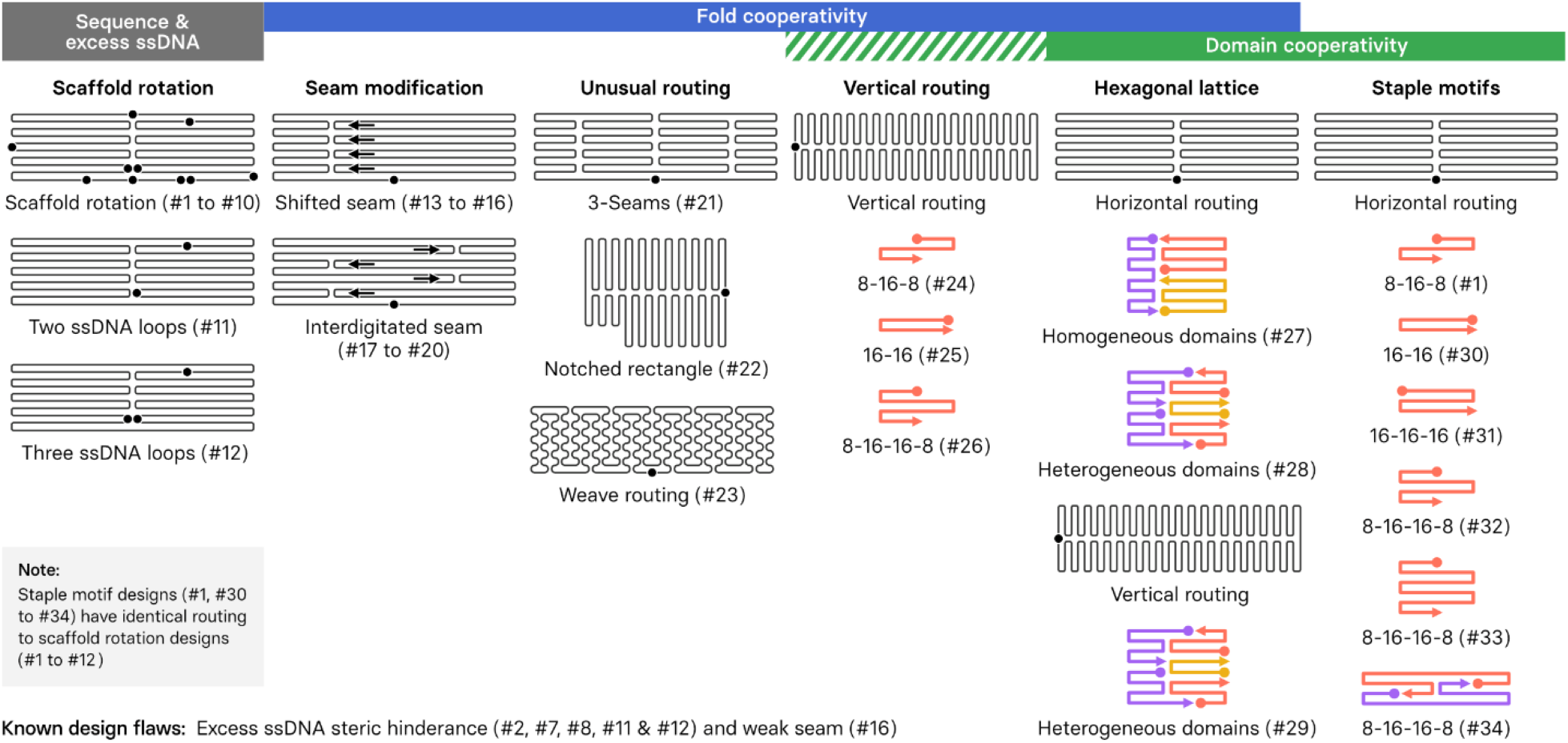
Design Survey. Each design is numbered #1 to #34 and comprises a scaffold routing with a sequence start position that also denotes the location of the excess ssDNA loop (labeled with a circle), an interhelical angle or lattice, and a staple motif. A tabular version of this, emphasizing comparable design features, is given in supplementary table S6. Designs #1 to #22 all had an 8-16-8 staple motif, and #23 had a 16-16 motif. The numeric order was assigned after analysis was completed to emphasize similarities and sources of cooperativity between designs.

To aid comparison, we varied designs relative to standard 2D origami features. These features for a typical design (design #1) are – helices connected at π/2 (90 °) angles (square lattice), that run parallel to the long direction of the structure, a single symmetric seam with the scaffold excess coiled at the center bottom of the structure, and an 8-16-8 staple motif. Although these design features cannot be fully varied independently of each other, we selected design variants to minimize the number of simultaneous changes.

To test the role of GC content and the excess ssDNA coil, we rotate the sequence start position, labeled as a dot in Fig. 3 and throughout the text. This is also annotated as −X or +Y nucleotides, where the negative value pulls the scaffold clockwise through the routing pattern. As the scaffold is circular, any rotation could be expressed by either a negative or positive number. We chose the smaller absolute value for our notation.

Our design variants comprised 6 staple motifs, 12 scaffold rotations, 8 designs which broke or modified the symmetry of the seam incrementally, 2 designs in which there were more than one seam, 4 designs in which the helices run perpendicular to the long direction of the rectangle, and finally we tested 3 designs in which the helices connected at π 2/3 rad (120°) angles typical of 3D designs rather than the π/2 rad (90°) typical of 2D designs. These last three designs had, by necessity, a mix of staple motifs, which were made to either have heterogenous or non-heterogeneous distributions of staple domain lengths.

Each source of cooperativity identified in the literature, Fig 1, is simple to describe and may be modeled on the level of an individual set of reactions. However, creating metrics to weight designs based on their relative propensity for each source of cooperativity is less straightforward. We describe our approach to doing so below.

For each source of cooperativity below, a more in-depth discussion is provided in SI Section 2, including alternate formulations.

Inter-domain cooperativity, Fig 1a, or the propensity staple domains to nucleate folds had several potential formulations. The one used here is the sum across all fold of the difference in predicted nearest neighbor T_m_ between the most and least stable staple domains normalized by the all-domain average T_m_.

Inter-fold cooperativity, Fig 1b, or reduction in conformational entropy penalties caused by neighboring folds, is straightforward to calculate for any two folds, but ill-defined across a whole origami. As for the previous source of cooperativity, we chose to represent it as a property of the distribution of possible folds. For this we chose the skew of the loop entropy penalty distribution for each fold assuming it was first to bind. The entropic penalty is asymptotic with increasing fold distances, ensuring that the skew is most sensitive to changes in the relevant short and medium folds. A distribution’s skew is also conveniently unitless. We chose this as we anticipate it will be correlated with how path-like or funnel-like the assembly process is.

Negative inter-domain cooperativity or blocking, Fig 1c, is quantified as in our previous work, which provides a useful frame of reference^24^. We calculate the portion of scaffolds that would bind two copies of the staple at thermodynamic equilibrium and our experimental concentrations, neglecting the effects of other staples. This assumes that there is an equilibrium between the folded and blocked states which balances the entropic penalty of folding with the entropy lost when a second staple copy binds.

Base stacking cooperativity, Fig 1d, was simply the sum across all domain terminuses of the enthalpy term of the nearest neighbor model for those terminal bases. We similarly summed enthalpies of base stacks at crossovers to examine the hypothesis by Cumberworth *et al*.^25^, see SI section 14.

Under typical anneal conditions (10 × to 1 000 × staple excess, 1 h to 12 h anneal from ≈ 80 °C to 4 °C) designs either fold into the desired shape with defect fractions on the order of a single-digit percentage, or not fold at all due to some catastrophic design flaw. Distinguishing small variations in defect fractions of a few percent with high confidence would require a prohibitive level of statistical sampling. For example, two designs which yield 90 % and 95 % well folded structures respectively would require classifying hundreds of structures each to ensure sampling uncertainty much lower than their difference in yield.

However, it requires far smaller sample sizes to distinguish between designs when the defective fractions are closer to 50 %. Analytical solutions of similar problems in cell counting serve as a useful example^40^. We therefore engineered our annealing conditions to increase both the defective fraction and its sensitivity to design changes. This approach is valid provided the mechanisms of defect formation are the same under poor annealing conditions as they are under typical ones. Our control experiments, SI section 6, indicate that the trends found under our poor annealing conditions will hold under more standard conditions.

To estimate defectivity, we classified origami into easily distinguished categories, which we defined as well-folded, damaged, ripped, and catastrophic (Fig. 4a and 4b, and SI section 6 & 7).

**Fig. 4.**
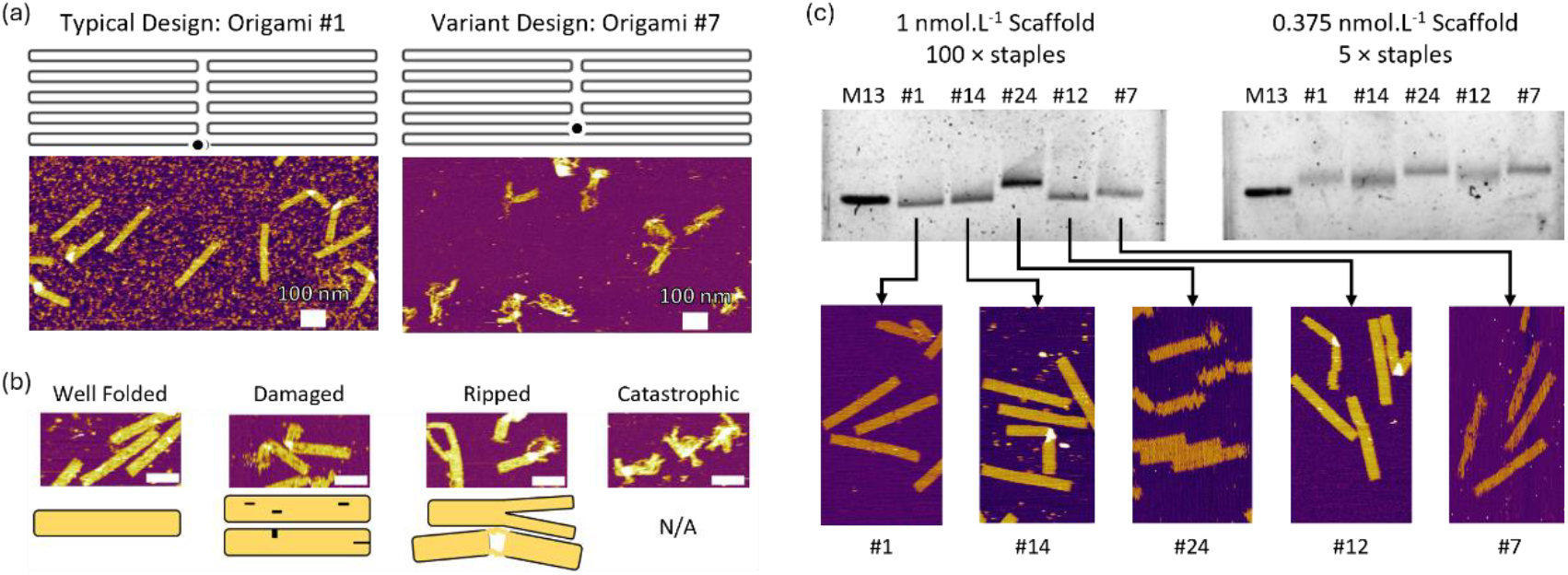
a) Example AFM images and routing pattern of one of the designs with the high yield under accelarated test conditions, design #1, and one of the worse performing designs, # 7. b) Classifications, AFM examples, and schematics. All scale bars 100 nm. (c) Gel images, in 1 % agarose, 100 V, 2 h, where the staple band is cropped out of the image. Bottom, AFM images of the example structures annealed at 100 × staple:scaffold and imaged on mica in liquid tapping modes

Well-folded structures exhibit no visible defects; damaged structures display one to two visible defects that did not change the overall geometry; ripped structures are ones whose integrity is obviously reduced, but which are still more assembled than not; and catastrophic origami have no recognizable structure. In lieu of ground-truth data sets of sufficient size to enable machine learning approaches, we use human analysts to classify the images.

We equate the defectivity, or ϕ, to the average number of missing or blocked staples for a design. The coefficients in Eq. 1, correspond to an estimate of the minimum number of defective staples necessary for a given level of visible damage with 1 staple, 5 staples, 15 staples, and 50 staples corresponding to the well-folded, damaged, ripped, and catastrophic classifications respectively. By choosing only four classifications we ensured each classification was sufficiently different from the others that the trends in the defectivity were robust to the exact coefficients used in Eq. 1.

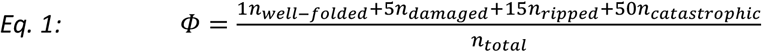

Our observed trends in ϕ were robust to differences in the exact coefficients because the classification categories were sufficiently distinct. However, we caution the reader to use this only as a very approximate substitute for the number of missing or blocked staples. The primary utility of ϕ is to facilitate comparison between designs.

As a control, we ran 1 % agarose gels of some of our best and worst performing structures. In Fig. 4c, we show separate gels of origami annealed under typical high-yield conditions on the left, and under our accelerated conditions on the right. The lower section of Fig. 4c shows AFM images from the high-yield condition and illustrates the difficulty in using gel electrophoresis for high-precision measurements of the defective fraction formed in an origami assembly. First, the origami are sufficiently large that only a significant amount of damage will shift the band; under high-yield anneal conditions, design #7 had damage in its AFM image, but exhibited a single clear band of roughly the same mobility as the well-folded designs, #1 and #14. Second, design variations can significantly shift the mobility even for well-folded structures: for example, designs #1, #14, and #24 were all reasonably well-folded under high-yield anneal conditions but had different mobilities. Third, it is difficult to self-consistently quantify band shifts of such large structures; the origami bands associated with accelerated anneal conditions migrated more slowly relative to the M13 control than their high-yield anneal condition counterparts, indicating damage. However, quantifying the degree of that damage would require intensive modeling of the electrophoretic mobility of each design.

However, other studies with a more constrained parameter space can make good use of migratory yield measurements. For example, 3D origami have dramatically larger band shifts for poorly folded structures and tend to fail more catastrophically than 2D ones. Studies concerning catastrophic levels of defects in 3D structures typically find gel electrophoresis sufficient^6,14,15^. Similarly, if a single design is being used to probe annealing condition optimization or nuclease degradation, band shifts for that design are more readily interpretable^41,42^.

### Classification and Uncertainty

We consider several sources of uncertainty in imaging yield: the variation between analysts in classifying origami; the sample size and statistics associated with calculating ϕ; the day-to-day variation in analyst classification; and variation in the synthesis yield of the staple pools for each design. These were quantified via control experiments and Monte Carlo simulation, all uncertainties presented here represent one standard deviation.

Variation between analysists was evaluated for a subset of our initial designs and evaluated as a consistency check, see SI section 6. While analysts disagreed on the exact defectivity value, there was agreement in the trend between designs for all analysts.

The three remaining uncertainty sources were quantified by a mix of control experiments and Monte Carlo simulation, discussed in section 8 of the SI. Our designs exhibited a range of defectivity, ϕ, from ≈ 10 to ≈ 35. Surprisingly, we found that variation in staple pool synthesis contributes the most to uncertainty in defectivity, where a single standard deviation contributed ± 2.8 in ϕ in the control. Sample size and day-to-day variation contributed less for a single standard deviation, from ± 0.2 to ± 1.7 in ϕ for the former and ± 0.1 to ± 0.3 in ϕ for the latter. Future developers of protocols who wish to optimize imaging yield experiments may consider imaging fewer origami from multiple pools of the same design to dramatically reduce the overall uncertainty for similar person-hour costs.

### Qualitative trends in defectivity as a function of design

Outside of our direct goal of probing cooperativity, several interesting qualitative trends in defectivity are observable within the families of design we examined. We detail these trends and the evidence for them in section 9 of the SI, and summarize them here as follows.

When the scaffold sequence was rotated in ways that kept the loop of excess scaffold (approximately 200 nucleotides) on the top or bottom edge, we observed no change in defectivity. When the excess loop of scaffold was internal to the structure or positioned to increase the loop distance of an entire row of staples, the defectivity dramatically increased. These designs are depicted as hollow circles in the following figures and are labeled as flawed due to catastrophic steric effects. We observed significant changes for modified staple motifs, but without a clear interpretation for these changes. This is likely because the motifs affect multiple sources of cooperativity and will have additional changes in staple pool synthesis variation due to their different staple lengths. Finally, altering the seam symmetry consistently increased defectivity, in spite of our hypothesis that interdigitated seams should improve yield. We attribute this to counterintuitive changes in the distribution of fold distances, as discussed in SI section 10.

Taken together, these results indicate that origami designers are well served by their aesthetic biases. Keeping symmetric seams, keeping staples short, keeping excess scaffold on the outside of the structure and using 8-16-8 motifs were effective approaches to reduce defectivity in the systems we examined.

Before examining combinations of metrics, we plot each metric against defectivity, Fig. 5Fig. 5 a to d. If a clear trend is visible, it indicates that the metric is strongly correlated with defectivity. However, the absence of a clear trend does not necessarily mean that a metric is uncorrelated with defectivity, but that a correlation may be obscured by collapsing a multi-dimensional space onto a single axis.

**Fig. 5:**
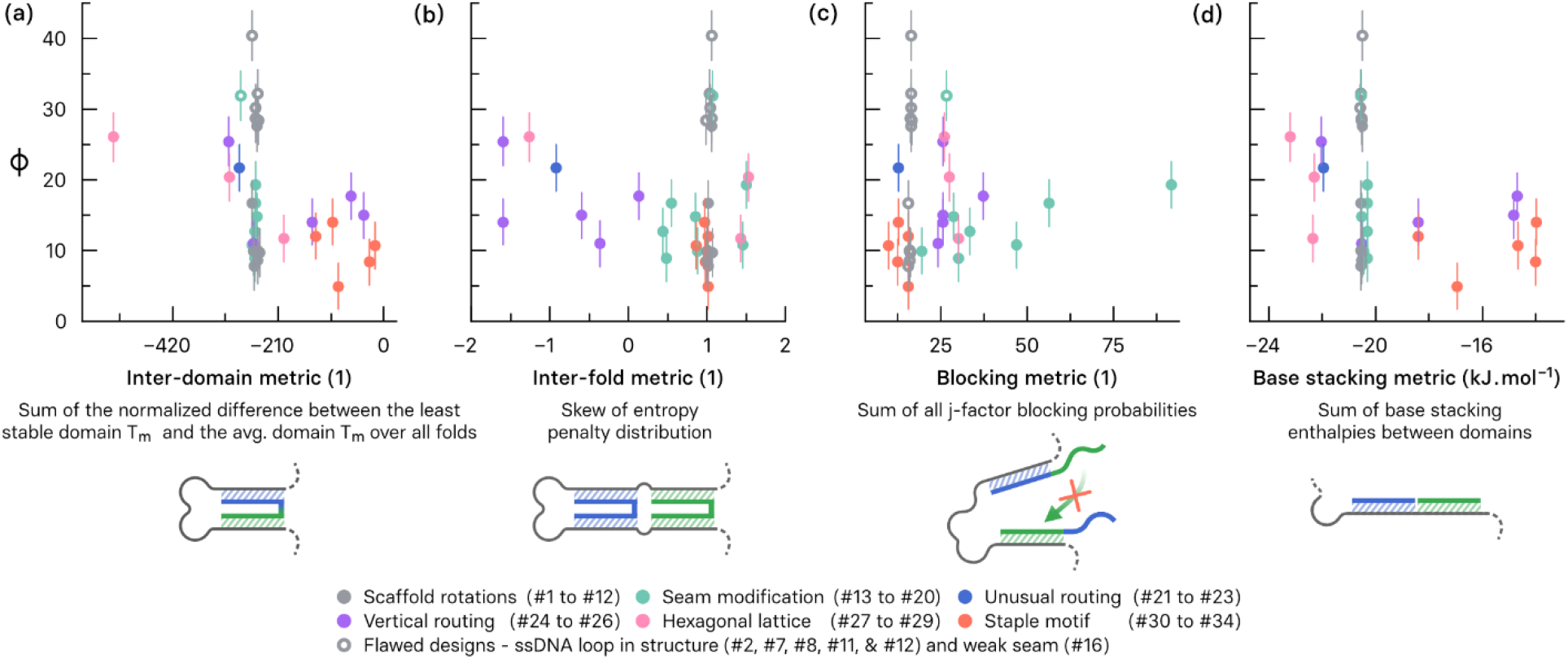
Defectivity φ as a function of several proposed design metrics. Datapoints are colored to correspond with the categories in Fig. 3, where hollow circles represent clearly flawed designs. (1) indicates a unitless quantity, error bars represent a single standard deviation and were calculated via Monte Carlo simulations as discussed in the SI.

The inter-fold cooperativity metric, as measured by the skew of the initial entropy penalty distribution, is strongly correlated with defectivity. We note that the strength of the correlation with negative inter-domain cooperativity (or blocking), as measured by the sum of the blocking probabilities across all folds, is more pronounced for some design families, particularly those with modified staple seams.

While it is intuitive that more blocking leads to higher defectivity, understanding the effect of the entropy penalty skew is more involved. The nucleation and growth picture of origami assembly is intrinsically linked to interfold cooperativity and loop entropy. It is computationally challenging to model the ensemble of different folding pathways for even a single design. However, it is possible to generate the entropy penalty for each fold if it is considered to be the first fold, generating a distribution across all folds. We may also expect the folds with the smallest penalty in this distribution to actually form first, nucleating assembly pathways. While, in principle, it would be possible to recalculate the distribution for the remaining folds, this may not be necessary. This is because, while the initial fold will reduce the penalty for subsequent events, it will not, on average, change the order in which subsequent folds occur. The shape of the distribution, represented by the skew, thus reveals the proportion of nucleation events that can propagate smoothly, i.e., with the smallest discontinuity in penalty, via intermediate to long folds and complete assembly, see SI sections 9 & 10 for histogram distributions of these penalties.

A large negative skew is indicative of many nucleation points, but a limited number of pathways leading to the geometrically necessary long folds. Conversely, a large positive skew represents few nucleation events and an excess of long folds, susceptible to blocking. While it might therefore be expected that a skew of zero would be ideal, the lowest defectivity occurs for a value of +1, which is consistent with the fact that geometrical constraints lead to an irreducible number of longer folds.

Because assembly occurs across a steep thermal gradient, it is useful to have an orthogonal, thermal measurement of these systems. This is also useful as thermal stability is often discussed in the context of assembly, yield, and defectivity^5,15,43,44^

Obtaining high-quality melt data is problematic as the large molar mass of origami and their excess staple populations make calorimetry and temperature-dependent UV-vis absorption measurements difficult, while fluorescence measurements using intercalating dyes are confounded by dye binding to ssDNA and dsDNA, and dye quantum yield, all of which change with temperature^45^. However, intercalating dye measurements are useful as a comparative metric of the assembly process, though we needed to perform the measurements at a higher concentration to ensure high signal, see the methods.

Fig. 6a shows two examples of melt and anneal curves which provide examples of the best and worst performing designs, #7 and #1 respectively. We plot these curves for each designs in section 17 of the SI. In Fig. 6b we plot the anneal and melt peak positions against defectivity. Note that we consider the flawed designs, indicated by hollow circles, as outliers that do not contribute to the trend of reduced defect rate as a function of increased thermal stability.

**Fig. 6.**
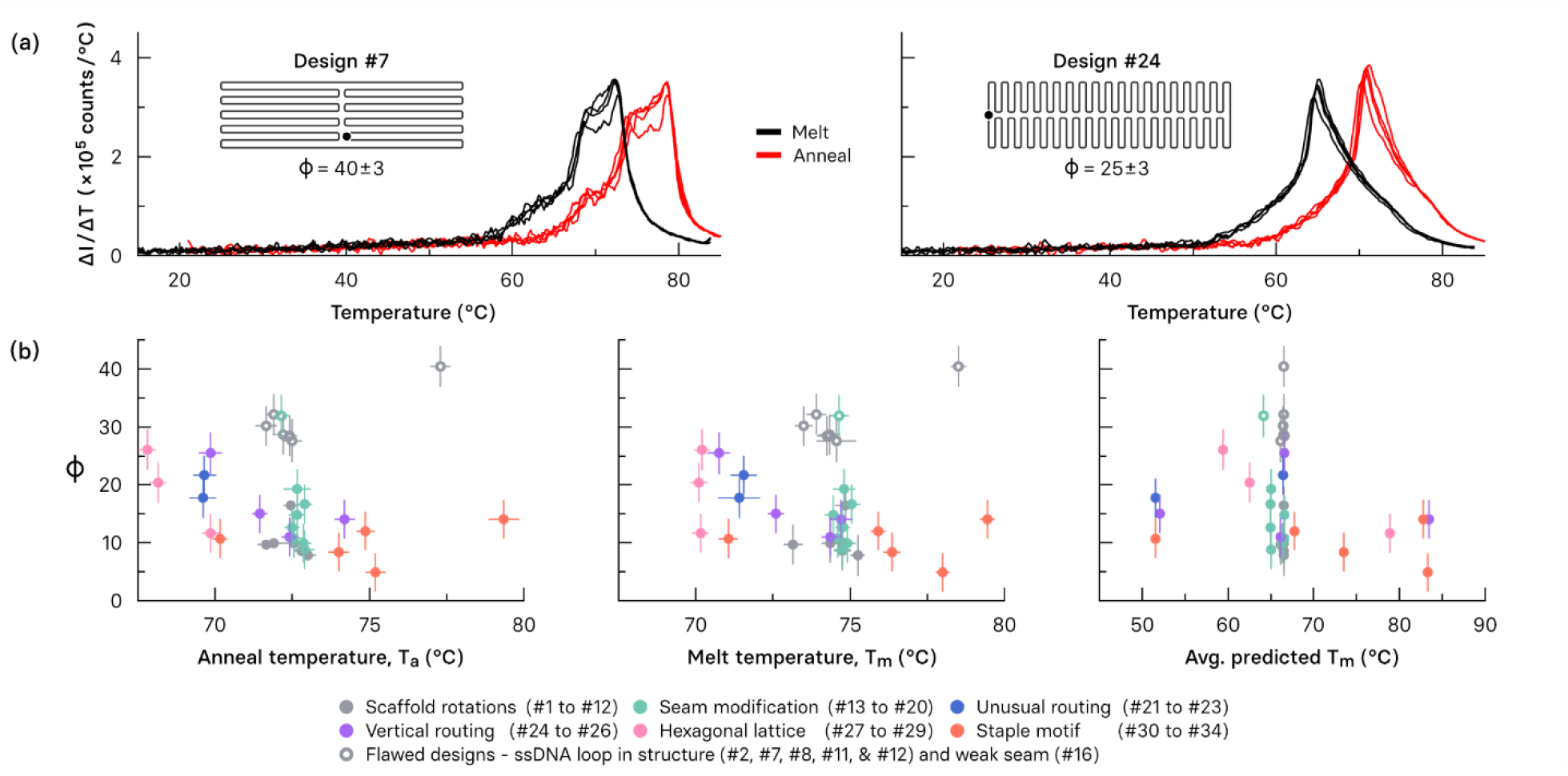
a) numerical derivatives of the melt (red) and anneal (black) curves for four replicate wells for two designs (#7 & #24). Each line represents a single measurement. The black circle indicates the loop of excess ssDNA and the start of the m13 sequence. b) plot of the defectivity, or φ, against the measured anneal temperature, melt temperature, and average predicted full staple melt temperature for each design. We note the similar distribution of different families of design in melt, anneal, and predicted melt temperature distributions. The measured melt and anneal temperatures were found by the derivative peak position. Datapoints are colored to correspond with the categories in Fig. 3, where hollow symbols represent clearly flawed designs. Y-error bars were calculated for φ as described in the text, and x-error bars represent the standard deviation in peak position between the four replicates for experimental T_m_ and T_a_.

The average predicted T_m_ for full staples, including loop entropy penalties is given in the rightmost panel of Fig. 6b. The predicted T_m_ were calculated for all staples in a design, assuming that each staple was the first to bind, and did so in one step, see SI section 2 and 12. This results in an underestimate of the stability of the origami, and a correspondingly lower predicted T_m_.

The observed relationship between stability and defectivity suggests interesting avenues for future work. Improved thermal stability prediction could be invaluable in optimizing designs for yield: Dannenberg *et al*. reported a proof-of-concept model for origami melting which incorporated conformational effects, but which was only experimentally verified for one design and required manual input of topological information^5^. Further testing of this model against multiple designs or annealing conditions could help to deconvolve cooperative contributions.

So far, we have identified and devised cooperativity metrics and shown that a subset correlate strongly with defectivity. Given the highly interconnected nature of cooperative effects in origami assembly, it is not obvious that even the strongly correlated metrics would form a suitably orthogonal basis, *i*.*e*., a set of independent parameters that can be calculated from an origami design, whose combination can effectively predict defectivity. If this is the case, then it implies that it is not necessary to simulate the details of all possible assembly pathways, because the behavior of the system can be understood from the aggregate properties of all possible first folds.

We test for this by combining our metrics, and alternate versions thereof, and determining if that combination collapses the data onto a single curve. Applying Occam’s razor, we fit the data with first order, linear combinations of normalized metrics, and endeavor to find the optimum combination and number of metrics that represent defectivity while not fitting noise. Normalizing, or regularizing, each metric across the distribution of values across all designs from 0 to 1 simplifies fitting and interpretation.

We took two approaches for testing combinations of metrics. The first was to simply combine the metrics with the strongest correlation with defectivity, namely the inter-fold cooperativity and blocking inter-domain cooperativity, with the experimental thermal stability. We then used a linear regression to weight their effect on defectivity. As steric contributions are unlikely to be captured by any of our metrics, the outlier flawed designs were likely to only map noise onto our model and were thus omitted from regression. As shown in Fig. 7 this model fit captured a surprising amount of variation in defectivity. This approach yielded roughly equivalent weights of 33 %, 32 %, and 35 %, for T_m_, entropy skew, and blocking, respectively where T_m_ and entropy skew had a negative sign as their metric is correlated with reduced defectivity.

**Fig. 7.**
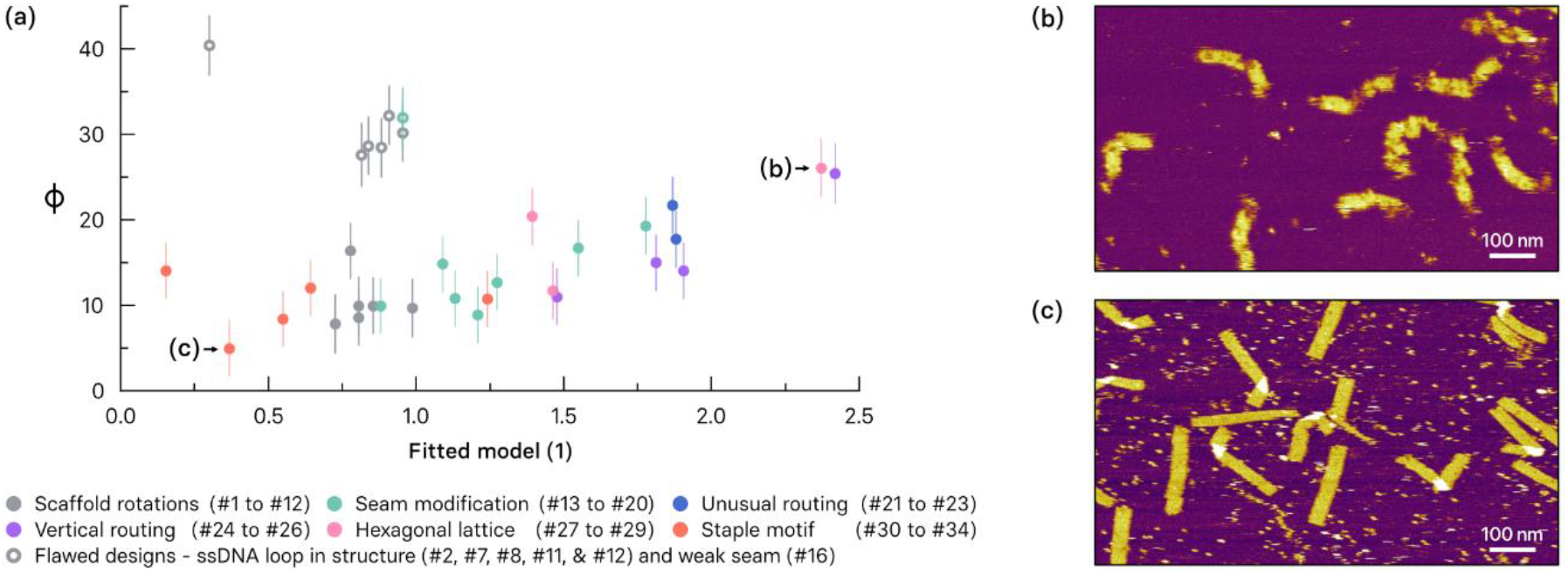
Combined semi-empirical fitted model combining normalized metrics for inter-fold cooperativity and negative inter-domain cooperativity (blocking) with thermal stability in the form of experimental T_m_, with example AFM images of a high defectivity (#28) and low defectivity (#33) designs. The scale bars represent 100 nm. X-error bars in the heuristic come exclusively from the uncertainty in measured melt temperature and fit underneath the data points.

Our second approach was to apply a basic Least Absolute Shrinkage and Selection Operator (LASSO) analysis. A full description of the analysis is provided in section 3 of the SI. In brief, LASSO analysis identifies which subset of input covariates, i.e., metrics, best correlate with our output, the defectivity. Critically, LASSO analysis uses a regularization parameter that can be used to adjust the balance between fit quality and number of fit parameters to avoid overfitting. It is obviously desirable to minimize the number of parameters to avoid fitting noise.

While we have so far presented a single metric for each source of cooperativity. However, it is not obvious, *a priori*, how to best construct a metric that effectively represents cooperativity in such a complex system. We therefore constructed multiple alternate versions (detailed in the SI). This results in an exponential increase in the number of possible combinations of metrics. The regularization feature of the LASSO analysis allows us to detect which metric combinations best capture defectivity.

The LASSO analysis consistently identified inter-fold cooperativity, as described by the skew of the entropy penalty distribution, and thermal stability, as represented by the inverse of the T_m_, as principal factors with similar weighting as our initial analysis. However, it consistently identified the positive inter-domain cooperativity (nucleating staple domains) instead of the negative inter-domain cooperativity, (blocking) and had a variety of other metrics contributing less than 10 % of coefficient weight. Given the variation in contributions between iterations of the LASSO analysis, it is difficult to say whether these metrics are meaningful, or simply a mapping of noise onto the model form.

In summary, it is both surprising and encouraging that simple linear combinations of cooperativity metrics and thermal stability metrics can so readily capture variability in defectivity, at least over the range our design variations can probe. That such a network of hundreds of interacting events can be so easily simplified supports the hypothesis that 2D origami is more funnel-like than path-like. This indicates that over the next decade a combination of physical insight and statistical analysis may, with achievable increases in dataset size and precision, build a clear understanding of how cooperativity propagates through these complex and interconnected model systems.

Additionally, our observations contribute to origami design best practice: absent a predictive model, keeping staples short, maintaining symmetry where possible, and keeping the excess scaffold ssDNA outside the structure on a helix rather than on a scaffold crossover are, collectively, reasonable approaches to good origami design.

## Conclusions

In the DNA nanofabrication literature, the majority of design and cooperativity studies sample small families of related structures and vary a single design parameter, such as staple motif, and have proven useful in identifying sources of cooperativity. Generalizing any observed trends is however problematic, given the complex relationships between design parameters and the various cooperative effects.

By measuring defectivity values precisely, and across a wide range of structures, we have found good correlation between defectivity, and the sources of cooperativity proposed in the literature. An initial statistical analysis indicates that a unified heuristic, created from a linear combination of these cooperativity metrics, once normalized, is surprisingly effective at capturing the trend in defectivity. This suggests that despite, or perhaps because of, the very large number of events and possible assembly pathways, the aggregate properties of event distributions in origami matter more than their precise propagation during assembly. Further, our analysis indicates that the inter-fold cooperativity and thermal stability, are the principal factors controlling defectivity. Interestingly, it appears that the human aesthetic bias towards symmetry reduces defectivity via its effect on the distribution of entropic penalties. The remainder of the variation in defectivity can be ascribed to either positive or negative inter-domain cooperativity, with a potential admixture of base stacking. However, a more extensive data set would be needed to distinguish these contributions.

Fortunately, emerging capabilities in automated design, synthesis, image classification, and even autonomous experimentation should enable the generation of sufficiently large data sets to construct a physics-based predictive model connecting defectivity and design in DNA nanostructures. More importantly, this simple, yet rich system enables a deeper exploration of cooperative effects. If the resulting insights could be applied to the interaction of nanostructure molecular motions, then it would be possible to emulate cellular mechanisms, such as the ribosome.

## Materials and Methods

### Origami anneals

Samples of each design were annealed at a low M13MP18 scaffold concentration of 0.375 nmol.L^-1^, a low staple excess of 5 ×, for a 45 minute period, during which the temperature was reduced from 80 °C to 4 °C. Buffer conditions were 40 mmol.L^-1^ Tris, 12.5 mmol.L^-1^ magnesium chloride, and 1 mmol.L^-1^ Ethylenediaminetetraacetic acid (EDTA), pH 8.0. Samples were then kept on ice for no more than 14 hours before imaging. As discussed above, these poor anneal conditions were chosen to ensure a higher distribution of defective origami to minimize the statistical sample size and AFM imaging time.

### Staple pool concentration

When staple concentrations and excesses are described in this manuscript, we refer to an estimated concentration of correctly synthesized staples. Such normalization is necessary as designs with longer staples will have lower concentrations of correctly synthesized staples^46^. As discussed in section 5 of the SI, we used Monte Carlo simulation of the staple synthesis yield assuming a 99.5 % coupling efficiency and no sequence dependence on coupling efficiency, along with standard extinction coefficient prediction^47,48^. Staple pools were purchased in multiple iterations, discussed in image classification below, though we do not anticipate staple pool age to be relevant to yield^49^.

### Intercalating dye melt and anneal curves

To ensure high signal, we performed these experiments at a much higher concentration and staple excess than in our accelerated test conditions (7.5 nmol.L^-1^ scaffold, 10 × staple concentration), with an intercalating dye (13.3 µmol.L^-1^ SYBR green – Note: Materials are identified in this paper in order to specify the experimental procedure adequately. Such identification is not intended to imply recommendation or endorsement of any product or service by NIST, nor is it intended to imply that the materials or equipment identified are necessarily the best available for the purpose.), and curves were obtained across four replicate wells. For more detailed experimental conditions see section 13 of the SI.

### Gel electrophoresis

1 % agarose gels in annealing buffer were run at 100 V for 2 h on ice, stained with SYBR green, and imaged.

### Atomic Force Microscopy

All images were taken on mica surfaces in liquid tapping mode. Buffer conditions were identical to those used in annealing. If significant precipitation was observed during imaging, imaging was continued so long as the structures could be clearly visualized, and buffer was filtered or prepared fresh for the next sample.

### Image Classification

The first round of classification (designs #2, #5 to #13, #15, and #22) were performed with four analysts. The images were, via script, shuffled and renamed alphanumerically. After classification, the data was unshuffled via key. After confirming trend consistency between analysts, the remainder of the designs were classified by a single analyst. These designs were renamed and were classified in a random order.

### Evaluation of design files

*T*o deconvolve the sources of cooperativity from defectivity data, it was necessary to condense design information into our devised metrics. As discussed in section 11 of the SI, we modified a version of cadnano to provide outputs including domain sequences, loop distances between domains, and basic nearest neighbor model thermodynamic predictions.

Section 4 of the SI discusses the organization of the supplemental zip file containing sequence lists for each design, broken up into folds, or sets of two domains on the same staple, with the predicted T_m_ for each domain, the loop distance, and estimated entropy penalty.

## Acknowledgments

We would like to thank John Majikes for help in troubleshooting and streamlining our analysis scripts. We would like to thank Sam Stavis, Andy Madison, Ruohong Shi, Sam Schaffter, and Arvind Balijepalli for useful discussions. We would also like to thank Sam Schaffter, Divita Mathur, and Dominic Scalise for their feedback on our methods and manuscript.

## Funding and Disclaimers

This work was performed under financial assistance award 70NANB21H182 from the National Institute of Standards and Technology, U.S. Department of Commerce. The statements, findings, conclusions, and recommendations are those of the author(s) and do not necessarily reflect the views of the National Institute of Standards and Technology or the U.S. Department of Commerce.

## Bibliography

1. Hartman, N. C. & Groves, J. T. Signaling clusters in the cell membrane. Current Opinion in Cell Biology vol. 23 Preprint at 10.1016/j.ceb.2011.05.003 (2011).

2. Levinthal, C. Are there pathways for protein folding? Journal de Chimie Physique 65, (1968).

3. Hill, A. V. The possible effects of the aggregation of the molecules of haemoglobin on its dissociation curves. Proceedings of the Physiological Society vol. 40 Preprint at (1910).

4. DeLuca, M. et al. Mechanism of DNA origami folding elucidated by mesoscopic simulations. Nat Commun 15, (2024).

5. Dannenberg, F., Dunn, K. E., Bath, J., Turberfield, A. J. & Ouldridge, T. E. Modelling DNA Origami Self-Assembly at the Domain Level. J Chem Phys 143, 165102 (2015).

6. Aksel, T., Navarro, E. J., Fong, N. & Douglas, S. M. Design principles for accurate folding of DNA origami. bioRxiv (2024).

7. Jun, H. et al. Autonomously designed free-form 2D DNA origami. Sci Adv 5, 1–9 (2019).

8. Babatunde, B., Cagan, J. & Taylor, R. E. An Improved Shape Annealing Algorithm for the Generation of Coated Deoxyribonucleic Acid Origami Nanostructures. Journal of Mechanical Design 146, (2024).

9. Shirt-Ediss +, B. et al. Optimizing DNA origami assembly through selection of scaffold sequences that minimise off-target interactions. bioRxiv 2025.01.29.635450 (2025) doi:10.1101/2025.01.29.635450.

10. Dunn, K. E. The business of DNA nanotechnology: Commercialization of origami and other technologies. Molecules 25, (2020).

11. Mathur, D. & Medintz, I. L. Analyzing DNA Nanotechnology: A Call to Arms For The Analytical Chemistry Community. Anal Chem acs.analchem.6b04033 (2017) doi:10.1021/acs.analchem.6b04033.

12. Kick, B., Praetorius, F., Dietz, H. & Weuster-Botz, D. Efficient Production of Single-Stranded Phage DNA as Scaffolds for DNA Origami. Nano Lett 15, 4672–4676 (2015).

13. Majikes, J. M. & Liddle, J. A. Synthesizing the biochemical and semiconductor worlds: the future of nucleic acid nanotechnology. Nanoscale 14, 15586–15595 (2022).

14. Ke, Y., Bellot, G., Voigt, N. V., Fradkov, E. & Shih, W. M. Two design strategies for enhancement of multilayer–DNA-origami folding: underwinding for specific intercalator rescue and staple-break positioning. Chem Sci 3, 2587 (2012).

15. Martin, T. G. & Dietz, H. Magnesium-free self-assembly of multi-layer DNA objects. Nat Commun 3, 1103–1106 (2012).

16. Jacobs, W. M., Reinhardt, A. & Frenkel, D. Rational design of self-assembly pathways for complex multicomponent structures. Proc Natl Acad Sci U S A 112, 6313–8 (2015).

17. Jacobs, W. M. & Frenkel, D. Self-Assembly of Structures with Addressable Complexity. J Am Chem Soc 138, 2457–2467 (2016).

18. Majikes, J. M., Nash, J. A. & LaBean, T. H. Search for effective chemical quenching to arrest molecular assembly and directly monitor DNA nanostructure formation. Nanoscale 9, 1637–1644 (2017).

19. Wah, J. L. T., David, C., Rudiuk, S., Baigl, D. & Estevez-Torres, A. Observing and Controlling the Folding Pathway of DNA Origami at the Nanoscale. ACS Nano 10, 1978–1987 (2016).

20. Dunn, K. E. et al. Guiding the folding pathway of DNA origami. Nature (2015) doi:10.1038/nature14860.

21. Song, J. et al. Direct visualization of transient thermal response of a DNA origami. J Am Chem Soc 134, 9844–7 (2012).

22. Arbona, J.-M., Aimé, J.-P. & Elezgaray, J. Cooperativity in the annealing of DNA origamis. J Chem Phys 138, 015105 (2013).

23. Snodin, B. E. K. et al. Direct Simulation of the Self-Assembly of a Small DNA Origami. ACS Nano 10, 1724–1737 (2016).

24. Majikes, J., Patrone, P., Kearsley, A., Zwolak, M. & Liddle, J. Failure Mechanisms in DNA Self-Assembly: Barriers to Single-Fold Yield. ACS Nano 15, 3284–3294.

25. Cumberworth, A., Frenkel, D. & Reinhardt, A. Simulations of DNA-Origami Self-Assembly Reveal Design-Dependent Nucleation Barriers. Nano Lett 22, (2022).

26. Rossi-Gendron, C. et al. Isothermal self-assembly of multicomponent and evolutive DNA nanostructures. ChemRXiv (2022).

27. Majikes, J. M., Nash, J. A. & LaBean, T. H. Competitive annealing of multiple DNA origami: formation of chimeric origami. New J Phys 18, 115001 (2016).

28. Schneider, F., Möritz, N. & Dietz, H. The sequence of events during folding of a DNA origami. Sci Adv 5, (2019).

29. Majikes, J. M. & Liddle, J. A. DNA Origami Design: A How-To Tutorial. J Res Natl Inst Stand Technol 126, (2021).

30. Wagenbauer, K. F. et al. How We Make DNA Origami. ChemBioChem 18, 1873–1885 (2017).

31. Halvorsen, K., Kizer, M. E., Wang, X., Chandrasekaran, A. R. & Basanta-Sanchez, M. Shear Dependent LC Purification of an Engineered DNA Nanoswitch and Implications for DNA Origami. Anal Chem acs.analchem.7b00791 (2017) doi:10.1021/acs.analchem.7b00791.

32. Martin, T. G. & Dietz, H. Magnesium-free self-assembly of multi-layer DNA objects. Nat Commun 3, 1103–1106 (2012).

33. Ko, S. H., Gallatin, G. M. & Liddle, J. A. Nanomanufacturing with DNA origami: Factors affecting the kinetics and yield of quantum dot binding. Adv Funct Mater 22, 1015–1023 (2012).

34. Sheheade, B. et al. Self-Assembly of DNA Origami Heterodimers in High Yields and Analysis of the Involved Mechanisms. Small 1902979, 1902979 (2019).

35. Ko, S. H., Gallatin, G. M. & Liddle, J. A. Nanomanufacturing with DNA origami: Factors affecting the kinetics and yield of quantum dot binding. Adv Funct Mater 22, 1015–1023 (2012).

36. Green, C. M., Hughes, W. L., Graugnard, E. & Kuang, W. Correlative Super-Resolution and Atomic Force Microscopy of DNA Nanostructures and Characterization of Addressable Site Defects. ACS Nano 15, 11597–11606 (2021).

37. Strauss, M. T., Schueder, F., Haas, D., Nickels, P. C. & Jungmann, R. Quantifying absolute addressability in DNA origami with molecular resolution. Nat Commun 9, 1–7 (2018).

38. Wagenbauer, K. F., Wachauf, C. H. & Dietz, H. Quantifying quality in DNA self-assembly. Nat Commun 5, 3691 (2014).

39. Berengut, J. F., Berg, W. R., Rizzuto, F. J. & Lee, L. K. Passivating Blunt-Ended Helices to Control Monodispersity and Multi-Subunit Assembly of DNA Origami Structures. Small Struct 5, (2024).

40. Fuentes-Arderiu, X. & Dot-Bach, D. Measurement uncertainty in manual differential leukocyte counting. Clin Chem Lab Med 47, 112–115 (2009).

41. Sobczak, J.-P. J., Martin, T. G., Gerling, T. & Dietz, H. Rapid folding of DNA into nanoscale shapes at constant temperature. Science 338, 1458–61 (2012).

42. Hernandez-Garcia, A. et al. Precise Coating of a Wide Range of DNA Templates by a Protein Polymer with a DNA Binding Domain. ACS Nano 11, 144–152 (2017).

43. Arbona, J. M., Elezgaray, J. & Aimé, J. P. Modelling the folding of DNA origami. EPL (Europhysics Letters) 100, 28006 (2012).

44. Harkness V R. W.; Avakyan, N., Sleiman, H. F. & Mittermaier, A. K. Mapping the energy landscapes of supramolecular assembly by thermal hysteresis. Nat Commun 9, (2018).

45. DeJaco, R. F., Majikes, J. M., Liddle, J. A. & Kearsley, A. J. Binding, brightness, or noise? Extracting temperature-dependent properties of dye bound to DNA. Biophys J 122, (2023).

46. Sabary, O. et al. SOLQC: Synthetic oligo library quality control tool. Bioinformatics 37, (2021).

47. Cavaluzzi, M. J. & Borer, P. N. Revised UV extinction coefficients for nucleoside-5’-monophosphates and unpaired DNA and RNA. Nucleic Acids Res 32, (2004).

48. Cantor, C. R., Warshaw, M. M. & Shapiro, H. Oligonucleotide interactions. III. Circular dichroism studies of the conformation of deoxyoligonucleolides. Biopolymers 9, 1059–1077 (1970).

49. Kielar, C. et al. Effect of Staple Age on DNA Origami Nanostructure Assembly and Stability. Molecules 24, (2019).

