## Supplemental Data for "Complex cooperativity in DNA origami revealed *via* design dependent defectivity"

#### Contents

Section 1: Cooperativity metrics across the hierarchy of folding events ... 2
Section 2: Metric calculations ... 3
Section 3: LASSO analysis ... 6
Section 4: Organization of design information ... 12
Section 5: Staple pool concentration normalization ... 14
Section 6: Imaging yield procedure and classification ... 14
Section 7: Raw Experimental Counts ... 15
Section 8: Uncertainty in imaging yield ... 18
Section 9: Qualitative Trends ... 24
Section 10: Seam symmetry designs ... 25
Section 11: cadnano modification for topological information ... 28
Section 12: Nearest neighbor calculations for predicted $T_m$ ... 29
Section 13: Intercalating dye melt data ... 29
Section 14: Comparison to literature metrics ... 31
Section 16: Supplementary bibliography ... 32
Section 17: Design information sheets ... 33

#### Section 1: Cooperativity metrics across the hierarchy of folding events

One persistent note of concern we raise in this manuscript is the risk of developing too many different model metrics for cooperativity – as there is no real difference between continuously making new models until finding one that fits well and fitting a polynomial of arbitrary order to the data. In both cases one will eventually fit the data perfectly well, but neither is likely to be predictive or representative of physical reality.

Any metric for an individual event must be considered across all such events for the entire origami. Cooperativity is most clearly defined as the excess energy consumed or evolved when one of two otherwise identical events or reactions occurs before the other. Origami are comprised of hundreds of hybridization events, some of which are highly independent and some of which are highly cooperative. Defining a metric which quantifies cooperativity across such a network of events is a much less well-defined problem than between a handful of events.

Additionally, there are many conceptual levels on which to view these events (single domain, single fold, full staple, *etc*), and many ways to consider the distribution of events (sum, average, mode, skew, *etc*). These distributions could also be taken across individual folding paths separately or all events together. In short, even if a source of cooperativity is well-defined between two events, one could easily create multiple, equally defensible equations that result in different final values.

The simplest possible event lowest on this hierarchy is the hybridization of a single domain. This is reasonably well modeled by the nearest neighbor model.

Above this is folding hybridization events. These occur when two domains that are shared by the same staple bind, simultaneously or in sequence. This incurs some conformational entropy penalty that is dependent on which other folds have already occurred. Unfortunately, when staples have more than 2 domains, averaging across folds or taking some moment of the distribution of properties of the folds will, by definition, double count the center domains.

At the next higher step of complexity is full staple hybridization. In this case one considers the binding of all domains on the staple simultaneously. While such events are improbable, treating cooperativity across whole staples has the benefit of including all the relevant domains and entropic penalties without double counting domains that neighbor multiple folds.

At the next higher step of complexity is what we term folding 'fingers' or pathways. These fingers are the large loops in the scaffold which approach the origami seam and the origami edge. The fingers of a scaffold inherently determine which staples are most cooperative with each other and seem associated with nucleation and growth behavior that has been observed[1,2]. Unfortunately, calculating some cooperativity metric across the folding fingers requires sufficiently many model function decisions that testing all of them would quickly create an even more overdetermined system.

As fully simulating these systems is infeasible, it is also tempting to weight the cooperative metrics by, for example, the $T_m$ of the domain/fold/staple relative to the average $T_m$ in the origami. Such a weighting could account for annealing across a thermal gradient as the more highly stable events would occur first and have a disproportionate effect.

In summary, one could make a nearly infinite number of metrics for a single source of cooperativity by considering that source at different levels of conceptual abstraction (domain, fold, staple, etc.), by taking different statistical moments of the distribution of events (average, mode, skew, etc.), and by weighting the events in the distribution by other relevant properties (thermal stability, order in a folding finger, *etc*).

#### Section 2: Metric calculations

Here we briefly discuss the various formulations we devised to quantify the description of individual sources of cooperativity not for a single staple, but for the whole origami design. We evaluated some other formulations and weighting schemes which we discarded, most of which involving weighting across folding pathways or fingers. The list below is organized by groups of related metrics, and is representative of what we input into the LASSO algorithm.

*Table 1: Names and definitions for various formulations of cooperativity metrics.*

| | **Name** | **Description** |
|---|---|---|
| **Experimental** | Exp.AnnealTemp | The experimental $T_a$, as determined by the peak in the anneal curve, averaged across four replicates and acquired using an intercalating dye |
| | Exp.MeltTemp | The experimental $T_m$, as determined by the peak in the anneal curve, averaged across four replicates and acquired using an intercalating dye |
| | hystArea | The area between the melt and anneal curves found by integrating between them, with a cutoff for the low temperature at which their difference is only noise. |
| **Thermal Stability** | averageFullStapleTm | The nearest neighbor (NN) predicted $T_m$ of all domains on the staple, accounting for loop entropy contributions of that staple being the first to bind. This value is then averaged across all staples in a design |
| | stdFullStapleTm | The standard deviation of the previous list of $T_m$ values. |
| | averageDomainTm | The average of NN predicted $T_m$ of all domains on all staples, with no loop entropy or base stacking contributions. |
| | stdDomainTm | The standard deviation of the previous list of $T_m$ values. |
| | foldAverageTm | The average of the $T_m$ for all folds across all folds in all staples. A fold is any set of two connected staple domains. Their $T_m$s are calculated as though those two domains bound simultaneously with a loop entropy contribution calculated as though that fold was first to bind to the scaffold. NOTE: when there are more than 2 domains, this effectively double counts domains in the middle of a staple. |
| | foldStdTm | The standard deviation of the previous list of $T_m$ values. |

| | | |
|---|---|---|
| | avgFullStaplePlain | The $T_m$ of all domains on the staple, as though all events occurred simultaneously, including dsDNA nucleation penalties for each domain, but neglecting entropy penalties. |
| | stdFullStaplePlain | The standard deviation of the previous list of $T_m$ values. |
| | - | |
| Positive Inter-Domain | Avg.DomainStdbyStaple | The average, across all staples, of the standard deviation between domain $T_m$s for that staple |
| | average( intraDomainDiff_normalized) | The average, across all staples, of the per-staple average difference between $T_m$s for each pair of domains in each fold in those staples normalized by the all domain average $T_m$ |
| | average(intraDomainSum) | The average, across all staples, of the sum of the average differences between $T_m$s for each pair of domains in each fold in those staples |
| | average(intraDomainStd) | The average, across all staples, of the standard deviation of the differences in Tm values for all staples that have more than two folds |
| | average(intraDomainStdWt) | The average, across all staples, of the differences in $T_m$ values for all staples that have more than two folds where each deviation is weighted by the Tm of the most stable domain normalized by the origami average $T_m$ |
| | sum(domainWeights) | For each fold, take the difference between the lowest $T_m$ domain and that of the domain average $T_m$ |
| Base stacking | sumDomainBaseStack | The sum of the enthalpy term of the nearest neighbor model for the DNA bases between staple enforced crossovers |
| | avgDomainBaseStack | The per-staple average of the above |
| | wtAvgDomainBaseStack | The per-staple average of the above weighted by the domain $T_m$ relative to the average domain $T_m$ |
| | - | |
| Positive Inter-Fold | EntropySkew | The skew in the distribution of loop entropy penalties incurred if each fold were the first to form |
| | LoopSkew | The skew in the distribution of looping distances folded if each fold were the first to form |
| | sSkewTmWeighted | The skew in the distribution of entropies, as weighted by the difference between the lowest domain Tm in the fold and all domain average |
| | sSkewDomainWeighted | The skew in the distribution of entropies, as weighted by the difference between the full fold Tm (w/ penalties) and the lowest domain Tm in the fold |
| | - | |

| | | |
|---|---|---|
| **Blocking Neg. Inter-Domain** | sum(blockList) | The sum, across all folds, of the blocking probabilities (described below) |
| | average(blockList) | The average, across all folds, of the blocking probabilities |
| **Misc. information** | skew(foldWeights) | The skew of the distribution for each fold of weighting values for the fold $T_m$s normalized by the origami's average fold $T_m$ |
| | skew(domainWeights) | The skew of the distribution for each fold of weighting values for that fold's lowest domain $T_m$ normalized by the origami's average domain $T_m$ |
| | sum(foldWeights) | For each fold, take the difference between the fold's $T_m$ and the average of all fold $T_m$s |
| | crossoverBaseStacking | The sum of the enthalpy term of the nearest neighbor model for the DNA bases between staple enforced crossovers, whether that is a single or double crossover |
| | avgCrossoverBase | The average enthalpy between bases engaged in single crossovers, averaged across all single crossovers |
| | numSingleCross | The total number of single crossovers whether they're engaged in a double crossover or are independent |
| | numDomains | The total number of domains |

For skewness we used the Fisher-Pearson coefficient of skewness as calculated by the SciPy toolkit[3].

*Loop entropy (for fold and staple Tm and entropy distribution skew):*

We calculate our loop entropy penalty as in previous work[4], given in Eq. 1 below. where, $\Delta S_{ref}$ is a reference value for a 30 nucleotide ssDNA loop, R is the gas constant, and $L_0$, $L_1$, & $L_2$ are the initial loop length and the lengths of the smaller loops (in nucleotides) it is folded into, respectively. The loop lengths are combined in this way because the ssDNA is not transitioning from a line to a loop, but from a loop to a figure-eight.

*Eq. 1* $$\Delta S_{loop} = \Delta S_{ref} + 2.44R\ ln\left(\frac{L_1 L_2}{30 L_0}\right) where\ j_{\text{fold}} = e^{\left(-\Delta G_{loop}/_{RT}\right)} = e^{\left(\Delta S_{loop}/_{R}\right)}$$

*Negative inter-domain cooperativity:*

Our negative inter-domain cooperativity (blocking) metric is based on of our previous work[5], though we include a short description here. For this metric, we first consider a single fold, in which staples with two binding domains hybridize with their respective complementary binding domains on the scaffold, where [S] is the staple, [T] is the scaffold or template, [B] is the defective or blocked state, and [F] is the desired folded state.

$$[F] + [S] \rightleftharpoons [\mathrm{B}]$$

Notably, though one would expect this reaction to be multi-step and its kinetics to be quite slow, the number of hybridized base pairs on either side of the reaction are equivalent. As such, the equilibrium populations should be determined only by the concentration of staples, the entropic penalty of the fold, and the entropic penalty of binding a second copy of a staple.

In short, we assume that at low temperatures potential fold positions will be at equilibrium between being folded and having bound two copies of a staple. The anticipated fraction of the blocked state for some fold f, $B_f$, at the end of an anneal is shown in Eq. 2. In which the initial staple concentration, $\mathcal{S}^o$, the initial scaffold concentration, $\mathcal{T}^o$, and the j-factor, $j_{fold}$, (described in Eq. 1) are taken as variables[40]. The j-factor and entropy penalty may also be interpreted as a measure of the local concentration of the two ends of the DNA being forced together. At long loop distances, there is a low local concentration, low j-factor, and large entropy penalty and a higher probability of forming the defective blocked state.

*Eq. 2*
$$B_f = 1 - \left(\frac{1}{2\mathcal{T}^o}\left(-(j_{\text{fold}} + \mathcal{S}^o - 2\mathcal{T}^o) + \sqrt{(j_{fold} + \mathcal{S}^o - 2\mathcal{T}^o)^2 + 4j_{\text{fold}}\mathcal{T}^o}\right)\right)$$

For a metric of negative inter-domain cooperativity across a whole origami design, we use the total anticipated number of blocked staples in an origami, *i.e.*, the sum of the probabilities that an individual fold enters the blocked state over all M folds in a design using the initial entropic penalties shown in Eq 3.

*Eq 3*
$$\#_{staplesBlocked} = \sum_{fold_i}^{M} \sum B_{f\_i}$$

#### Section 3: LASSO analysis

The Least Absolute Shrinkage and Selection Operator (LASSO) or L1 regularized linear regression automatically selects variables by setting some regression coefficients exactly to zero. More generally, it is a method for preventing overfitting in linear regression problems[6]. In situations with many input variables, even more than then number of samples, it can efficiently find those most predictive for the output variable.

The LASSO does this by adding a penalty to the ordinary residual sum of squares objective; the penalty is the sum of the absolute value of the regression coefficients (the intercept is typically omitted). The regularization parameter λ controls the strength of this penalty. When λ is increased, more coefficients for input variables are forced to zero, and the output variable in the training data set will not be predicted as well as by ordinary least squares. When λ is decreased, fewer covariates will be forced to zero, and the resulting model will predict the output variable in the training data more similarly to ordinary least squares, perhaps in a non-physical way if there are many input variables.

In this work we used the glmnet package in R, and ran the LASSO analysis with two sets of input covariates and two selections of the λ parameter[7,8]. Both sets of input covariates included all the covariates listed in Table 1 along with their inverses, normalized from 0 to 1 as described in the main text and used defectivity as the output. The first set of inputs contained only these, while the other also included the experimental $T_m$, and $T_a$ (as defined by the peak in the first derivative of the melt or anneal curve) in addition to the hysteresis area (as defined by the integration of the area between melt and anneal curves, see SI section 13). The first selection of regularization parameter was static and was chosen to mirror our initial fit of the defectivity data (in which we chose entropy skew, experimental $T_m$, and blocking metrics based on our intuition and performed a linear regression to weight their contributions). This approach assigned a penalty equivalent to $\sum_i |\beta_i| = 3$. Our second selection of a regularization parameter was performed through a cross-validation approach in which the samples were partitioned randomly into 10 sets (2 or 3 samples in each set). Each set is left-out once, and the other 9 sets are used to fit the model for many different penalty values. The penalty value leading to the best predictions of the left-out sets is selected. Since the original partition into 10 sets is random the entire cross-validation procedure is performed 10 times to evaluate the effect of the random splitting. Input variables receiving non-zero coefficients across many or all of the random splits are more likely to be important predictors. For all LASSO analyses, the high defectivity designs which were flawed due to steric hindrance were neglected in fitting, as none of our metrics should capture steric effects.

We begin with a brief summary and representative weightings, then present full tables of the fitted coefficients.

The first observation we note is that the LASSO results were reasonably consistent across the iterations where λ was found via cross validation. This was particularly true for those covariates whose relative size were more than 5 % of the total, i.e., $\frac{\beta_i}{\sum_j \beta_j} > 0.05$. The minor instability between random data splits for less heavily weighted covariates could indicate either slight overfitting, or that there are multiple semi-stable sets of covariates.

Table 1 gives a brief summary of the relative sizes of the non-zero fitted coefficients to the overall assigned weights for our four analysis conditions and Table 2, Table 3, and Table 4 provide all the coefficients.

We also note that in most cases the analysis identified inverse $T_m$ or $T_a$ rather than the untransformed value. We believe this to be an artifact of our normalization. The $T_m$ and inverse $T_m$ values are normalized to the interval [0, 1] individually, and as such are almost perfectly linearly correlated except in that the latter slightly compresses the distance between the more thermally stable designs and slightly expands the distance between less stable designs. Despite this, the LASSO analysis places more weight on the experimental $T_m$ rather than the average staple predicted $T_m$.

*Table 2: Summary of weights for different iterations of LASSO analysis. The weights for the cross-validated λ were averaged across all 10 iterations, for those with a greater than 1 % contribution*

| With experimental $T_m$ and $T_a$ | No experimental $T_m$ or $T_a$ |
|---|---|

| | | | | |
|---|---|---|---|---|
| *λ by Cross Validated* | invExp.MeltTemp | 25.0 % | EntropySkew | 38.1 % |
| | EntropySkew | 24.6 % | invaverageFullStapleTm | 22.5 % |
| | sum(domainWeights) | 16.9 % | sum(domainWeights) | 14.1 % |
| | invLoopSkew | 12.3 % | invLoopSkew | 10.2 % |
| | invstdFullStaplePlain | 8.6 % | foldStdTm | 8.4 % |
| | skew(foldWeights) | 6.0 % | invstdFullStaplePlain | 3.9 % |
| | avgDomainBaseStack | 3.9 % | skew(foldWeights) | 1.9 % |
| | stdDomainTm | 1.3 % | LoopSkew | 0.9 % |
| | LoopSkew | 1.0 % | | |
| | invsSkewDomainWeighted | 0.4 % | | |
| $\sum_i \lvert\beta_i\rvert = 3$ | EntropySkew | 26.2 % | EntropySkew | 36.8 % |
| | invExp.MeltTemp | 26.1 % | invaverageFullStapleTm | 22.4 % |
| | sum(domainWeights) | 18.5 % | sum(domainWeights) | 14.4 % |
| | invLoopSkew | 13.6 % | invLoopSkew | 11.6 % |
| | invstdFullStaplePlain | 8.6 % | foldStdTm | 8.6 % |
| | skew(foldWeights) | 5.4 % | invstdFullStaplePlain | 4.0 % |
| | avgDomainBaseStack | 1.3 % | skew(foldWeights) | 2.2 % |
| | stdDomainTm | 0.3 % | | |

We note also that metrics that were weighted less than 20 %, typically had a high relative standard deviation of coefficient weighting between the 10 replicate runs of the LASSO analysis. In particular the *sum(domainWeights)* consistently appear together as a group, but have relatively weak individual weightings. These metrics were weighted noticeably less where the experimental $T_m$ data was included.

In SI *Figure 1* we plot the defectivity against the models provided by the LASSO analyses where $\sum_i |\beta_i|$static at 3, and note they both plot similarly well to the model in which we chose three 3 metrics via physical intuition and weighted their contributions equally. We interpret the consistently high weighting of entropySkew and $T_m$ in Table 2 to support our intuition that these metrics are important for predicting defectivity.

For completeness, the three metrics chosen by intuition were regressed on defectivity using ordinary least squares (OLS), and were found to have relative coefficient sizes as below. The estimated coefficient for the blocking metric (*sum(blockList)*) had the opposite sign as the coefficients for the inter-fold cooperativity metric and the experimentally derived thermal stability.

| | Coeff | % of total |
|---|---|---|
| Intercept | 2.26 | |
| Exp.MeltTemp | -1.22 | 33.7 % |
| EntropySkew | -1.14 | 31.5 % |
| sum(blockList) | 1.25 | 34.8 % |

*Figure 1* plots the fitted model side-by-side with the two LASSO analyses in which $\sum_i |\beta_i|$was held at 3, which is roughly equivalent to $\sum_i |\beta_i|$ for the OLS fitted model. We note that both plots which used the

experimental $T_m$ had an equivalently clear trend and that all three had clear correlations with defectivity.

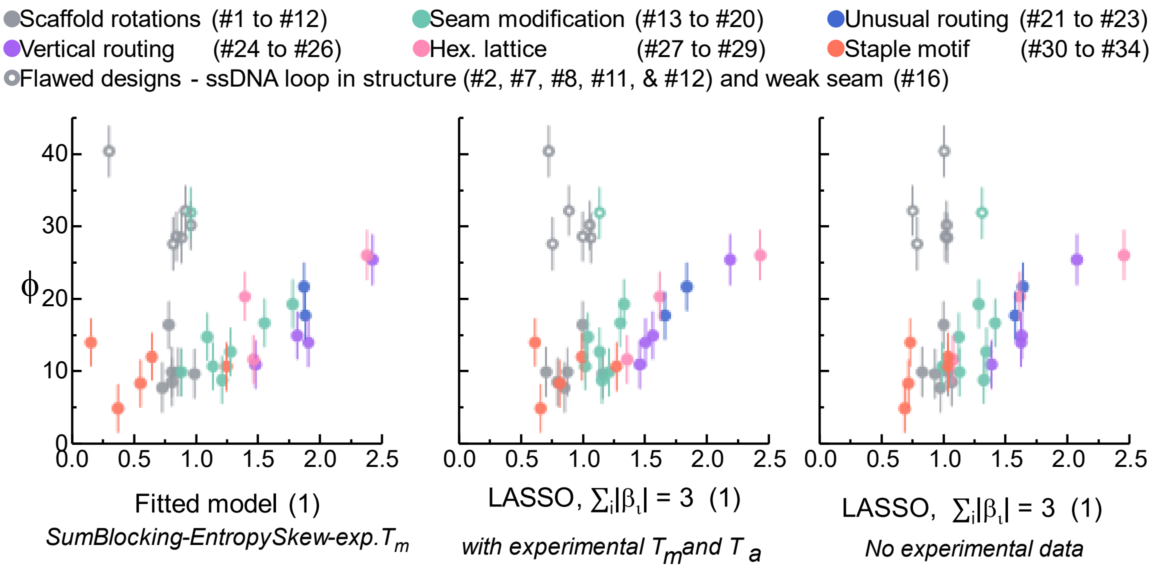

*Figure 1: Measured defectivity plotted against the various fitted models representing cooperativity. Color scheme is as in the main text and error bars are calculated as in the main text and represent a single standard deviation including estimated contributions of staple pool synthesis, and day-to-day variation as well as sample size.*

*Table 3: All Fitted coefficients, λ via cross-validation, Experimental $T_m$ & $T_a$ included*

| | *s1* | *s2* | *s3* | *s4* | *s5* | *s6* | *s7* | *s8* | *s9* | *s10* | *Avg.* | *% of Total* | *Rel. Std.* |
|---|---|---|---|---|---|---|---|---|---|---|---|---|---|
| *(Intercept)* | 2.14 | 1.90 | 1.44 | 2.11 | 1.98 | 2.26 | 1.94 | 2.05 | 1.54 | 2.08 | | | |
| *Exp.AnnealTemp* | 0.00 | 0.00 | 0.00 | 0.00 | 0.00 | 0.00 | 0.00 | 0.00 | 0.00 | 0.00 | 0.00 | 0.0 % | #N/A |
| *Exp.MeltTemp* | 0.00 | 0.00 | 0.00 | 0.00 | 0.00 | 0.00 | 0.00 | 0.00 | 0.00 | 0.00 | 0.00 | 0.0 % | #N/A |
| *hystArea* | 0.00 | 0.00 | 0.00 | 0.00 | 0.00 | 0.00 | 0.00 | 0.00 | 0.00 | 0.00 | 0.00 | 0.0 % | #N/A |
| *averageFullStapleTm* | 0.00 | 0.00 | 0.00 | 0.00 | 0.00 | 0.00 | 0.00 | 0.00 | 0.00 | 0.00 | 0.00 | 0.0 % | #N/A |
| *stdFullStapleTm* | 0.00 | 0.00 | 0.00 | 0.00 | 0.00 | 0.00 | 0.00 | 0.00 | 0.00 | 0.00 | 0.00 | 0.0 % | #N/A |
| *averageDomainTm* | 0.00 | 0.00 | 0.00 | 0.00 | 0.00 | 0.00 | 0.00 | 0.00 | 0.00 | 0.00 | 0.00 | 0.0 % | #N/A |
| *stdDomainTm* | 0.07 | 0.01 | 0.00 | 0.06 | 0.03 | 0.11 | 0.02 | 0.05 | 0.00 | 0.05 | 0.04 | 1.3 % | 88.7% |
| *foldAverageTm* | 0.00 | 0.00 | 0.00 | 0.00 | 0.00 | 0.00 | 0.00 | 0.00 | 0.00 | 0.00 | 0.00 | 0.0 % | #N/A |
| *foldStdTm* | 0.00 | 0.00 | 0.00 | 0.00 | 0.00 | 0.00 | 0.00 | 0.00 | 0.00 | 0.00 | 0.00 | 0.0 % | #N/A |
| *avgFullStaplePlain* | 0.00 | 0.00 | 0.00 | 0.00 | 0.00 | 0.00 | 0.00 | 0.00 | 0.00 | 0.00 | 0.00 | 0.0 % | #N/A |
| *stdFullStaplePlain* | 0.00 | 0.00 | 0.00 | 0.00 | 0.00 | 0.00 | 0.00 | 0.00 | 0.00 | 0.00 | 0.00 | 0.0 % | #N/A |
| *avg.DomainStdbyStaple* | 0.00 | 0.00 | 0.00 | 0.00 | 0.00 | 0.00 | 0.00 | 0.00 | 0.00 | 0.00 | 0.00 | 0.0 % | #N/A |
| *average(intraDomainDiff_normalized)* | 0.00 | 0.00 | 0.00 | 0.00 | 0.00 | 0.00 | 0.00 | 0.00 | 0.00 | 0.00 | 0.00 | 0.0 % | #N/A |
| *average(intraDomainSum)* | 0.00 | 0.00 | 0.00 | 0.00 | 0.00 | 0.00 | 0.00 | 0.00 | 0.00 | 0.00 | 0.00 | 0.0 % | #N/A |
| *average(intraDomainStd)* | 0.00 | 0.00 | 0.00 | 0.00 | 0.00 | 0.00 | 0.00 | 0.00 | 0.00 | 0.00 | 0.00 | 0.0 % | #N/A |
| *average(intraDomainStdWt)* | 0.00 | 0.00 | 0.00 | 0.00 | 0.00 | 0.00 | 0.00 | 0.00 | 0.00 | 0.00 | 0.00 | 0.0 % | #N/A |
| *sumDomainBaseStack* | 0.00 | 0.00 | 0.00 | 0.00 | 0.00 | 0.00 | 0.00 | 0.00 | 0.00 | 0.00 | 0.00 | 0.0 % | #N/A |
| *avgDomainBaseStack* | -0.21 | -0.04 | 0.00 | -0.19 | -0.09 | -0.30 | -0.07 | -0.14 | 0.00 | -0.17 | -0.12 | 3.9 % | 82.1% |
| *wtAvgDomainBaseStack* | 0.00 | 0.00 | 0.00 | 0.00 | 0.00 | 0.00 | 0.00 | 0.00 | 0.00 | 0.00 | 0.00 | 0.0 % | #N/A |
| *numSingleCross* | 0.00 | 0.00 | 0.00 | 0.00 | 0.00 | 0.00 | 0.00 | 0.00 | 0.00 | 0.00 | 0.00 | 0.0 % | #N/A |
| *crossoverBaseStacking* | 0.00 | 0.00 | 0.00 | 0.00 | 0.00 | 0.00 | 0.00 | 0.00 | 0.00 | 0.00 | 0.00 | 0.0 % | #N/A |
| *EntropySkew* | -0.82 | -0.78 | -0.57 | -0.81 | -0.79 | -0.85 | -0.78 | -0.80 | -0.60 | -0.81 | -0.76 | 24.6 % | 12.6% |
| *LoopSkew* | 0.00 | 0.00 | -0.14 | 0.00 | 0.00 | 0.00 | 0.00 | 0.00 | -0.17 | 0.00 | -0.03 | 1.0 % | 211.5% |
| *sSkewTmWeighted* | 0.00 | 0.00 | 0.00 | 0.00 | 0.00 | 0.00 | 0.00 | 0.00 | 0.00 | 0.00 | 0.00 | 0.0 % | #N/A |
| *sSkewDomainWeighted* | 0.00 | 0.00 | 0.00 | 0.00 | 0.00 | 0.00 | 0.00 | 0.00 | 0.00 | 0.00 | 0.00 | 0.0 % | #N/A |

| | | | | | | | | | | | | | |
|---|---|---|---|---|---|---|---|---|---|---|---|---|---|
| *sum(blockList)* | 0.00 | 0.00 | 0.00 | 0.00 | 0.00 | 0.00 | 0.00 | 0.00 | 0.00 | 0.00 | 0.00 | 0.0 % | #N/A |
| *average(blockList)* | 0.00 | 0.00 | 0.00 | 0.00 | 0.00 | 0.00 | 0.00 | 0.00 | 0.00 | 0.00 | 0.00 | 0.0 % | #N/A |
| *skew(foldWeights)* | -0.27 | -0.16 | 0.00 | -0.26 | -0.19 | -0.34 | -0.18 | -0.23 | 0.00 | -0.24 | -0.19 | 6.0 % | 59.5% |
| *skew(domainWeights)* | 0.00 | 0.00 | 0.00 | 0.00 | 0.00 | 0.00 | 0.00 | 0.00 | 0.00 | 0.00 | 0.00 | 0.0 % | #N/A |
| *sum(foldWeights)* | 0.00 | 0.00 | 0.00 | 0.00 | 0.00 | 0.00 | 0.00 | 0.00 | 0.00 | 0.00 | 0.00 | 0.0 % | #N/A |
| *sum(domainWeights)* | -0.66 | -0.55 | -0.08 | -0.65 | -0.58 | -0.72 | -0.57 | -0.62 | -0.16 | -0.63 | -0.52 | 16.9 % | 41.8% |
| *crossoverCrossover BaseStacking* | 0.00 | 0.00 | 0.00 | 0.00 | 0.00 | 0.00 | 0.00 | 0.00 | 0.00 | 0.00 | 0.00 | 0.0 % | #N/A |
| *numDomains* | 0.00 | 0.00 | 0.00 | 0.00 | 0.00 | 0.00 | 0.00 | 0.00 | 0.00 | 0.00 | 0.00 | 0.0 % | #N/A |
| *invExp.AnnealTemp* | 0.00 | 0.00 | 0.00 | 0.00 | 0.00 | 0.00 | 0.00 | 0.00 | 0.00 | 0.00 | 0.00 | 0.0 % | #N/A |
| *invExp.MeltTemp* | 0.82 | 0.77 | 0.62 | 0.81 | 0.79 | 0.86 | 0.78 | 0.80 | 0.65 | 0.81 | 0.77 | 25.0 % | 9.9% |
| *invhystArea* | 0.00 | 0.00 | 0.00 | 0.00 | 0.00 | 0.00 | 0.00 | 0.00 | 0.00 | 0.00 | 0.00 | 0.0 % | #N/A |
| *invaverageFullStapleTm* | 0.00 | 0.00 | 0.00 | 0.00 | 0.00 | 0.00 | 0.00 | 0.00 | 0.00 | 0.00 | 0.00 | 0.0 % | #N/A |
| *invstdFullStapleTm* | 0.00 | 0.00 | 0.00 | 0.00 | 0.00 | 0.00 | 0.00 | 0.00 | 0.00 | 0.00 | 0.00 | 0.0 % | #N/A |
| *invaverageDomainTm* | 0.00 | 0.00 | 0.00 | 0.00 | 0.00 | 0.00 | 0.00 | 0.00 | 0.00 | 0.00 | 0.00 | 0.0 % | #N/A |
| *invstdDomainTm* | 0.00 | 0.00 | 0.00 | 0.00 | 0.00 | 0.00 | 0.00 | 0.00 | 0.00 | 0.00 | 0.00 | 0.0 % | #N/A |
| *invfoldAverageTm* | 0.00 | 0.00 | 0.00 | 0.00 | 0.00 | 0.00 | 0.00 | 0.00 | 0.00 | 0.00 | 0.00 | 0.0 % | #N/A |
| *invfoldStdTm* | 0.00 | 0.00 | 0.00 | 0.00 | 0.00 | 0.00 | 0.00 | 0.00 | 0.00 | 0.00 | 0.00 | 0.0 % | #N/A |
| *invavgFullStaplePlain* | 0.00 | 0.00 | 0.00 | 0.00 | 0.00 | 0.00 | 0.00 | 0.00 | 0.00 | 0.00 | 0.00 | 0.0 % | #N/A |
| *invstdFullStaplePlain* | -0.35 | -0.26 | -0.04 | -0.34 | -0.29 | -0.40 | -0.27 | -0.31 | -0.08 | -0.33 | -0.27 | 8.6 % | 44.2 % |
| *invaverage(intraDomainDiff _normalized)* | 0.00 | 0.00 | 0.00 | 0.00 | 0.00 | 0.00 | 0.00 | 0.00 | 0.00 | 0.00 | 0.00 | 0.0 % | #N/A |
| *invaverage(intraDomainSum)* | 0.00 | 0.00 | 0.00 | 0.00 | 0.00 | 0.00 | 0.00 | 0.00 | 0.00 | 0.00 | 0.00 | 0.0 % | #N/A |
| *invaverage(intraDomainStd)* | 0.00 | 0.00 | 0.00 | 0.00 | 0.00 | 0.00 | 0.00 | 0.00 | 0.00 | 0.00 | 0.00 | 0.0 % | #N/A |
| *invaverage(intraDomainStdWt)* | 0.00 | 0.00 | 0.00 | 0.00 | 0.00 | 0.00 | 0.00 | 0.00 | 0.00 | 0.00 | 0.00 | 0.0 % | #N/A |
| *invsumDomainBaseStack* | 0.00 | 0.00 | 0.00 | 0.00 | 0.00 | 0.00 | 0.00 | 0.00 | 0.00 | 0.00 | 0.00 | 0.0 % | #N/A |
| *invavgDomainBaseStack* | 0.00 | 0.00 | 0.00 | 0.00 | 0.00 | 0.00 | 0.00 | 0.00 | 0.00 | 0.00 | 0.00 | 0.0 % | #N/A |
| *invwtAvgDomainBaseStack* | 0.00 | 0.00 | 0.00 | 0.00 | 0.00 | 0.00 | 0.00 | 0.00 | 0.00 | 0.00 | 0.00 | 0.0 % | #N/A |
| *invnumSingleCross* | 0.00 | 0.00 | 0.00 | 0.00 | 0.00 | 0.00 | 0.00 | 0.00 | 0.00 | 0.00 | 0.00 | 0.0 % | #N/A |
| *invcrossoverBaseStacking* | 0.00 | 0.00 | 0.00 | 0.00 | 0.00 | 0.00 | 0.00 | 0.00 | 0.00 | 0.00 | 0.00 | 0.0 % | #N/A |
| *invEntropySkew* | 0.00 | 0.00 | 0.00 | 0.00 | 0.00 | 0.00 | 0.00 | 0.00 | 0.00 | 0.00 | 0.00 | 0.0 % | #N/A |
| *invLoopSkew* | 0.52 | 0.40 | 0.00 | 0.50 | 0.44 | 0.54 | 0.42 | 0.47 | 0.00 | 0.49 | 0.38 | 12.3 % | 53.9 % |
| *invsSkewTmWeighted* | 0.00 | 0.00 | 0.00 | 0.00 | 0.00 | 0.00 | 0.00 | 0.00 | 0.00 | 0.00 | 0.00 | 0.0 % | #N/A |
| *invsSkewDomainWeighted* | 0.00 | 0.00 | 0.00 | 0.00 | 0.00 | 0.13 | 0.00 | 0.00 | 0.00 | 0.00 | 0.01 | 0.4 % | 316.2 % |
| *invsum(blockList)* | 0.00 | 0.00 | 0.00 | 0.00 | 0.00 | 0.00 | 0.00 | 0.00 | 0.00 | 0.00 | 0.00 | 0.0 % | #N/A |
| *invaverage(blockList)* | 0.00 | 0.00 | 0.00 | 0.00 | 0.00 | 0.00 | 0.00 | 0.00 | 0.00 | 0.00 | 0.00 | 0.0 % | #N/A |
| *invskew(foldWeights)* | 0.00 | 0.00 | 0.00 | 0.00 | 0.00 | 0.00 | 0.00 | 0.00 | 0.00 | 0.00 | 0.00 | 0.0 % | #N/A |
| *invskew(domainWeights)* | 0.00 | 0.00 | 0.00 | 0.00 | 0.00 | 0.00 | 0.00 | 0.00 | 0.00 | 0.00 | 0.00 | 0.0 % | #N/A |
| *invsum(foldWeights)* | 0.00 | 0.00 | 0.00 | 0.00 | 0.00 | 0.00 | 0.00 | 0.00 | 0.00 | 0.00 | 0.00 | 0.0 % | #N/A |
| *invsum(domainWeights)* | 0.00 | 0.00 | 0.00 | 0.00 | 0.00 | 0.00 | 0.00 | 0.00 | 0.00 | 0.00 | 0.00 | 0.0 % | #N/A |
| *invCrossoverBaseStacking* | 0.00 | 0.00 | 0.00 | 0.00 | 0.00 | 0.00 | 0.00 | 0.00 | 0.00 | 0.00 | 0.00 | 0.0 % | #N/A |
| *invnumDomains* | 0.00 | 0.00 | 0.00 | 0.00 | 0.00 | 0.00 | 0.00 | 0.00 | 0.00 | 0.00 | 0.00 | 0.0 % | #N/A |

*Table 4: All Fitted coefficients, λ via cross-validation, Experimental $T_m$ & $T_a$ excluded*

| | *s1* | *s2* | *s3* | *s4* | *s5* | *s6* | *s7* | *s8* | *s9* | *s10* | *Avg.* | *% of Total* | *Rel. Std.* |
|---|---|---|---|---|---|---|---|---|---|---|---|---|---|
| *(Intercept)* | 1.24 | 2.03 | 1.94 | 1.96 | 2.03 | 1.40 | 1.99 | 2.03 | 1.92 | 1.99 | | | |
| *averageFullStapleTm* | 0.00 | 0.00 | 0.00 | 0.00 | 0.00 | 0.00 | 0.00 | 0.00 | 0.00 | 0.00 | 0.00 | 0.0 % | #N/A |
| *stdFullStapleTm* | 0.00 | 0.00 | 0.00 | 0.00 | 0.00 | 0.00 | 0.00 | 0.00 | 0.00 | 0.00 | 0.00 | 0.0 % | #N/A |
| *averageDomainTm* | 0.00 | 0.00 | 0.00 | 0.00 | 0.00 | 0.00 | 0.00 | 0.00 | 0.00 | 0.00 | 0.00 | 0.0 % | #N/A |
| *stdDomainTm* | 0.00 | 0.00 | 0.00 | 0.00 | 0.00 | 0.00 | 0.00 | 0.00 | 0.00 | 0.00 | 0.00 | 0.0 % | #N/A |
| *foldAverageTm* | 0.00 | 0.00 | 0.00 | 0.00 | 0.00 | 0.00 | 0.00 | 0.00 | 0.00 | 0.00 | 0.00 | 0.0 % | #N/A |
| *foldStdTm* | 0.00 | 0.25 | 0.26 | 0.20 | 0.25 | 0.00 | 0.26 | 0.25 | 0.22 | 0.26 | 0.20 | 8.4 % | 53.7 % |
| *avgFullStaplePlain* | 0.00 | 0.00 | 0.00 | 0.00 | 0.00 | 0.00 | 0.00 | 0.00 | 0.00 | 0.00 | 0.00 | 0.0 % | #N/A |
| *stdFullStaplePlain* | 0.00 | 0.00 | 0.00 | 0.00 | 0.00 | 0.00 | 0.00 | 0.00 | 0.00 | 0.00 | 0.00 | 0.0 % | #N/A |
| *avg.DomainStdbyStaple* | 0.00 | 0.00 | 0.00 | 0.00 | 0.00 | 0.00 | 0.00 | 0.00 | 0.00 | 0.00 | 0.00 | 0.0 % | #N/A |
| *average(intraDomainDiff_ normalized)* | 0.00 | 0.00 | 0.00 | 0.00 | 0.00 | 0.00 | 0.00 | 0.00 | 0.00 | 0.00 | 0.00 | 0.0 % | #N/A |
| *average(intraDomainSum)* | 0.00 | 0.00 | 0.00 | 0.00 | 0.00 | 0.00 | 0.00 | 0.00 | 0.00 | 0.00 | 0.00 | 0.0 % | #N/A |
| *average(intraDomainStd)* | 0.00 | 0.00 | 0.00 | 0.00 | 0.00 | 0.00 | 0.00 | 0.00 | 0.00 | 0.00 | 0.00 | 0.0 % | #N/A |
| *average(intraDomainStdWt)* | 0.00 | 0.00 | 0.00 | 0.00 | 0.00 | 0.00 | 0.00 | 0.00 | 0.00 | 0.00 | 0.00 | 0.0 % | #N/A |
| *sumDomainBaseStack* | 0.00 | 0.00 | 0.00 | 0.00 | 0.00 | 0.00 | 0.00 | 0.00 | 0.00 | 0.00 | 0.00 | 0.0 % | #N/A |
| *avgDomainBaseStack* | 0.00 | 0.00 | 0.00 | 0.00 | 0.00 | 0.00 | 0.00 | 0.00 | 0.00 | 0.00 | 0.00 | 0.0 % | #N/A |

| | | | | | | | | | | | | | |
|---|---|---|---|---|---|---|---|---|---|---|---|---|---|
| *wtAvgDomainBaseStack* | 0.00 | 0.00 | 0.00 | 0.00 | 0.00 | 0.00 | 0.00 | 0.00 | 0.00 | 0.00 | 0.00 | 0.0 % | #N/A |
| *numSingleCross* | 0.00 | 0.00 | 0.00 | 0.00 | 0.00 | 0.00 | 0.00 | 0.00 | 0.00 | 0.00 | 0.00 | 0.0 % | #N/A |
| *crossoverBaseStacking* | 0.00 | 0.00 | 0.00 | 0.00 | 0.00 | 0.00 | 0.00 | 0.00 | 0.00 | 0.00 | 0.00 | 0.0 % | #N/A |
| *EntropySkew* | 0.00 | -1.11 | -1.10 | -1.00 | -1.11 | -0.23 | -1.10 | -1.11 | -1.04 | -1.10 | -0.89 | 38.1 % | 46.4 % |
| *LoopSkew* | 0.00 | 0.00 | 0.00 | -0.14 | 0.00 | 0.00 | 0.00 | 0.00 | -0.07 | 0.00 | -0.02 | 0.9 % | 224.1 % |
| *sSkewTmWeighted* | 0.00 | 0.00 | 0.00 | 0.00 | 0.00 | 0.00 | 0.00 | 0.00 | 0.00 | 0.00 | 0.00 | 0.0 % | #N/A |
| *sSkewDomainWeighted* | 0.00 | 0.00 | 0.00 | 0.00 | 0.00 | 0.00 | 0.00 | 0.00 | 0.00 | 0.00 | 0.00 | 0.0 % | #N/A |
| *sum(blockList)* | 0.00 | 0.00 | 0.00 | 0.00 | 0.00 | 0.00 | 0.00 | 0.00 | 0.00 | 0.00 | 0.00 | 0.0 % | #N/A |
| *average(blockList)* | 0.00 | 0.00 | 0.00 | 0.00 | 0.00 | 0.00 | 0.00 | 0.00 | 0.00 | 0.00 | 0.00 | 0.0 % | #N/A |
| *skew(foldWeights)* | 0.00 | -0.10 | -0.03 | 0.00 | -0.10 | 0.00 | -0.07 | -0.10 | 0.00 | -0.07 | -0.05 | 1.9 % | 96.3% |
| *skew(domainWeights)* | 0.00 | 0.00 | 0.00 | 0.00 | 0.00 | 0.00 | 0.00 | 0.00 | 0.00 | 0.00 | 0.00 | 0.0 % | #N/A |
| *sum(foldWeights)* | 0.00 | 0.00 | 0.00 | 0.00 | 0.00 | 0.00 | 0.00 | 0.00 | 0.00 | 0.00 | 0.00 | 0.0 % | #N/A |
| *sum(domainWeights)* | 0.00 | -0.46 | -0.40 | -0.33 | -0.46 | 0.00 | -0.43 | -0.46 | -0.34 | -0.43 | -0.33 | 14.1 % | 54.6 % |
| *crossoverCrossoverBaseStacking* | 0.00 | 0.00 | 0.00 | 0.00 | 0.00 | 0.00 | 0.00 | 0.00 | 0.00 | 0.00 | 0.00 | 0.0 % | #N/A |
| *numDomains* | 0.00 | 0.00 | 0.00 | 0.00 | 0.00 | 0.00 | 0.00 | 0.00 | 0.00 | 0.00 | 0.00 | 0.0 % | #N/A |
| *invaverageFullStapleTm* | 0.00 | 0.68 | 0.67 | 0.61 | 0.68 | 0.00 | 0.67 | 0.68 | 0.63 | 0.67 | 0.53 | 22.5 % | 52.9% |
| *invstdFullStapleTm* | 0.00 | 0.00 | 0.00 | 0.00 | 0.00 | 0.00 | 0.00 | 0.00 | 0.00 | 0.00 | 0.00 | 0.0 % | #N/A |
| *invaverageDomainTm* | 0.00 | 0.00 | 0.00 | 0.00 | 0.00 | 0.00 | 0.00 | 0.00 | 0.00 | 0.00 | 0.00 | 0.0 % | #N/A |
| *invstdDomainTm* | 0.00 | 0.00 | 0.00 | 0.00 | 0.00 | 0.00 | 0.00 | 0.00 | 0.00 | 0.00 | 0.00 | 0.0 % | #N/A |
| *invfoldAverageTm* | 0.00 | 0.00 | 0.00 | 0.00 | 0.00 | 0.00 | 0.00 | 0.00 | 0.00 | 0.00 | 0.00 | 0.0 % | #N/A |
| *invfoldStdTm* | 0.00 | 0.00 | 0.00 | 0.00 | 0.00 | 0.00 | 0.00 | 0.00 | 0.00 | 0.00 | 0.00 | 0.0 % | #N/A |
| *invavgFullStaplePlain* | 0.00 | 0.00 | 0.00 | 0.00 | 0.00 | 0.00 | 0.00 | 0.00 | 0.00 | 0.00 | 0.00 | 0.0 % | #N/A |
| *invstdFullStaplePlain* | 0.00 | -0.13 | -0.11 | -0.09 | -0.13 | 0.00 | -0.12 | -0.13 | -0.10 | -0.12 | -0.09 | 3.9 % | 54.2 % |
| *invaverage(intraDomainDiff_normalized)* | 0.00 | 0.00 | 0.00 | 0.00 | 0.00 | 0.00 | 0.00 | 0.00 | 0.00 | 0.00 | 0.00 | 0.0 % | #N/A |
| *invaverage(intraDomainSum)* | 0.00 | 0.00 | 0.00 | 0.00 | 0.00 | 0.00 | 0.00 | 0.00 | 0.00 | 0.00 | 0.00 | 0.0 % | #N/A |
| *invaverage(intraDomainStd)* | 0.00 | 0.00 | 0.00 | 0.00 | 0.00 | 0.00 | 0.00 | 0.00 | 0.00 | 0.00 | 0.00 | 0.0 % | #N/A |
| *invaverage(intraDomainStdWt)* | 0.00 | 0.00 | 0.00 | 0.00 | 0.00 | 0.00 | 0.00 | 0.00 | 0.00 | 0.00 | 0.00 | 0.0 % | #N/A |
| *invsumDomainBaseStack* | 0.00 | 0.00 | 0.00 | 0.00 | 0.00 | 0.00 | 0.00 | 0.00 | 0.00 | 0.00 | 0.00 | 0.0 % | #N/A |
| *invavgDomainBaseStack* | 0.00 | 0.00 | 0.00 | 0.00 | 0.00 | 0.00 | 0.00 | 0.00 | 0.00 | 0.00 | 0.00 | 0.0 % | #N/A |
| *invwtAvgDomainBaseStack* | 0.00 | 0.00 | 0.00 | 0.00 | 0.00 | 0.00 | 0.00 | 0.00 | 0.00 | 0.00 | 0.00 | 0.0 % | #N/A |
| *invnumSingleCross* | 0.00 | 0.00 | 0.00 | 0.00 | 0.00 | 0.00 | 0.00 | 0.00 | 0.00 | 0.00 | 0.00 | 0.0 % | #N/A |
| *invcrossoverBaseStacking* | 0.00 | 0.00 | 0.00 | 0.00 | 0.00 | 0.00 | 0.00 | 0.00 | 0.00 | 0.00 | 0.00 | 0.0 % | #N/A |
| *invEntropySkew* | 0.00 | 0.00 | 0.00 | 0.00 | 0.00 | 0.00 | 0.00 | 0.00 | 0.00 | 0.00 | 0.00 | 0.0 % | #N/A |
| *invLoopSkew* | 0.00 | 0.37 | 0.33 | 0.08 | 0.37 | 0.00 | 0.35 | 0.37 | 0.18 | 0.35 | 0.24 | 10.2 % | 66.2 % |
| *invsSkewTmWeighted* | 0.00 | 0.00 | 0.00 | 0.00 | 0.00 | 0.00 | 0.00 | 0.00 | 0.00 | 0.00 | 0.00 | 0.0 % | #N/A |
| *invsSkewDomainWeighted* | 0.00 | 0.00 | 0.00 | 0.00 | 0.00 | 0.00 | 0.00 | 0.00 | 0.00 | 0.00 | 0.00 | 0.0 % | #N/A |
| *invsum(blockList)* | 0.00 | 0.00 | 0.00 | 0.00 | 0.00 | 0.00 | 0.00 | 0.00 | 0.00 | 0.00 | 0.00 | 0.0 % | #N/A |
| *invaverage(blockList)* | 0.00 | 0.00 | 0.00 | 0.00 | 0.00 | 0.00 | 0.00 | 0.00 | 0.00 | 0.00 | 0.00 | 0.0 % | #N/A |
| *invskew(foldWeights)* | 0.00 | 0.00 | 0.00 | 0.00 | 0.00 | 0.00 | 0.00 | 0.00 | 0.00 | 0.00 | 0.00 | 0.0 % | #N/A |
| *invskew(domainWeights)* | 0.00 | 0.00 | 0.00 | 0.00 | 0.00 | 0.00 | 0.00 | 0.00 | 0.00 | 0.00 | 0.00 | 0.0 % | #N/A |
| *invsum(foldWeights)* | 0.00 | 0.00 | 0.00 | 0.00 | 0.00 | 0.00 | 0.00 | 0.00 | 0.00 | 0.00 | 0.00 | 0.0 % | #N/A |
| *invsum(domainWeights)* | 0.00 | 0.00 | 0.00 | 0.00 | 0.00 | 0.00 | 0.00 | 0.00 | 0.00 | 0.00 | 0.00 | 0.0 % | #N/A |
| *invCrossoverBaseStacking* | 0.00 | 0.00 | 0.00 | 0.00 | 0.00 | 0.00 | 0.00 | 0.00 | 0.00 | 0.00 | 0.00 | 0.0 % | #N/A |
| *invnumDomains* | 0.00 | 0.00 | 0.00 | 0.00 | 0.00 | 0.00 | 0.00 | 0.00 | 0.00 | 0.00 | 0.00 | 0.0 % | #N/A |

*Table 5: All Fitted coefficients,* $\sum_i |\beta_i| = 3$

| *w/ Experimental $T_m$ & $T_a$* | *s1* | *% of Total* | *s1* | *% of Total* | *No Experimental $T_m$ or $T_a$* |
|---|---|---|---|---|---|
| *(Intercept)* | 1.90 | | 1.57 | | (Intercept) |
| *Exp.AnnealTemp* | 0.00 | 0.00 % | - | - | Exp.AnnealTemp |
| *Exp.MeltTemp* | 0.00 | 0.00 % | - | - | Exp.MeltTemp |
| *hystArea* | 0.00 | 0.00 % | - | - | hystArea |
| *averageFullStapleTm* | 0.00 | 0.00 % | 0.00 | 0.0 % | averageFullStapleTm |
| *stdFullStapleTm* | 0.00 | 0.00 % | 0.00 | 0.0 % | stdFullStapleTm |
| *averageDomainTm* | 0.00 | 0.00 % | 0.00 | 0.0 % | averageDomainTm |
| *stdDomainTm* | 0.01 | 0.29 % | 0.00 | 0.0 % | stdDomainTm |
| *foldAverageTm* | 0.00 | 0.00 % | 0.00 | 0.0 % | foldAverageTm |
| *foldStdTm* | 0.00 | 0.00 % | 0.26 | 8.6 % | foldStdTm |
| *avgFullStaplePlain* | 0.00 | 0.00 % | 0.00 | 0.0 % | avgFullStaplePlain |
| *stdFullStaplePlain* | 0.00 | 0.00 % | 0.00 | 0.0 % | stdFullStaplePlain |

| | | | | | |
|---|---|---|---|---|---|
| *avg.DomainStdbyStaple* | 0.00 | 0.00 % | 0.00 | 0.0 % | avg.DomainStdbyStaple |
| *average(intraDomainDiff_normalized)* | 0.00 | 0.00 % | 0.00 | 0.0 % | average(intraDomainDiff_normalized) |
| *average(intraDomainSum)* | 0.00 | 0.00 % | 0.00 | 0.0 % | average(intraDomainSum) |
| *average(intraDomainStd)* | 0.00 | 0.00 % | 0.00 | 0.0 % | average(intraDomainStd) |
| *average(intraDomainStdWt)* | 0.00 | 0.00 % | 0.00 | 0.0 % | average(intraDomainStdWt) |
| *sumDomainBaseStack* | 0.00 | 0.00 % | 0.00 | 0.0 % | sumDomainBaseStack |
| *avgDomainBaseStack* | -0.04 | 1.27 % | 0.00 | 0.0 % | avgDomainBaseStack |
| *wtAvgDomainBaseStack* | 0.00 | 0.00 % | 0.00 | 0.0 % | wtAvgDomainBaseStack |
| *numSingleCross* | 0.00 | 0.00 % | 0.00 | 0.0 % | numSingleCross |
| *crossoverBaseStacking* | 0.00 | 0.00 % | 0.00 | 0.0 % | crossoverBaseStacking |
| *EntropySkew* | -0.78 | 26.24 % | -1.10 | 36.8 % | EntropySkew |
| *LoopSkew* | 0.00 | 0.00 % | 0.00 | 0.0 % | LoopSkew |
| *sSkewTmWeighted* | 0.00 | 0.00 % | 0.00 | 0.0 % | sSkewTmWeighted |
| *sSkewDomainWeighted* | 0.00 | 0.00 % | 0.00 | 0.0 % | sSkewDomainWeighted |
| *sum(blockList)* | 0.00 | 0.00 % | 0.00 | 0.0 % | sum(blockList) |
| *average(blockList)* | 0.00 | 0.00 % | 0.00 | 0.0 % | average(blockList) |
| *skew(foldWeights)* | -0.16 | 5.37 % | -0.07 | 2.2 % | skew(foldWeights) |
| *skew(domainWeights)* | 0.00 | 0.00 % | 0.00 | 0.0 % | skew(domainWeights) |
| *sum(foldWeights)* | 0.00 | 0.00 % | 0.00 | 0.0 % | sum(foldWeights) |
| *sum(domainWeights)* | -0.55 | 18.50 % | -0.43 | 14.4 % | sum(domainWeights) |
| *crossoverCrossoverBaseStacking* | 0.00 | 0.00 % | 0.00 | 0.0 % | crossoverCrossoverBaseStacking |
| *numDomains* | 0.00 | 0.00 % | 0.00 | 0.0 % | numDomains |
| *invExp.AnnealTemp* | 0.00 | 0.00 % | - | - | invExp.AnnealTemp |
| *invExp.MeltTemp* | 0.77 | 26.09 % | - | - | invExp.MeltTemp |
| *invhystArea* | 0.00 | 0.00 % | - | - | invhystArea |
| *invaverageFullStapleTm* | 0.00 | 0.00 % | 0.67 | 22.4 % | invaverageFullStapleTm |
| *invstdFullStapleTm* | 0.00 | 0.00 % | 0.00 | 0.0 % | invstdFullStapleTm |
| *invaverageDomainTm* | 0.00 | 0.00 % | 0.00 | 0.0 % | invaverageDomainTm |
| *invstdDomainTm* | 0.00 | 0.00 % | 0.00 | 0.0 % | invstdDomainTm |
| *invfoldAverageTm* | 0.00 | 0.00 % | 0.00 | 0.0 % | invfoldAverageTm |
| *invfoldStdTm* | 0.00 | 0.00 % | 0.00 | 0.0 % | invfoldStdTm |
| *invavgFullStaplePlain* | 0.00 | 0.00 % | 0.00 | 0.0 % | invavgFullStaplePlain |
| *invstdFullStaplePlain* | -0.26 | 8.64 % | -0.12 | 4.0 % | invstdFullStaplePlain |
| *invaverage(intraDomainDiff_normalized)* | 0.00 | 0.00 % | 0.00 | 0.0 % | invaverage(intraDomainDiff_normalized) |
| *invaverage(intraDomainSum)* | 0.00 | 0.00 % | 0.00 | 0.0 % | invaverage(intraDomainSum) |
| *invaverage(intraDomainStd)* | 0.00 | 0.00 % | 0.00 | 0.0 % | invaverage(intraDomainStd) |
| *invaverage(intraDomainStdWt)* | 0.00 | 0.00 % | 0.00 | 0.0 % | invaverage(intraDomainStdWt) |
| *invsumDomainBaseStack* | 0.00 | 0.00 % | 0.00 | 0.0 % | invsumDomainBaseStack |
| *invavgDomainBaseStack* | 0.00 | 0.00 % | 0.00 | 0.0 % | invavgDomainBaseStack |
| *invwtAvgDomainBaseStack* | 0.00 | 0.00 % | 0.00 | 0.0 % | invwtAvgDomainBaseStack |
| *invnumSingleCross* | 0.00 | 0.00 % | 0.00 | 0.0 % | invnumSingleCross |
| *invcrossoverBaseStacking* | 0.00 | 0.00 % | 0.00 | 0.0 % | invcrossoverBaseStacking |
| *invEntropySkew* | 0.00 | 0.00 % | 0.00 | 0.0 % | invEntropySkew |
| *invLoopSkew* | 0.40 | 13.61 % | 0.35 | 11.6 % | invLoopSkew |
| *invsSkewTmWeighted* | 0.00 | 0.00 % | 0.00 | 0.0 % | invsSkewTmWeighted |
| *invsSkewDomainWeighted* | 0.00 | 0.00 % | 0.00 | 0.0 % | invsSkewDomainWeighted |
| *invsum(blockList)* | 0.00 | 0.00 % | 0.00 | 0.0 % | invsum(blockList) |
| *invaverage(blockList)* | 0.00 | 0.00 % | 0.00 | 0.0 % | invaverage(blockList) |
| *invskew(foldWeights)* | 0.00 | 0.00 % | 0.00 | 0.0 % | invskew(foldWeights) |
| *invskew(domainWeights)* | 0.00 | 0.00 % | 0.00 | 0.0 % | invskew(domainWeights) |
| *invsum(foldWeights)* | 0.00 | 0.00 % | 0.00 | 0.0 % | invsum(foldWeights) |
| *invsum(domainWeights)* | 0.00 | 0.00 % | 0.00 | 0.0 % | invsum(domainWeights) |
| *invCrossoverBaseStacking* | 0.00 | 0.00 % | 0.00 | 0.0 % | invcCrossoverBaseStacking |
| *invnumDomains* | 0.00 | 0.00 % | 0.00 | 0.0 % | invnumDomains |

#### Section 4: Organization of design information

Information on the individual designs are provided in section 17 of the SI in the form of page long datasheets. These contain the routing pattern, staple motif, final calculated ϕ, percentages of yield classifications, circle plots (depicting staple fold distances across the scaffold), square plots (depicting fold nestedness), and histograms of fold distances, entropic penalties and *predicted* $T_m$s for each fold.

Staple List: A list of the individual staple sequences for that design. Provided as a separate set of files.

Sequence topologies: A list of the individual staple sequences divided into staple domains with scaffold positions to indicate fold distances, looping distances, and calculated j-factors.

*Table 6: Descriptions of designs and their organization into related families*

| # | Helix θ | Routing | | Design Details |
|---|---|---|---|---|
| 1-10 | 90° (π/2 rad) | Horizontal Routing | Staple Motif: 8-16-8 | **Scaff. start position**:<br>**(#1)** 0 nt, **(#2)** -672 nt, **(#3)** -168 nt, **(#4)** +168 nt, **(#5)** +184 nt, **(#6)** +368 nt,<br>**(#7)** +736 nt, **(#8)** +2360 nt, **(#9)** +3680 nt, **(#10)** +5630 nt |
| 11,12 | | | | **Excess ssDNA divided into multiple start positions:**<br>**(#11)** +736 nt & +2360 nt, **(#12)** -672 nt, +736 nt, & +2360 nt |
| 13-16 | | | | **Seam shifted,** distance from edge**:**<br>**(#13)** 240 nt, **(#14)** 144 nt, **(#15)** 16 nt, **(#16)** 162 nt with staple motif flaw |
| 17-20 | | | | **Seam interdigitated,** distance from edge**:**<br>**(#17)** 240 nt, **(#18)** 144 nt, **(#19)** 16 nt, **(#20)** 272 nt |
| 21 | | | | **(#21) 3 Seam Routing** |
| 22 | | | | **(#22) Notched Rectangle Routing** |
| 23 | | | | **(#23) Weave Routing: Staple Motif:** 16-16 |
| 24-26 | | | | **Vertical Routing: Staple Motif: (#24)** 16-16, **(#25)** 8-16-8, **(#26)** 8-16-16-16-8 |
| 27, 28 | 120° (2/3 π) | | | **Horizontal Routing: Staple Motif:** domain heterogeneity<br>**(#27)** homogeneous [7-7-7-7-7-7-7-7, 2x(14-14-14-14)]<br>**(#28)** heterogeneous[2x(7-7-7-14),2x(7-14-14),14-14] |
| 29 | | | | **Vertical Routing: Staple Motif:**<br>**(#29)** heterogeneous domains [2x(7-7-7-14),2x(7-14-14),14-14] |
| 30-34 | 90° (π/2 ) | | | **Staple Motif:**<br>**(#30)** 16-16, **(#1)** 8-16-8, **(#31)** 16-16-16, **(#32)** 8-16-16-8, **(#33)** 8-16-16-16-8,<br>**(#34)** 8-32-16-8 |

Description of Plots in worksheets:

*Routing Plot:* The routing plot depicts how the circular scaffold routes through an origami. The X on a routing plot indicates both the start of the M13MP18 sequence, as well as the position where the ~100 bases of excess scaffold (which are not consumed by folding) enter and exit the design.

*Circle plot:* the circle plots depict the scaffold as a circular ring, where the bottom of the plot corresponds to the sequence start/stop (indicated by an X in the routing plot). Each fold within each staple is denoted by an arc bridging two small data markers, where the small data markers are the bases on the scaffold that the two staple domains force into proximity.

*Square Plot:* The square plots are made by depicting the scaffold as a line, where the beginning of the scaffold sequence is at 0,0 with the sequence progressing rightwards. Each fold is organized by the two scaffold positions it forces into proximity, indicated by small datamarkers, with a y-axis value corresponding to the total distance being bridged by the staple fold. While this is somewhat information redundant, it clearly depicts folding pathways beginning at the shortest folds on the top, down to progressively longer folds.

#### Section 5: Staple pool concentration normalization

The yield, y, of perfectly synthesized DNA oligomers is typically modeled as in Eq. 4 below, where addition of each nucleotide to the oligomer is treated as a reaction with a single coupling efficiency, given as e below. The coupling efficiency is typically advertised as >99.5 %. This is repeated n times, where n is the number of nucleotides in the oligomer.

*Eq. 4:* $$y = e^n \text{ where } e \approx 99.5\ \%$$

As the number of nucleotides in a strand increases, the yield decreases dramatically. Typical staples between 24 to 48 bases in length therefore have yield of non-truncated oligomers between 89 % to 79 % respectively.

Truncated staple strands will present an additional kinetic trap, and as longer strands of DNA are synthesized in lower yield. Staple pools are rarely, if ever, gel purified to remove truncated oligomer products, and doing so is expensive. This presents a challenge in comparing the yield of DNA origami with staple motifs of different average length.

Fully addressing this challenge is beyond the scope of this work. As a moderate mitigation step we chose to normalize the various staple pools to the anticipated concentration of correctly synthesized oligomer.

To do so rigorously, we wrote a simple python script to randomly generate thousands of truncated oligomer sequences by randomly removing bases. The ultraviolet absorption of these strands was then calculated by Cavaluzzi and Borer[9]. These extinction coefficients were then weighted by the probability of that number truncations.

When calculating staple pool concentration, these weighted extinction coefficients were used so the concentration reflected the anticipated yield of correctly synthesized staples, neglecting truncated products.

#### Section 6: Imaging yield procedure and classification

Figure 2 below is an example of image classification performed in this work. The origami were labeled in powerpoint, or with pen and paper and classified as well-folded, damaged, ripped, or catastrophic as described in the main text.

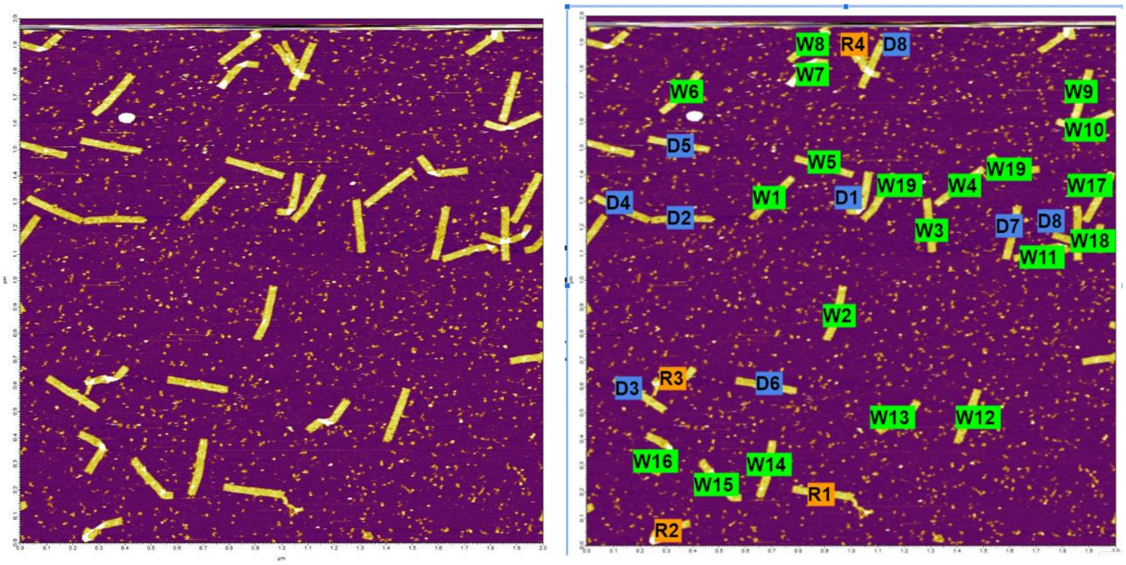

*Figure 2: Example classification of origami into well-folded, damaged, ripped, and catastrophic (in this image 19 origami were classified as well-folded, 9 were classified as damaged, 4 were classified as ripped, and none were classified as catastrophic*

These total number of each classification was then transcribed into a spreadsheet, given as a separate supplemental file, and summarized in SI section 7".

#### Section 7: Raw Experimental Counts

As described in the main text, Analyst 3 ( the first author) classified all the designs and annealing conditions presented in the text. The images for designs #1 through #7, #9, #33, and #34 were also classified by Analysts 1, 2 and 4. The raw counts of Well-Folded, Damaged, Ripped, and Catastrophic origami, as well as the number of images and origami per image are reported in the tables below.

*Table 7: Raw origami counts for all designs by analyst #3*

| Design # | Name | Well-Folded | Damaged | Ripped | Catastrophic | Total | # images | Origami Per image | ϕ |
|---|---|---|---|---|---|---|---|---|---|
| 1 | *mid0000* | 57 | 21 | 27 | 6 | 111 | 6 | 18.5 | 7.8 |
| 2 | *1FL0000* | 28 | 46 | 134 | 173 | 381 | 6 | 63.5 | 28.7 |
| 3 | *n168_0000* | 39 | 130 | 86 | 14 | 269 | 7 | 38.4 | 10.0 |
| 4 | *p168_0000* | 25 | 66 | 37 | 9 | 137 | 8 | 17.1 | 9.9 |
| 5 | *p184_0000* | 92 | 95 | 78 | 13 | 278 | 8 | 34.8 | 8.6 |
| 6 | *p368_0000* | 2 | 16 | 54 | 49 | 121 | 8 | 15.1 | 27.6 |
| 7 | *1FR* | 16 | 5 | 14 | 121 | 156 | 11 | 14.2 | 40.4 |
| 8 | *H_2368* | 2 | 13 | 137 | 102 | 254 | 8 | 31.8 | 28.4 |
| 9 | *midFlip0000* | 35 | 54 | 45 | 16 | 150 | 6 | 25.0 | 11.9 |
| 10 | *H5630* | 36 | 31 | 41 | 6 | 114 | 12 | 9.5 | 9.7 |
| 11 | *X2Goatee* | 4 | 3 | 105 | 113 | 225 | 5 | 45.0 | 32.2 |
| 12 | *X3Goatee* | 0 | 3 | 208 | 163 | 374 | 5 | 74.8 | 30.2 |
| 13 | *Shift_16* | 91 | 88 | 126 | 22 | 327 | 6 | 54.5 | 10.8 |
| 14 | *Shift_144* | 14 | 62 | 332 | 21 | 429 | 5 | 85.8 | 14.8 |
| 15 | *Shift_240* | 86 | 131 | 167 | 14 | 398 | 6 | 66.3 | 9.9 |

| | | | | | | | | | |
|---|---|---|---|---|---|---|---|---|---|
| 16 | *midShift0000* | 7 | 10 | 57 | 80 | 154 | 6 | 25.7 | 31.9 |
| 17 | *ID_16* | 23 | 68 | 175 | 70 | 336 | 9 | 37.3 | 16.7 |
| 18 | *ID_144* | 20 | 107 | 259 | 60 | 446 | 5 | 89.2 | 12.7 |
| 19 | *ID_260* | 33 | 73 | 168 | 15 | 289 | 4 | 72.3 | 5.0 |
| 20 | *ID_272* | 68 | 92 | 78 | 10 | 248 | 10 | 24.8 | 8.9 |
| 21 | *X3Seam* | 59 | 135 | 183 | 17 | 394 | 6 | 65.7 | 11.0 |
| 22 | *NR* | 8 | 111 | 248 | 130 | 497 | 7 | 71.0 | 21.7 |
| 23 | *HW_16-16* | 26 | 75 | 200 | 60 | 361 | 7 | 51.6 | 17.7 |
| 24 | *v-8-16-8* | 0 | 10 | 98 | 50 | 158 | 8 | 19.8 | 25.4 |
| 25 | *V1616* | 32 | 96 | 307 | 40 | 475 | 8 | 59.4 | 15.0 |
| 26 | *v8-16-16-168* | 10 | 73 | 186 | 17 | 286 | 8 | 35.8 | 14.0 |
| 27 | *honeyH_bad* | 18 | 75 | 129 | 7 | 229 | 8 | 28.6 | 11.7 |
| 28 | *honeyV_Mix* | 0 | 17 | 110 | 66 | 193 | 7 | 27.6 | 26.1 |
| 29 | *honeyH_Mix* | 17 | 31 | 159 | 56 | 263 | 10 | 26.3 | 20.4 |
| 30 | *H16_16* | 21 | 73 | 71 | 8 | 173 | 7 | 24.7 | 10.7 |
| 31 | *H_16-16-16* | 126 | 139 | 135 | 12 | 412 | 7 | 58.9 | 19.3 |
| 32 | *H_8-16-16-8* | 89 | 145 | 266 | 32 | 532 | 7 | 76.0 | 8.4 |
| 33 | *P8-16-16-16-8_0000* | 158 | 43 | 30 | 7 | 238 | 7 | 34.0 | 4.9 |
| 34 | *p32-16-8_0000* | 82 | 140 | 258 | 58 | 538 | 14 | 38.4 | 14.0 |

*Table 8: Raw origami counts for the analyst-to-analyst control designs from analyst #1*

| Design # | Name | Well-Folded | Damaged | Ripped | Catastrophic | Total | # images | Origami Per image | ϕ |
|---|---|---|---|---|---|---|---|---|---|
| 1 | mid0000 | 64 | 40 | 14 | 5 | 123 | 6 | 20.5 | 5.9 |
| 2 | 1FL0000 | 23 | 71 | 92 | 60 | 246 | 6 | 41.0 | 19.3 |
| 3 | n168_0000 | 33 | 179 | 60 | 5 | 277 | 7 | 39.6 | 7.5 |
| 4 | p168_0000 | 26 | 75 | 25 | 2 | 128 | 8 | 16.0 | 6.8 |
| 5 | p184_0000 | 79 | 160 | 74 | 10 | 323 | 8 | 40.4 | 7.7 |
| 6 | p368_0000 | 10 | 17 | 51 | 42 | 120 | 8 | 15.0 | 24.7 |
| 7 | 1FR | 13 | 14 | 12 | 116 | 155 | 11 | 14.1 | 39.1 |
| 9 | midFlip0000 | 37 | 78 | 50 | 13 | 178 | 6 | 29.7 | 10.3 |
| 16 | midShift0000 | 1 | 33 | 46 | 84 | 164 | 6 | 27.3 | 30.8 |
| 33 | P8-16-16-16-8_0000 | 121 | 116 | 13 | 3 | 253 | 7 | 36.1 | 4.1 |
| 34 | p32-16-8_0000 | 42 | 238 | 257 | 34 | 571 | 14 | 40.8 | 11.9 |

*Table 9: Raw origami counts for the analyst-to-analyst control designs from analyst #2*

| Design # | Name | Well-Folded | Damaged | Ripped | Catastrophic | Total | # images | Origami Per image | ϕ |
|---|---|---|---|---|---|---|---|---|---|

| | | | | | | | | | |
|---|---|---|---|---|---|---|---|---|---|
| 1 | mid0000 | 109 | 22 | 4 | 1 | 136 | 6 | 22.7 | 2.4 |
| 2 | 1FL0000 | 82 | 125 | 102 | 149 | 458 | 6 | 76.3 | 21.2 |
| 3 | n168_0000 | 147 | 118 | 29 | 4 | 298 | 7 | 42.6 | 4.6 |
| 4 | p168_0000 | 65 | 76 | 14 | 1 | 156 | 8 | 19.5 | 4.5 |
| 5 | p184_0000 | 204 | 126 | 21 | 2 | 353 | 8 | 44.1 | 3.5 |
| 6 | p368_0000 | 21 | 38 | 57 | 23 | 139 | 8 | 17.4 | 15.9 |
| 7 | 1FR | 23 | 5 | 24 | 122 | 174 | 11 | 15.8 | 37.4 |
| 9 | midFlip0000 | 90 | 78 | 19 | 3 | 190 | 6 | 31.7 | 4.8 |
| 16 | midShift0000 | 33 | 50 | 45 | 49 | 177 | 6 | 29.5 | 19.3 |
| 33 | P8-16-16-16-8_0000 | 228 | 42 | 4 | 1 | 275 | 7 | 39.3 | 2.0 |
| 34 | p32-16-8_0000 | 201 | 250 | 238 | 9 | 698 | 14 | 49.9 | 7.8 |

*Table 10:Raw origami counts for the analyst-to-analyst control designs from analyst #4*

| Design # | Name | Well-Folded | Damaged | Ripped | Catastrophic | Total | # images | Origami Per image | ϕ |
|---|---|---|---|---|---|---|---|---|---|
| 1 | mid0000 | 27 | 53 | 20 | 5 | 105 | 6 | 17.5 | 8.0 |
| 2 | 1FL0000 | 10 | 75 | 134 | 213 | 432 | 6 | 72.0 | 30.2 |
| 3 | n168_0000 | 4 | 187 | 94 | 10 | 295 | 7 | 42.1 | 9.7 |
| 4 | p168_0000 | 6 | 84 | 50 | 5 | 145 | 8 | 18.1 | 9.8 |
| 5 | p184_0000 | 29 | 186 | 98 | 19 | 332 | 8 | 41.5 | 10.2 |
| 6 | p368_0000 | 0 | 15 | 68 | 39 | 122 | 8 | 15.3 | 25.0 |
| 7 | 1FR | 5 | 19 | 0 | 169 | 193 | 11 | 17.5 | 44.3 |
| 9 | midFlip0000 | 14 | 69 | 80 | 9 | 172 | 6 | 28.7 | 11.7 |
| 16 | midShift0000 | 0 | 24 | 63 | 83 | 170 | 6 | 28.3 | 30.7 |
| 33 | P8-16-16-16-8_0000 | 59 | 156 | 17 | 3 | 235 | 7 | 33.6 | 5.3 |
| 34 | p32-16-8_0000 | 12 | 164 | 407 | 43 | 626 | 14 | 44.7 | 14.5 |

*Table 11: Raw origami counts for excess staple concentration controls, from analyst #3*

| Design # | Name | Well-Folded | Damaged | Ripped | Catastrophic | Total | # images | Origami Per image | ϕ |
|---|---|---|---|---|---|---|---|---|---|
| 9 | *Mid_10xStap* | 613 | 169 | 216 | 7 | 1005 | 12 | 83.8 | 3.3 |
| 9 | *Mid_100xStap* | 592 | 68 | 97 | 3 | 760 | 10 | 76.0 | 18.3 |
| 7 | *OFR_10xStap* | 46 | 205 | 584 | 63 | 898 | 9 | 99.8 | 14.5 |
| 7 | *OFR_100xStap* | 533 | 233 | 315 | 8 | 1089 | 11 | 99.0 | 6.3 |

#### Section 8: Uncertainty in imaging yield

*Sample size uncertainty*: this uncertainty was evaluated by monte carlo simulation. For any design the experimental population percentages were taken as a ground truth, and 2 000 simulations were run; in each simulation the number of experimentally imaged origami were pulled randomly from the ground truth distribution. The $\phi$ value for each of these simulations was calculated, and the standard deviation in the simulated distribution of $\phi$ was used to represent the uncertainty in experimental $\phi$ associated with sample size. The sample size uncertainty contributed from 0.2 to 1.7 to $\phi$.

*Day-to-Day variation in classification*: A single image was reclassified 9 times, with few to no reclassifications on the same day. This image had only 29 origami to facilitate tracking of variation in classification of individual origami and was randomly rotated and flipped to minimize analyst recognition.

A key image was created to index the origami, Figure 3 below. Images of the classifications and appropriately rotated key are given in Figure 4.

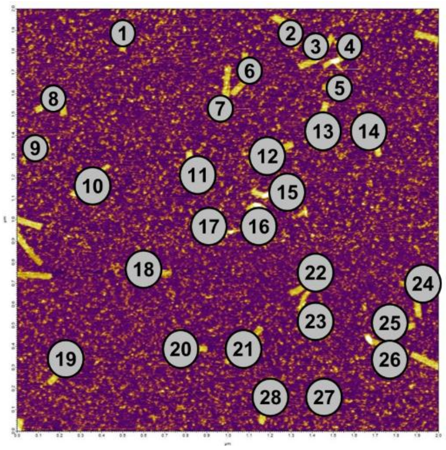

*Figure 3: Image key to identify individual origami*

The base, unnumbered image, was then repeatedly re-classified as below. After classification the key image was copied and used to register which origami were classified differently.

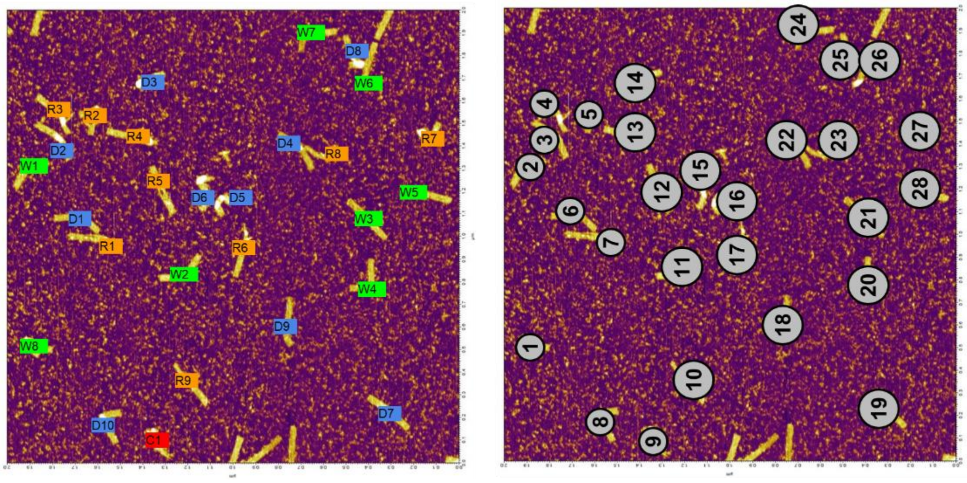

day 1

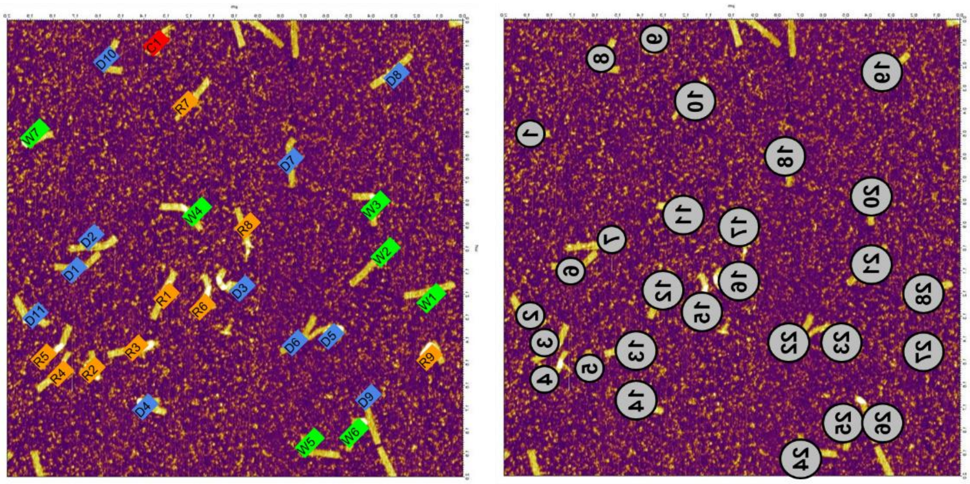

day 2

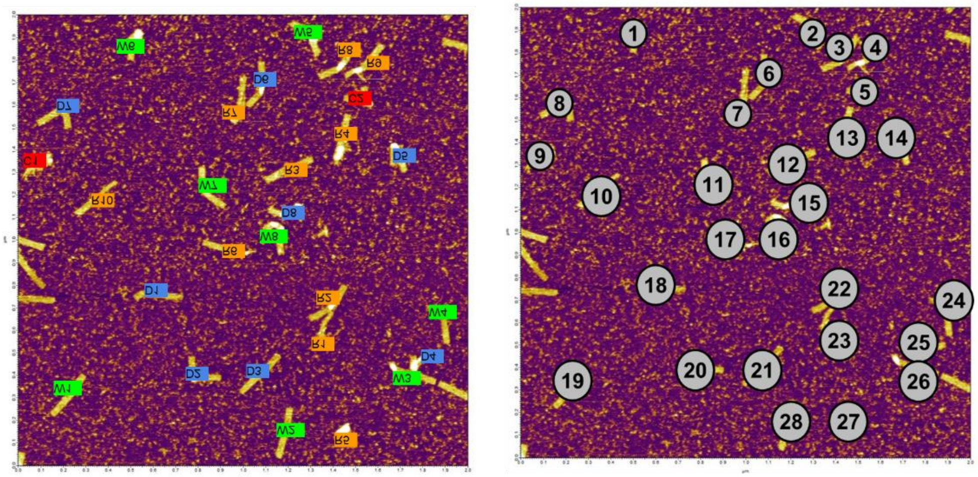

day 3

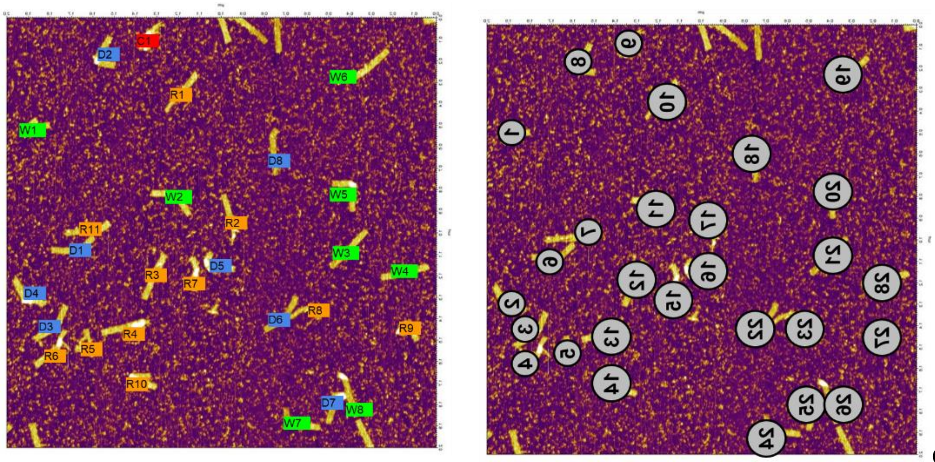

day 4

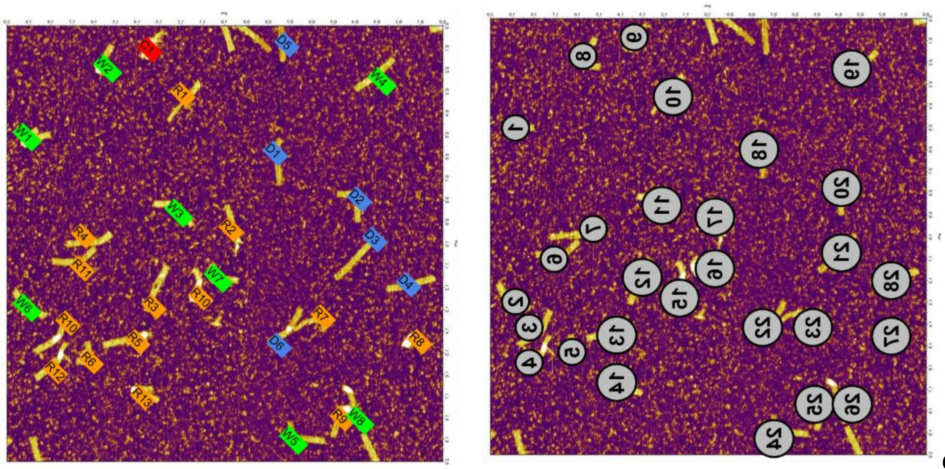

day 5

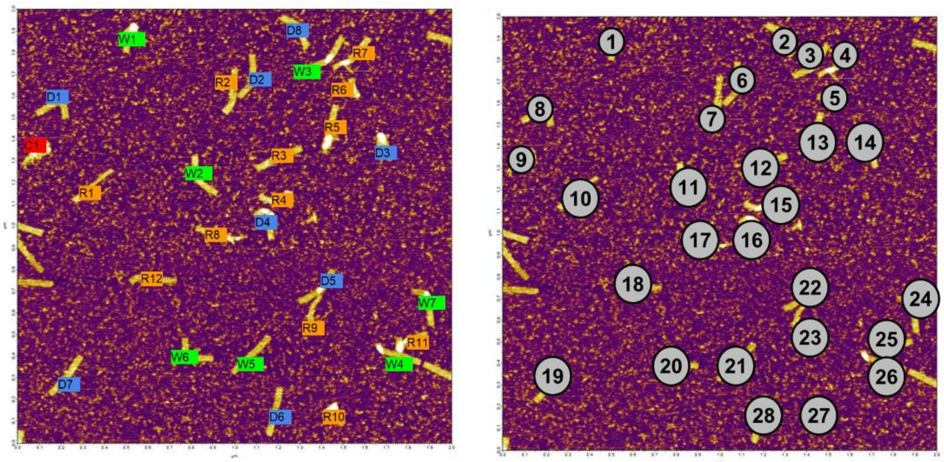

day 6

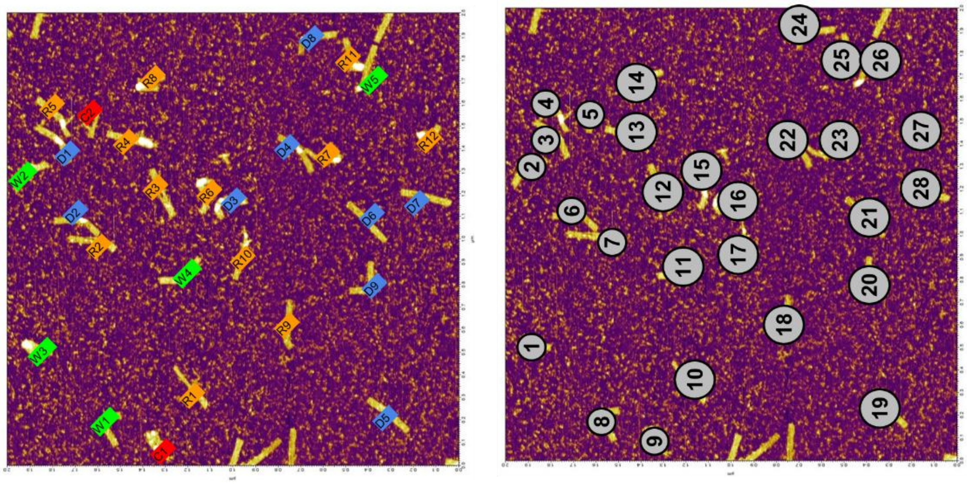

day 7

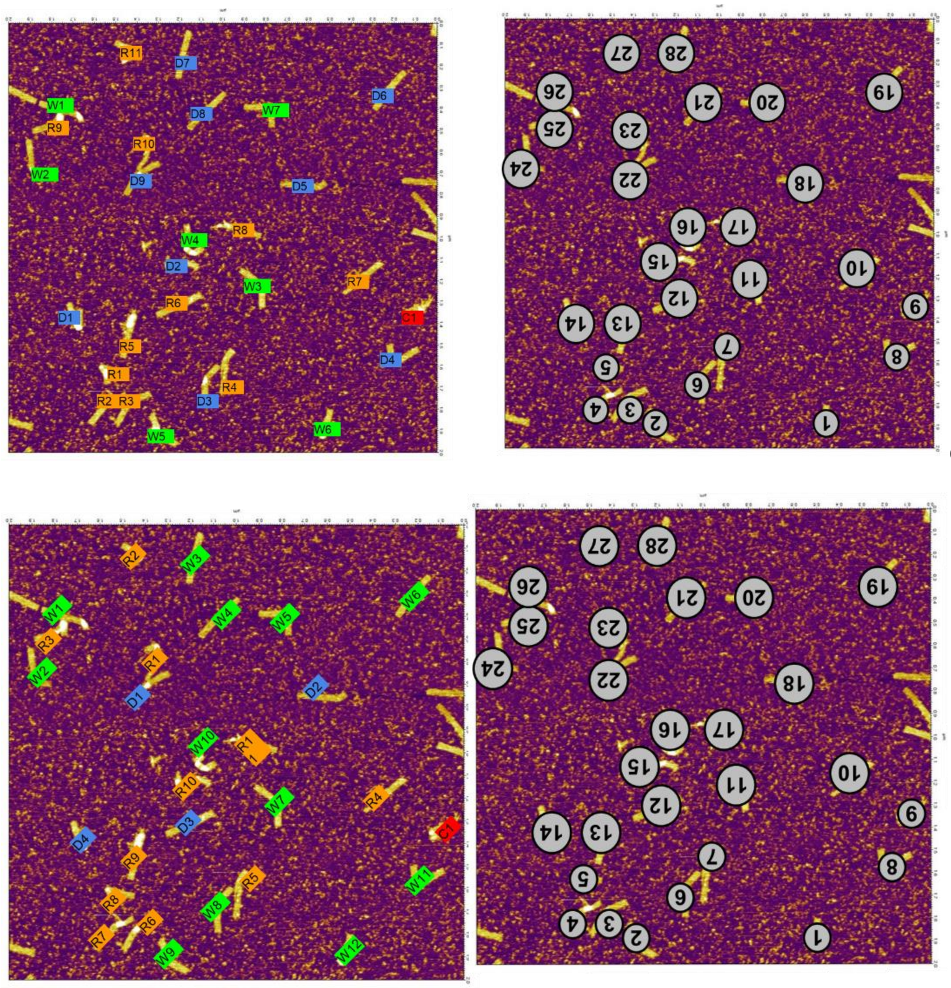

day 8

day 9

*Figure 4: repeated classifications of a single image to quantify day-to-day uncertainty*

Table 12 shows the counts of the reclassifications given in *Figure 1*. While some days, for example day 9, or day 7, varied significantly from the mean, the re-classifications were largely self-consistent.

*Table 12: Classification counts for day-today controls*

| | # Well-Folded | # Damaged | # Ripped | # Catastrophic | Total | | φ |
|---|---|---|---|---|---|---|---|
| **Day1** | 8 | 10 | 9 | 1 | 28 | | 8.68 |
| **Day2** | 7 | 11 | 9 | 1 | 28 | | 8.82 |
| **Day3** | 8 | 8 | 10 | 2 | 28 | | 10.64 |
| **Day4** | 8 | 8 | 11 | 1 | 28 | | 9.39 |
| **Day5** | 8 | 6 | 13 | 1 | 28 | | 10.11 |
| **Day6** | 7 | 8 | 12 | 1 | 28 | | 9.89 |
| **Day7** | 5 | 9 | 12 | 2 | 28 | | 11.79 |
| **Day8** | 7 | 9 | 11 | 1 | 28 | | 9.54 |
| **Day9** | 12 | 4 | 11 | 1 | 28 | | 8.82 |

| | | | | | | | |
|---|---|---|---|---|---|---|---|
| **Avg.** | 7.78 | 8.11 | 10.89 | 1.22 | | | 9.74 |
| **Std. Dev** | 1.75 | 1.97 | 1.29 | 0.42 | | | 0.95 |
| **Min** | 5 | 4 | 9 | 1 | | | ➔ 9 marginal origami |
| **Max** | 12 | 11 | 13 | 2 | | | |

Collectively, this system exhibits a larger variation in ϕ than whole origami samples; this is because the catastrophic origami are heavily weighted and because there are only 28 origami in this image. However, when the origami # (key images with grey circles) it can be seen that a core set of origami for each classification is constant, and that it is only the marginal cases which are classified into different categories on a day-to-day basis.

We take the minimum number of origami in each classification as the number that are unquestionably of that classification, and not marginal. This is 9 origami, as in the table above. It is also worth noting that of the four classifications, the Ripped origami had the lowest variability relative to their total number. This makes sense as this classification is the most distinctive of the four.

To estimate the number of marginal origami which could be classified into either neighboring category we began with the well-folded and catastrophic states. As any difference between the minimum and maximum number of origami between re-classifications for those must by definition come from the pool of marginal origami, that difference was used to estimate the number of marginal origami between the well-folded and damaged categories and between the ripped and catastrophic categories. The marginal origami between the damaged and ripped categories were then assumed to be the remainder of the 28 origami.

| Definitive Well-Folded | Marginal W or D | Definitive Damaged | Marginal D or R | Definitive Ripped | Marginal R or C | Definitive Catastrophic | |
|---|---|---|---|---|---|---|---|
| 5 | 7 | 4 | 1 | 9 | 1 | 1 | 28 |

| % (relative to the two neighboring categories) | |
|---|---|
| Marginal WD | 43.8 % |
| Marginal DR | 7.1 % |
| Marginal RC | 9.1 % |

We used these percentages to inform our Monte Carlo simulations and to estimate day-to-day user uncertainty. Because images were classified in random order, we assume each was classified on a different day. As in sampling uncertainty, we took the classified counts of each origami image to be a ground truth, then used the percentages above to calculate the number of marginal origami for that image in each of 1), 2), and 3). These marginal origami were then randomly reclassified into the neighboring categories randomly.

In our Monte Carlo simulations this was represented by using the total number of images and average origami per image for each design. The experimentally counted origami were used as a ground truth, the fraction of marginal origami was calculated, then for each image the marginal origami were randomly re-assigned to a neighboring category. This process was repeated 2,000 times.

The day-to-day variation contributed between 0.1 to 0.3 to ϕ.

*Staple pool synthesis variation:* This source of variation was quantified by ordering a single staple pool for a single design, #9, five times. These pools were then imaged and classified as separate samples, counts below.

*Table 13: Staple pool synthesis variation controls, raw and simulated counts*

| | *Well-Folded* | *Damaged* | *Ripped* | *Catastrophic* | *Total origami* | *Avg. Origami/image* | *Phi* |
|---|---|---|---|---|---|---|---|
| Pool1 | 23 | 55 | 254 | 35 | 367 | 73.4 | 16.0 |
| *Pool2* | 62 | 101 | 368 | 64 | 595 | 99.16 | 15.6 |
| *Pool3* | 3 | 39 | 179 | 58 | 279 | 55.8 | 20.7 |
| *Pool4* | 19 | 43 | 179 | 47 | 288 | 36 | 18.3 |
| *Pool5* | 35 | 54 | 45 | 16 | 150 | 25 | 11.9 |
| *Dummy1* | 32 | 64 | 225 | 46 | 367 | 73.4 | 16.4 |
| *Dummy2* | 53 | 103 | 363 | 76 | 595 | 99.2 | 16.5 |
| *Dummy3* | 24 | 49 | 171 | 35 | 279 | 55.8 | 16.4 |
| *Dummy4* | 24 | 52 | 176 | 36 | 288 | 36 | 16.4 |
| *Dummy5* | 13 | 27 | 91 | 19 | 150 | 25 | 16.4 |

In order to extract the staple pool variation from the variation between pools 1 through 5 of design #9, the sample size, and day-to-day sources of variation had to be removed. To do this, 5 dummy datasets were made. These dummy datasets had the same defectivity as the average of the pools 1 through 5, and had the same total origami and average origami/image.

The variation in defectivity between these dummy datasets was subtracted in quadrature from the variation in defectivity in pools 1 through 5. This should remove the contributions of sample size and day-to-day sources of variation from the staple pool synthesis variation. The staple pool synthesis variation contributed a variation of 2.8 to ϕ in this sample.

*The staple pool synthesis variation was much larger than we had anticipated and comprised the majority of our reported uncertainty in φ.* We imaged approximately 200 origami to 400 origami for each design with an uncertainty of 2.8 to 3.3 in ϕ. An optimized protocol which accounted for this source of variation would have imaged only ≈ 30 origami from 10 separate pools for each design; this would have halved the total uncertainty in ϕ. Future, higher throughput, experiments using image recognition will need to account for this as the designs we measured had a range of ϕ from ≈10 to ≈35, which was only ≈7.5 × the uncertainty.

#### Section 9: Qualitative Trends

While our goal is to deconvolve the sources of cooperativity, several interesting qualitative trends in defectivity are observable within the families of design we examined.

The first qualitative trend was seen when rotating the scaffold sequence. In designs #1 to #10 we simultaneously rotated the sequence and moved the loop of 209 nucleotides of excess ssDNA. As shown in Fig. 5a, ϕ appears constant for rotations which keep the seam on the outside horizontal edge of the origami. However, when the excess ssDNA is brought inside the origami, or when it is on the vertical edges (therefore increasing loop entropy of many staples along a scaffold fold) ϕ increased dramatically. These results support the notion that sequence rotation does not meaningfully contribute to the thermodynamics of assembly[10,11]. These results also emphasize the negative steric hindrance effect associated with placing the excess ssDNA loop inside the structure.

The second trend, illustrated in Fig. 5b, concerns design symmetry. The overwhelming majority of 2D DNA origami designs detailed in the literature minimize the number of scaffold seams and distribute them symmetrically along the long axis of the design. We hypothesized that designs which broke this symmetry would increase or reduce yield in proportion to how evenly they distributed loop entropy penalties, *i.e.*, that a smoother free energy funnel should improve yield. If this were true, one would anticipate that a shifted seam should increase defectivity while an interdigitated one should reduce it.

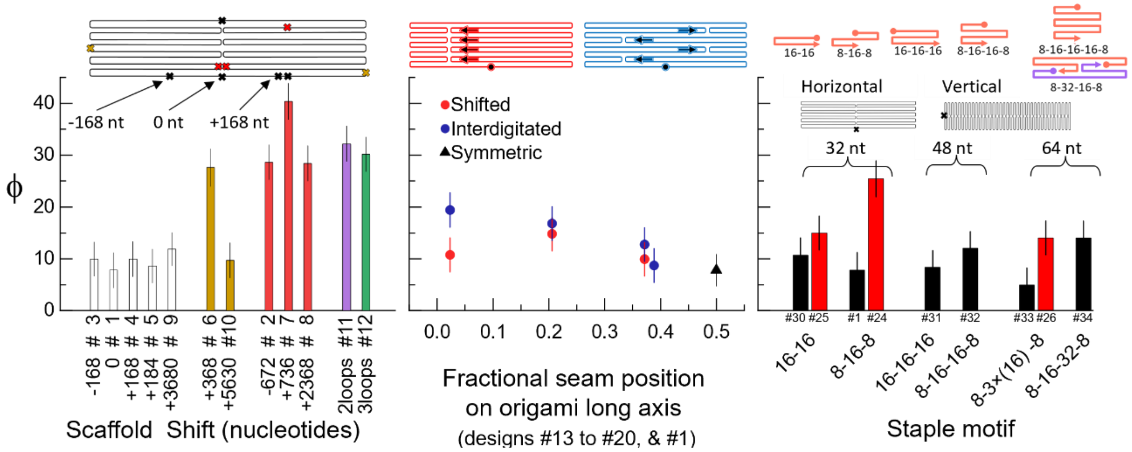

Fig. 5: a- Damage factor (ϕ) as the scaffold, and loop of excess ssDNA, are rotated through the design. The color of the data points corresponds to the colored x's in the above schematic. b- ϕ as the seam is pulled to one side of the design (red) or interdigitated (blue). c- variation in ϕ as the staple motif is modified for vertical routings (red) and horizontal routings (black), grouped by overall staple length. Error bars correspond to a single standard deviation and include anticipated staple synthesis yield.

However, as shown in Fig. 5b, any breaking of seam symmetry increased defectivity to a similar degree even though the two seam modifications shift entropic penalties in different ways. Shifting the entire seam (designs #13 to #16) converts mid-to-short-length folds into mid-to-long-distance folds without changing the number or length of the longest or shortest folds, see the following SI section. Interdigitating the seam (designs #17 to #20) consumes some mid-length and long folds to create a more even distribution of fold distances. In both cases, reducing seam symmetry moderately increased ϕ. It should be noted that designs #21 and #22 also modified the number of seams, and had an inconsistent effect on yield, with ϕ of 11 ± 3 and 18 ± 3 respectively.

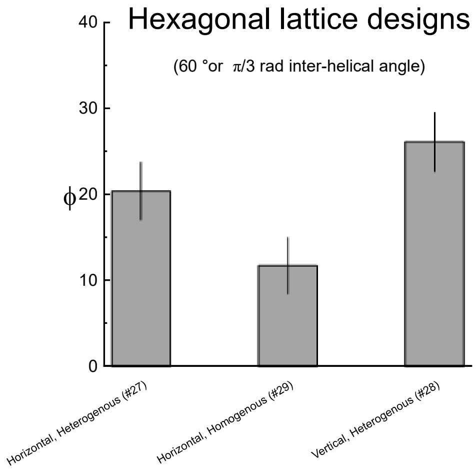

*Figure 6: Hexagonal lattice structure defectivity. Error on defectivity calculated as in the main text.*

Finally, it interesting that we are unable to observe a clear trend in ϕ as a function of staple motif, Fig. 5c. Given the importance of energetically heterogeneous staple domains for 3D structures and DNA bricks[12–14] we anticipated a clearer set of trends in yield. Further, we observed that 2D hexagonal lattice designs with more energetically heterogenous staple domains actually performed more poorly, see Figure 6. We suspect some difference between 2D and 3D structures is the cause of this discrepancy, particularly differences in the constraints on valid staple tiling. As many of these pools have varied staple lengths, the role of staple pool synthesis variation could also be exacerbated (see SI section 5).

Taken together, these results indicate that origami designers are again well served by their aesthetic biases. Keeping symmetric seams, keeping staples short, and using 8-16-8 motifs were effective approaches to reduce ϕ in the systems we examined.

#### Section 10: Seam symmetry designs

The designs which break seam symmetry provide a useful frame of reference for our result that the skew of the entropy penalty distribution of folding events appears to be negatively correlated with defectivity and place the results in the previous section, Fig. 5, in context.

Before showing the loop distributions for each design, we begin by comparing two more straightforward examples. Figure 7 shows the routing plots and square plots for the default horizontal (#1) and vertical (#24) origami at the top to give an intuitive sense of the distribution of figures. At the bottom it shows two sets of histogram comparisons. In each comparison the top panel is the histogram of folding distances for each case. The bottom panel is either the loop entropy distribution or the blocking probability. Figure 7 illustrates that the loop entropy penalty takes the fold distance distribution and flips it (larger fold, larger negative penalty) and compresses long folds together. In contrast the blocking probability does not flip the order of the distribution, and compresses folds shorter than at least several hundred bases.

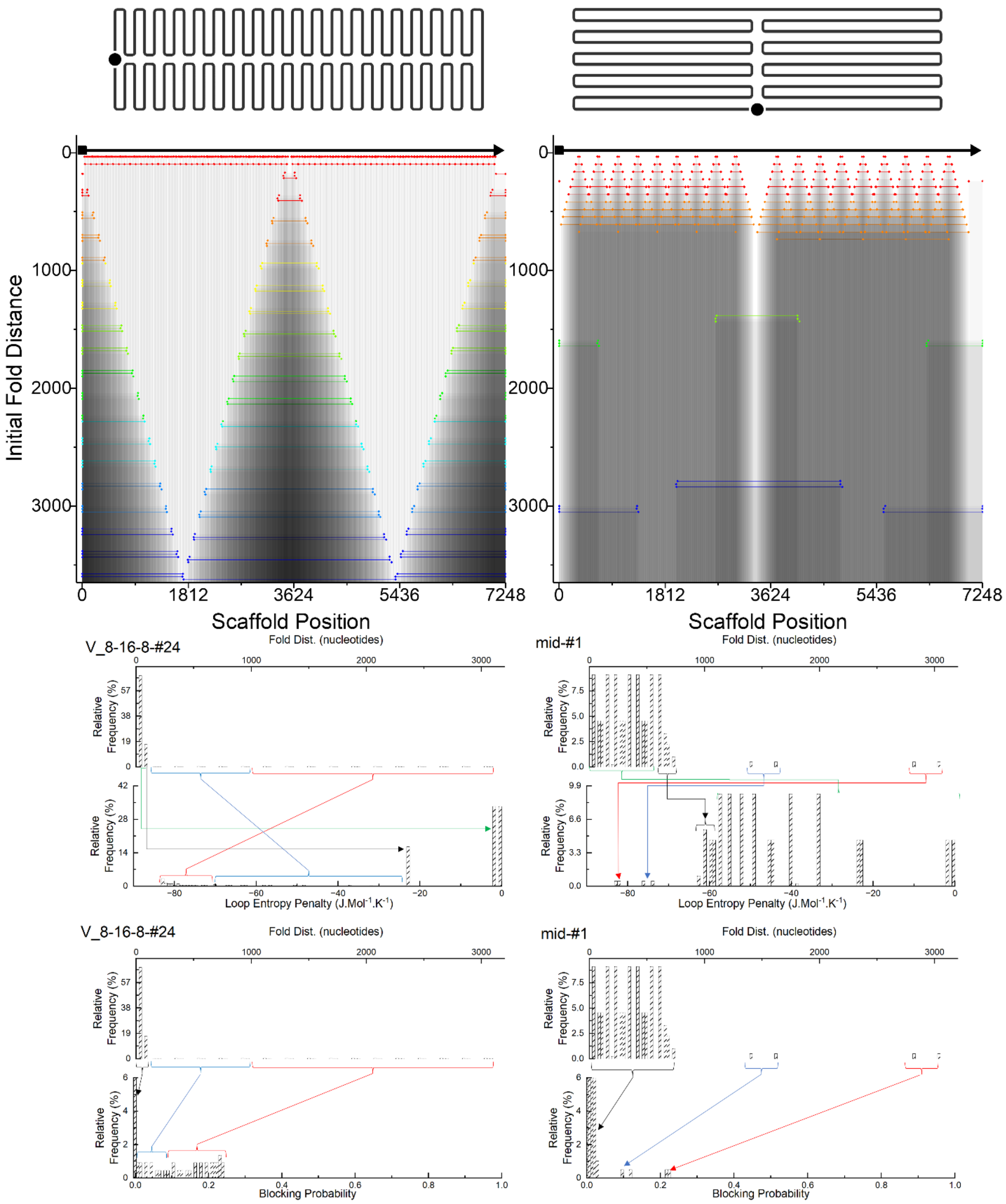

*Figure 7: Relationship between distribution of folding distances and the distributions of loop penalties and blocking probabilities for two example designs (#1 and #24). Transforming the fold distance distribution into loop penalties compresses the difference between long folds and expands the difference between short folds. By contrast the blocking probability calculation compresses the difference between all folds, and any fold less than approximately 1 000 nucleotides has a minimal chance of forming the blocked state at our low staple excess (5×).*

With this relationship between fold distances in mind, we then present Table 14 below, which shows the routing patterns (left column) for the designs with shifted seams (#13 to #16), and interdigitated seams (#17 to #20), and the histogram of fold distances. As is visible from these plots, altering seam symmetry appears to increase the number of mid-length folds which can block folding, without necessarily creating a more even distribution of entropy penalties.

*Table 14: Progression of fold distance distributions and fold $T_m$ distributions for designs which incrementally modify the origami seam*

Design #13 Horizontal – shifted (16 nt)

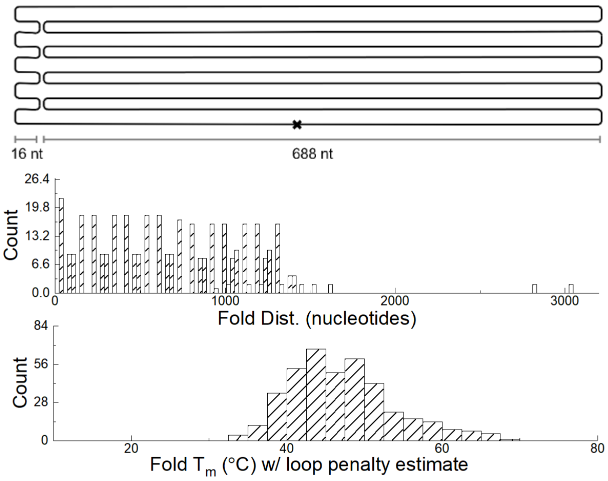

Design #14 Horizontal – Shifted (144 nt)

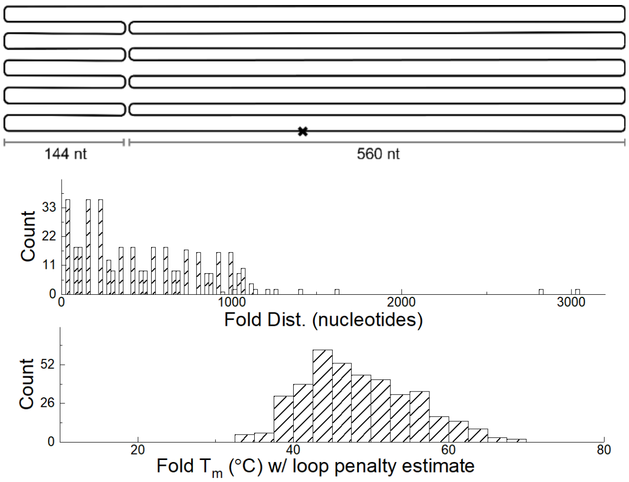

Design #15 Horizontal – Shifted (260 nt)

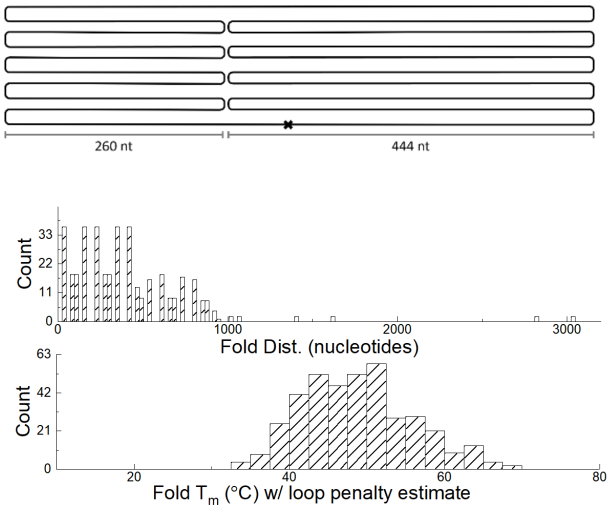

Design #16 Horizontal – shift (165 nt)

w/ design flaw

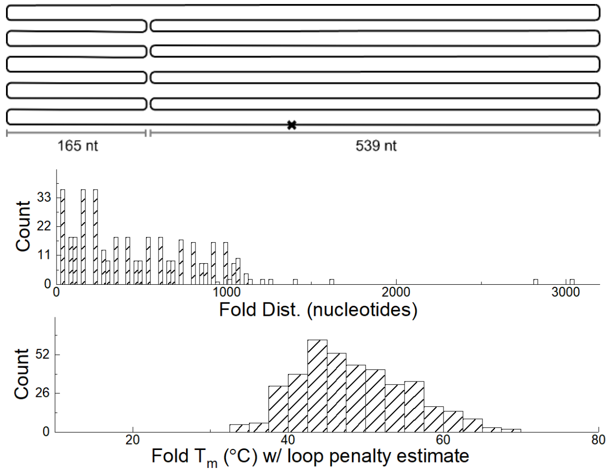

Design #17 – inter-digitated (16 nt)

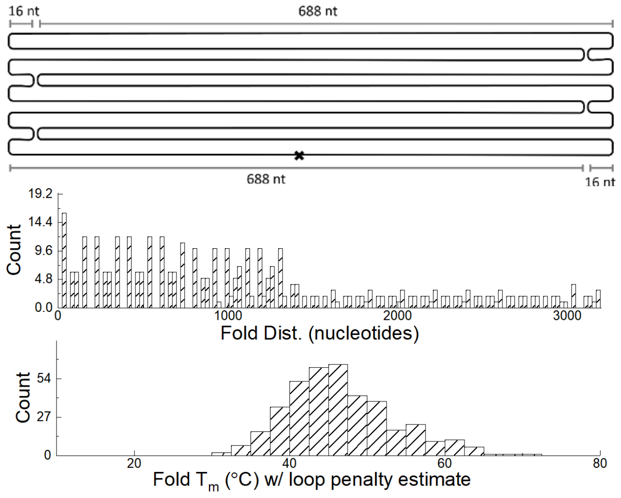

Design #18 – inter-digitated (144 nt)

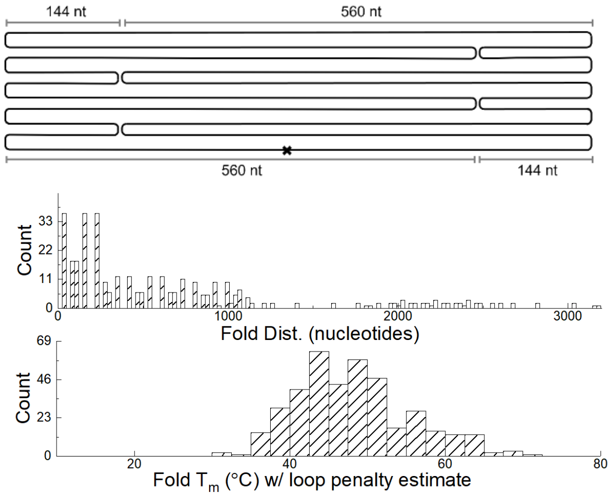

Design #19 – Interdigitated, 260 nt

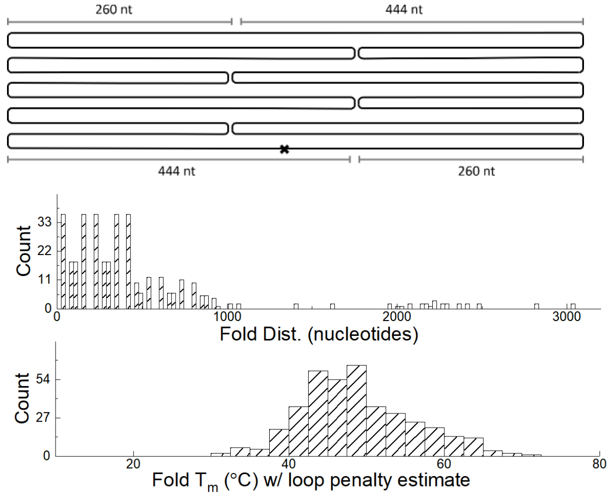

Design #20 – Interdigitated, 272 nt

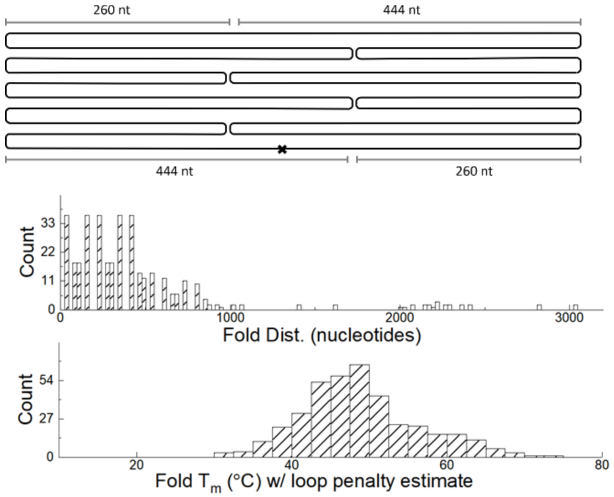

#### Section 11: cadnano modification for topological information

For this work, we used a modified version of cadnano[15] to obtain topological information for further data processing, built off of Nash's cadnanovis tool on nanohub[16]. We are happy to provide the code on request, but note that it was not made to be built on further, and that it is limited to providing this information from designs one at a time, and that each of our designs was created manually. In short, as our final heuristic correlating defectivity to design requires normalization across many designs, we believe several more advances are necessary before automated design is possible.

The cadnanovis tool is designed to create the circle plots discussed in SI section 10 and presented for each design in SI section 17. This tool exports topology information, specifically the scaffold base # for each base of each staple.

We further modified this tool to include custom classes for staples, domains, and folds. Staples were comprised of folds which were comprised of domains, each with indices pointing back up to the parent object. Each object retained the nucleotide sequences, corresponding scaffold indices, and initial looping distances associated with it. We used these classes to populate hierarchical lists of staples, folds, and domains, which we then used to calculate fold distance dependent calculations and from these calculated our metrics.

As cadnano's data structure is, reasonably, built around its visualization of the design, it is worth noting that populating topological information must be cognizant of a variety of edge cases. In particular, we performed manual cross checking of topological information with small designs comprising <10 staples, paying close attention to the seams and edges where the scaffold crosses over between helices.

#### Section 12: Nearest neighbor calculations for predicted $T_m$

The nearest neighbor calculations were performed as per Santa Lucia et al.[17] assuming a 0.375 nMol.$L^{-1}$ concentration of scaffold and a 5 × excess of staples to scaffold and 1 mol.$L^{-1}$ NaCl for simplicity. The loop entropy penalty of closing the scaffold circle into a figure eight was calculated as per our previous work and described in Eq. 1 of this SI[4].

#### Section 13: Intercalating dye melt data

The intercalating dye melt and anneal data were performed in a 96 well plate in temperature controlled fluorimeter (RT-PCR) unit. Each design had four replicate wells, each of which 50 µL in volume. All had a final M13-MP18 concentration of 7.5 nMol.$L^{-1}$ (nM) and a staple concentration of 10× the scaffold (75 nMol.$L^{-1}$). The buffer used was 1× TAE (40 mMol.$L^{-1}$ Tris. pH'd to 7 with acetic acid) with 12.5 mMol.$L^{-1}$ $MgCl_2$. This buffer was supplemented with 13.3 µMol.$L^{-1}$ SYBR Green intercalating dye. This amounts to approximately 1 dye per 4 bases of origami dsDNA, or 1 dye per 45 bases of ssDNA.

This dye concentration is much higher than used in some other studies[18]. They were able to use much lower concentration of dye in part because their anneals ran on the order of days rather than hours. This is critical as the dyes will also bind to ssDNA in the excess staples.

insight. This is because the multistate dye, ssDNA, and dsDNA is particularly complex to unravel. As such, we do not use the more stringent methods, cacodylate buffer to preserve pH with temperature and affine transformations for data analysis that we might otherwise use.

Both melt and anneal ramps were performed in 0.2 °C steps, but due to a program template error the melt curves had a 15 second dwell time while the anneal curves had a 20 sec dwell time at each step. The anneal curves stepped down from 80 °C to 13 °C, after a short 30 second denaturation at 95 °C. The first melt plate ran from 10 °C to 80 °C, but slightly truncated the final edge of some melt curves, so the second plate was ran from 20 °C to 90 °C. We chose not to regather this data as the difference in dwell time is expected to result in minimal difference (A separate set of testing on only a few origami showed near perfect overlap between melt curves and between anneal curves over the course of a 12 hr and 2 hr anneal). Additionally, the melt and anneal curves presented here could not be gathered at identical conditions to our accelerated anneal protocol, so their use is limited to a comparative metric of thermal stability between designs.

Melt-anneal hysteresis was identified by taking all data points in the temperature range and subtracting the anneal intensity counts from the melt intensity counts and multiplying the difference by 0.2 °C, which was the temperature step size. If the difference at high temperature was larger than the difference at low temperatures where the systems should be identical, it was assumed to linearly decrease to a negligible difference at 95 °C. The contribution of these assumptions was consistently less than 5 % of the hysteresis area. This was necessary as a minority of the melt curves were not fully melted at 80 °C. Attempts to reduce concentration resulted in suboptimal derivative curves. Future studies using more sensitive equipment and careful variation of annealing and melt rates may yield useful information.

The hysteresis calculated by area and the hysteresis calculated by derivative peak position were in reasonable agreement, see Figure 8.

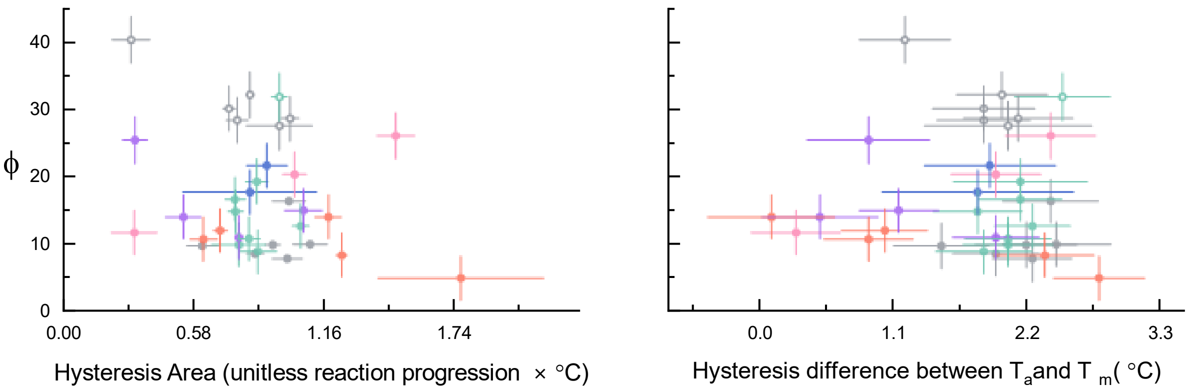

*Figure 8. Comparison of hysteresis calculations. (left) the area integrated between melt and anneal curves. (right) difference between average Tm and Ta values. Y-error bars calculated as elsewhere, X-error bars (left) are a single standard deviation between the four replicate wells of each design and (right) are the addition in quadrature of the standard deviation between the four replicate $T_m$ and $T_a$ values for each design.*

For completeness we plot the derivative peak $T_m$ and $T_a$ against each other in Figure 9. We note that in all cases the $T_m$ is higher as one might anticipate.

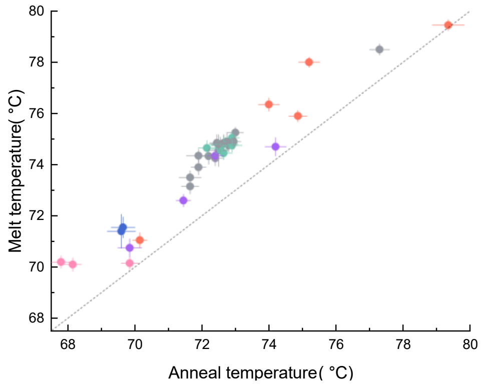

*Figure 9: Plot of derivative peak calculated $T_m$ and $T_a$ against each other to illustrate melt anneal hysteresis. The dotted line indicates positions where $T_m$ would equal $T_a$. Error bars represent a single standard deviation and datapoint color matches design family as elsewhere in the text.*

#### Section 14: Comparison to literature metrics

While in the main text we have constrained ourself to examining cooperativity across the broad network of folding events in origami, some other metrics or ways to evaluate an origami design have been suggested in the literature and in informal conversations. This SI section will briefly address them.

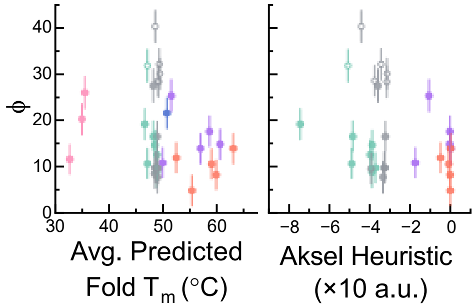

*Figure 10. Comparison of the heuristic published by Aksel et al. with the Average Fold Predicted $T_m$, which is most similar value we calculate here. Note: the webtool used to calculate the Aksel heuristic crashed when presented with planar hexagonal lattice designs (pink designs). Data point color legend and error bars are identical to all other plots in this work.*

Figure 10 compares the average of the predicted fold $T_m$ for each design to the heuristic developed by the Akesel et al[19]. Their heuristic takes the same predicted fold $T_m$ that we use and propagates it through the design with an application of graph theory to allow for iterative optimization. Their webtool had some difficulties with our planar hexagonal lattice structures, but otherwise had a similar trend to the average of fold $T_m$s across a design.

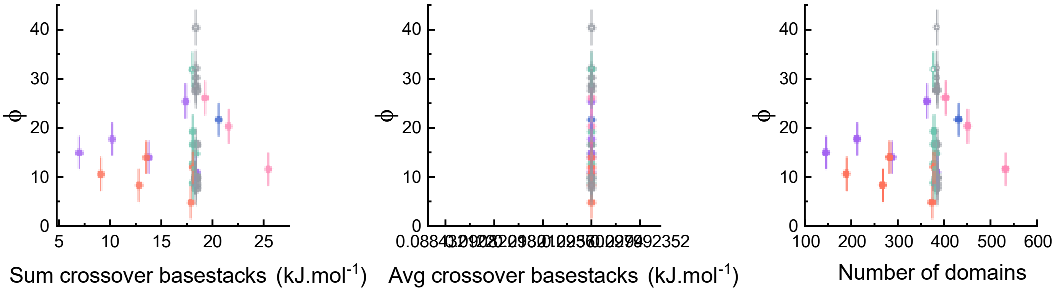

*Figure 11. Comparison of defectivity against the sum of crossover base stacks, the average of all crossover base stacks, and the number of domains. Note that our algorithm did not distinguish between single and double crossovers. Error bars and data color scheme are the same as in all other plots.*

Another potential effect correlated with defectivity discussed in the literature is a hypothesis proposed by Cumberworth et al[20], that high GC content at crossovers might act as a kinetic roadblock to assembly. This is slightly different from the domain base stacking metric used for positive interdomain cooperativity as it only sums the enthalpic contributions at positions where the staples cross over between scaffold positions. Figure 11 shows that the variation in base stacking energy is due almost entirely to the number of crossovers, with variation in GC base content being so relatively small so as to be on the same order as rounding errors. This is supported by Fig. 5, in which the 16-16 motifs do not have noticeably lower defectivity than the 8-16-8 motifs. If double crossovers with high GC content formed meaningful barriers to assembly one would anticipate the 16-16 motif designs to have much lower defectivity as they inherently have only single crossovers.

#### Section 16: Supplementary bibliography

1. Majikes, J. M., Nash, J. A. & LaBean, T. H. Competitive annealing of multiple DNA origami: formation of chimeric origami. *New J Phys* **18**, 115001 (2016).

2. Dunn, K. E. *et al.* Guiding the folding pathway of DNA origami. *Nature* (2015) doi:10.1038/nature14860.

3. Virtanen, P. *et al.* SciPy 1.0: fundamental algorithms for scientific computing in Python. *Nat Methods* **17**, (2020).

4. Majikes, J. M. *et al.* Revealing thermodynamics of DNA origami folding via affine transformations. *Nucleic Acids Res* 1–13 (2020) doi:10.1093/nar/gkaa283.

5. Majikes, J., Patrone, P., Kearsley, A., Zwolak, M. & Liddle, J. Failure Mechanisms in DNA Self-Assembly: Barriers to Single-Fold Yield. *ACS Nano* **15**, 3284–3294.

6. Tibshirani, R. Regression shrinkage and selection via the lasso: A retrospective. *J R Stat Soc Series B Stat Methodol* **73**, (2011).

7. Friedman, J., Hastie, T. & Tibshirani, R. Regularization paths for generalized linear models via coordinate descent. *J Stat Softw* **33**, (2010).

8. Team, R. C. R Core Team 2023 R: A language and environment for statistical computing. R foundation for statistical computing. https://www.R-project.org/. *R Foundation for Statistical Computing* (2023).

9. Cavaluzzi, M. J. & Borer, P. N. Revised UV extinction coefficients for nucleoside-5'-monophosphates and unpaired DNA and RNA. *Nucleic Acids Res* **32**, (2004).

10. Dannenberg, F., Dunn, K. E., Bath, J., Turberfield, A. J. & Ouldridge, T. E. Modelling DNA Origami Self-Assembly at the Domain Level. *J Chem Phys* **143**, 165102 (2015).

11. Arbona, J.-M., Aimé, J.-P. & Elezgaray, J. Cooperativity in the annealing of DNA origamis. *J Chem Phys* **138**, 015105 (2013).

12. Jacobs, W. M. & Frenkel, D. Self-Assembly of Structures with Addressable Complexity. *J Am Chem Soc* **138**, 2457–2467 (2016).

13. Jacobs, W. M., Reinhardt, A. & Frenkel, D. Rational design of self-assembly pathways for complex multicomponent structures. *Proc Natl Acad Sci U S A* **112**, 6313–8 (2015).

14. Ke, Y., Bellot, G., Voigt, N. V., Fradkov, E. & Shih, W. M. Two design strategies for enhancement of multilayer–DNA-origami folding: underwinding for specific intercalator rescue and staple-break positioning. *Chem Sci* **3**, 2587 (2012).

15. Douglas, S. M. *et al.* Rapid prototyping of 3D DNA-origami shapes with caDNAno. *Nucleic Acids Res* **37**, 5001–5006 (2009).

16. Nash, J. A. DNA Origami Visualization Tools. Preprint at (2017).

17. SantaLucia, J. & Hicks, D. The thermodynamics of DNA structural motifs. *Annu Rev Biophys Biomol Struct* **33**, 415–440 (2004).

18. Sobczak, J.-P. J., Martin, T. G., Gerling, T. & Dietz, H. Rapid folding of DNA into nanoscale shapes at constant temperature. *Science* **338**, 1458–61 (2012).

19. Aksel, T., Navarro, E. J., Fong, N. & Douglas, S. M. Design principles for accurate folding of DNA origami. *Proceedings of the National Academy of Sciences* **121**, e2406769121 (2024).

20. Cumberworth, A., Frenkel, D. & Reinhardt, A. Simulations of DNA-Origami Self-Assembly Reveal Design-Dependent Nucleation Barriers. *Nano Lett* **22**, (2022).

#### Section 17: Design information sheets

The below pages contain individual information sheets for each design including circle and square plots as well as distributions of predicted Tm and loop distances.

### Design #1 Horizontal Middle Break Origami

**ϕ – 7.8 ± 3.4**

**W=52±6 %; D = 19±5 %; R= 24±4 %, C= 5 ±2 %**

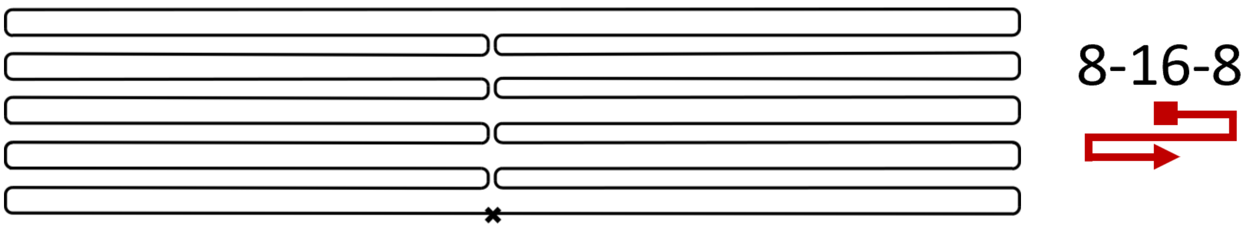

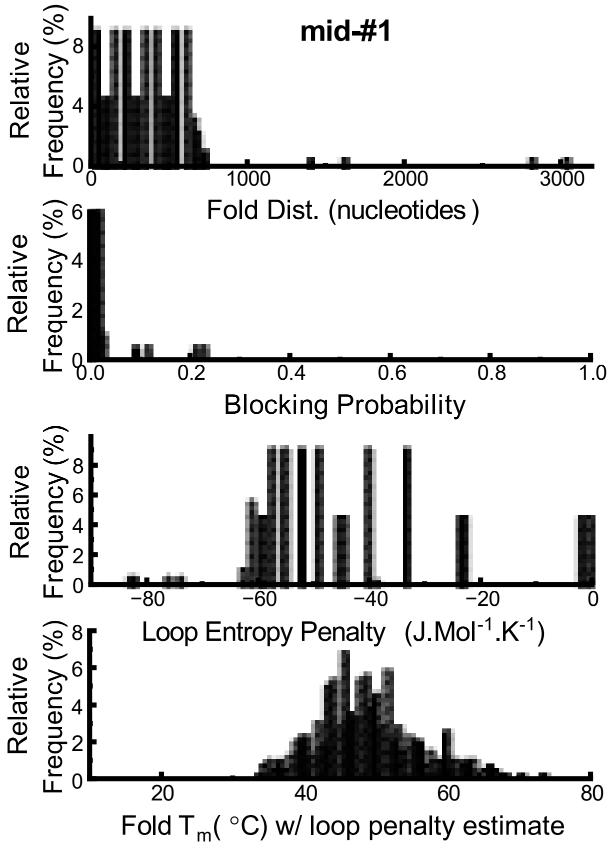

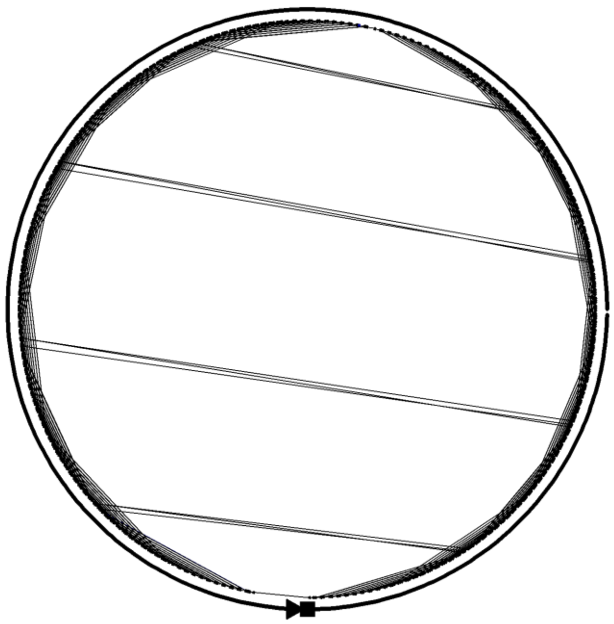

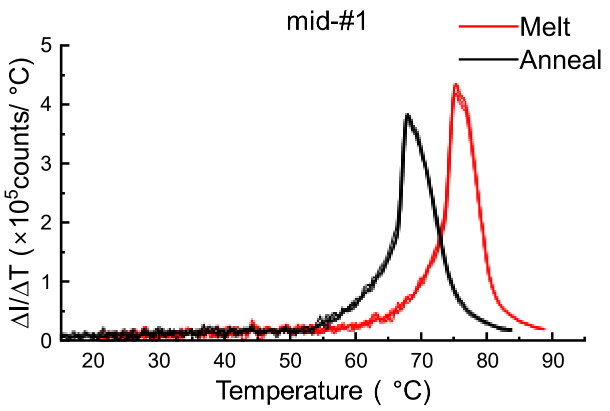

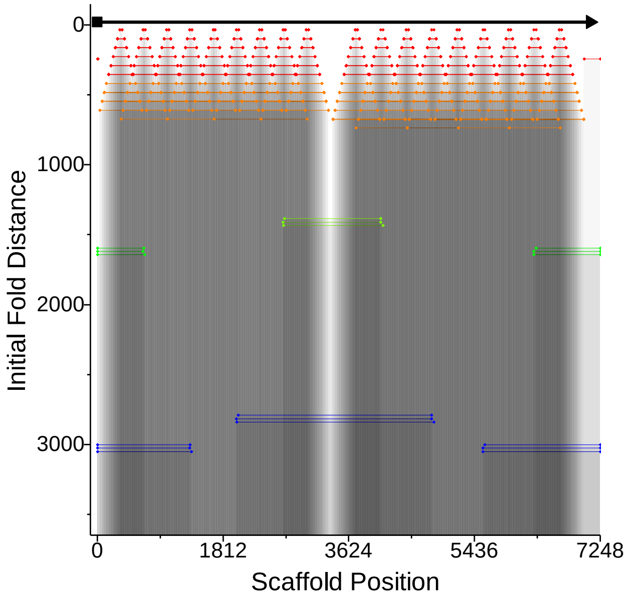

#### Design #2 Horizontal neg. 672 nucleotides

**ϕ – 28.7 ± 3.0**

**W=7±2 %; D = 12±2 %; R= 35±3 %, C= 45±3 %**

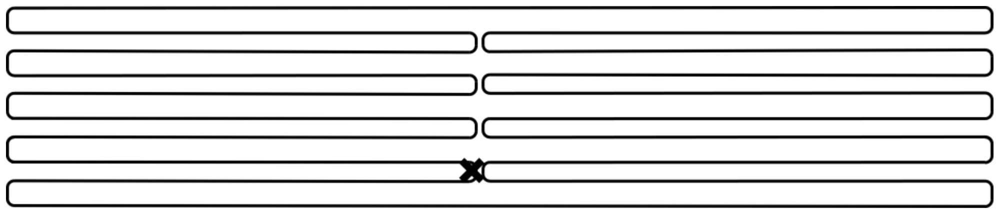

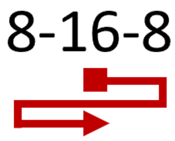

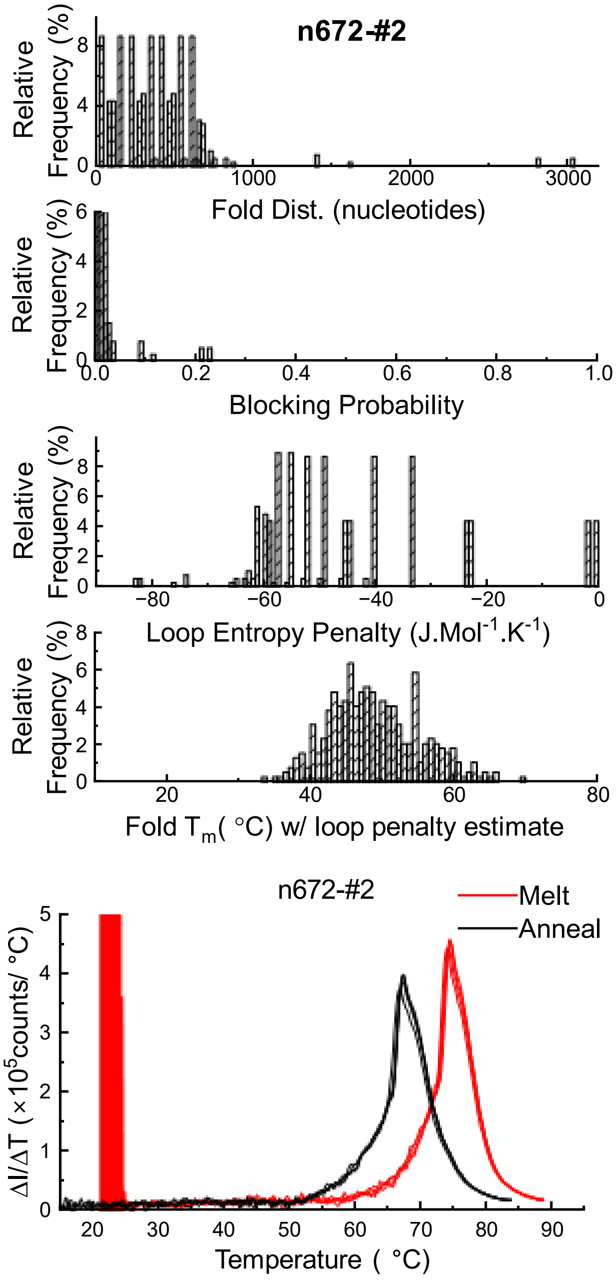

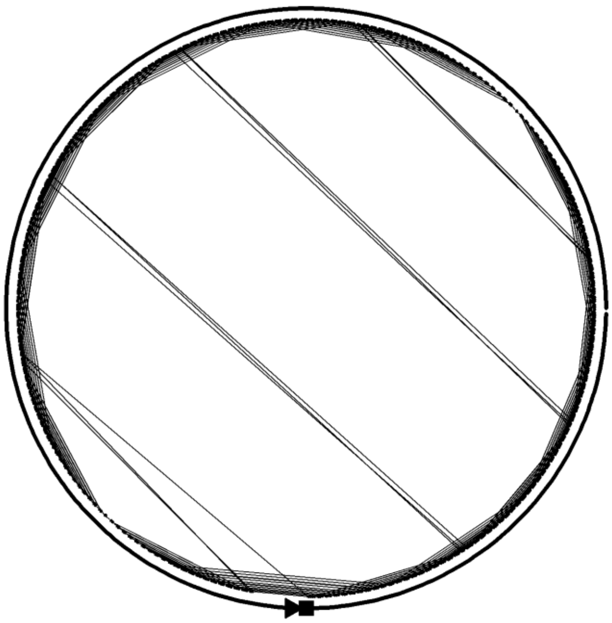

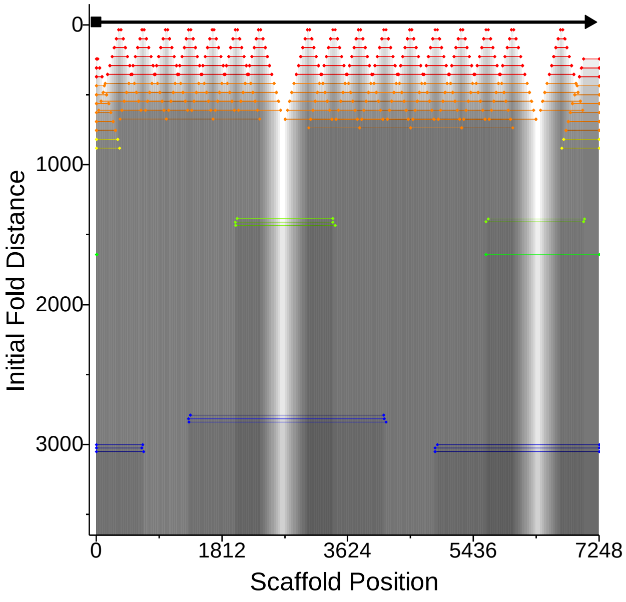

### Design # 3 Horizontal neg. 168 nucleotides

**ϕ – 10 ± 3.3**

**W=15±3 %; D = 48±4 %; R= 32±3 %, C= 5±1 %**

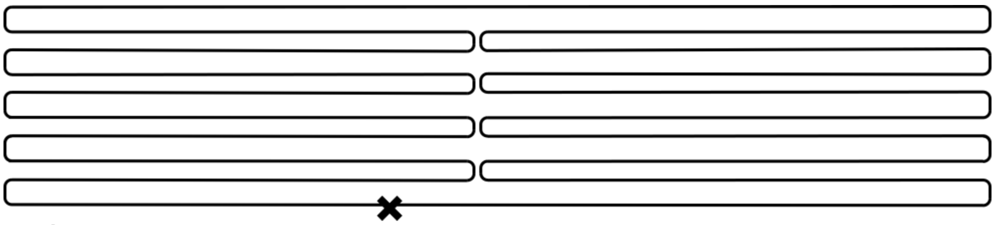

8-16-8

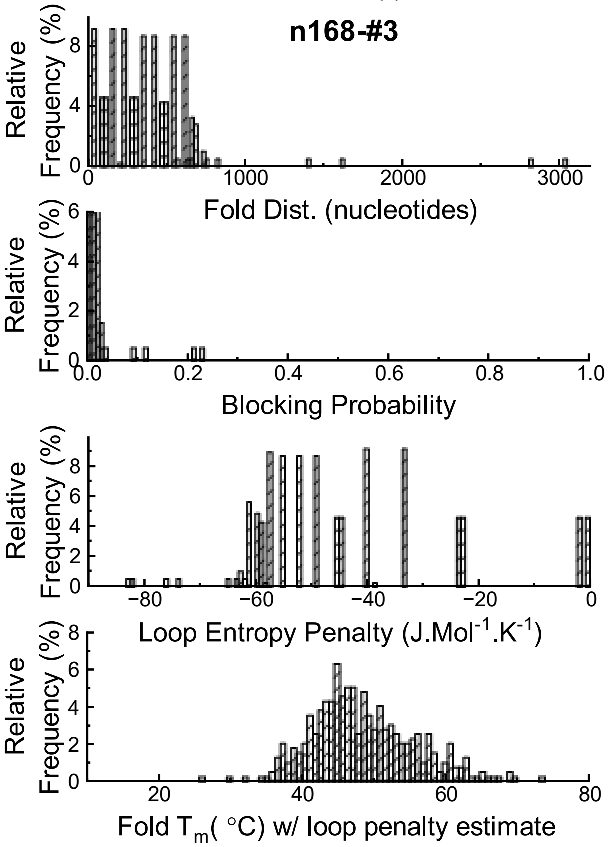

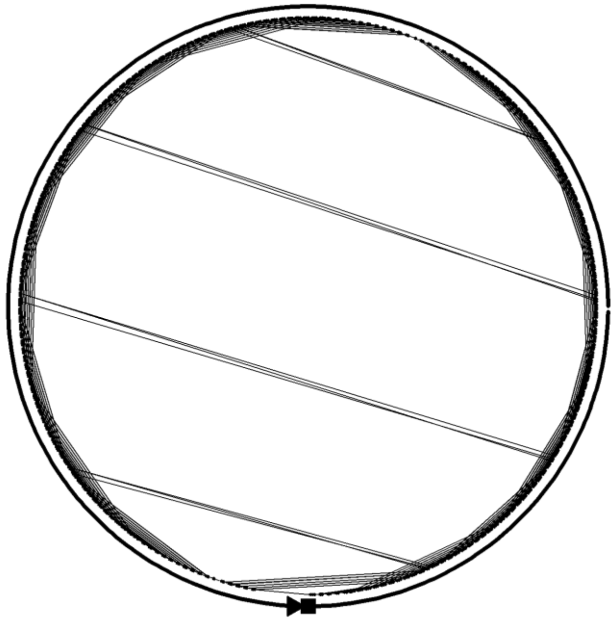

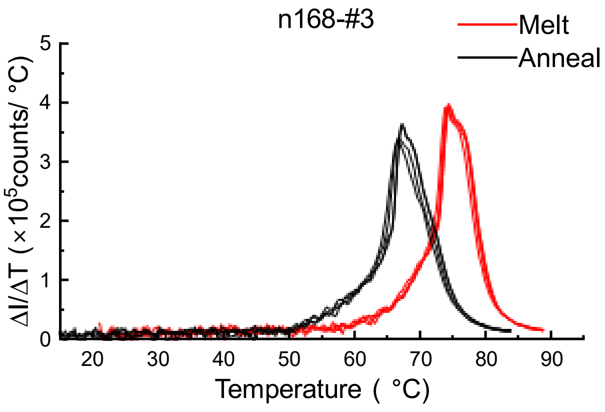

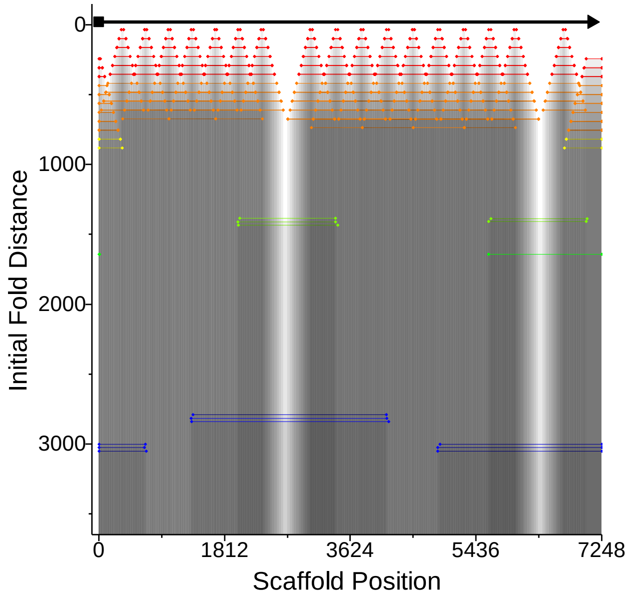

### Design # 4 Horizontal pos. 168 nucleotides

**ϕ – 9.9 ± 3.4**

**W=18±4 %; D = 48±5 %; R= 27±4 %, C= 7 ±2 %**

8-16-8

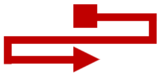

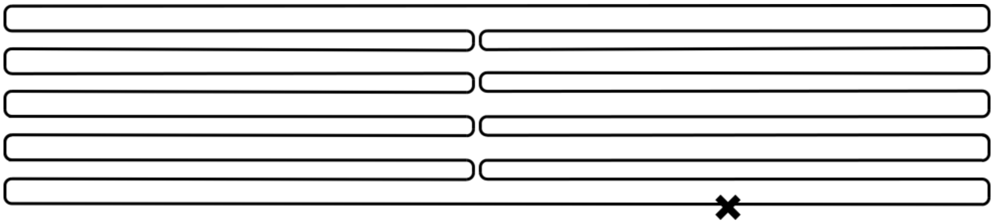

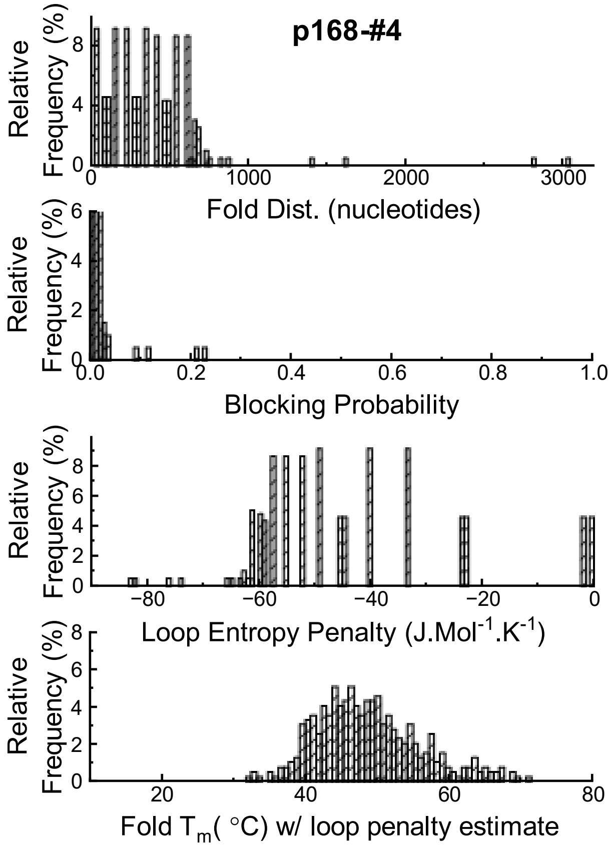

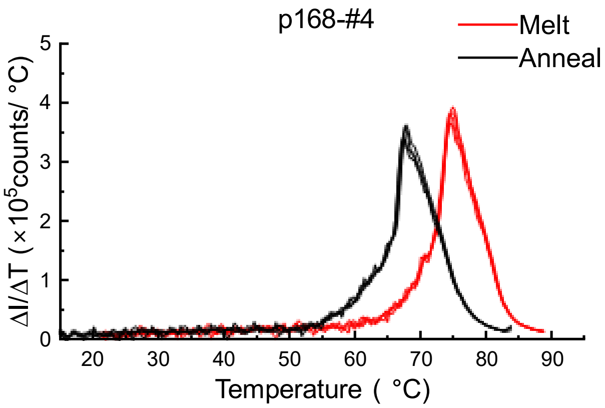

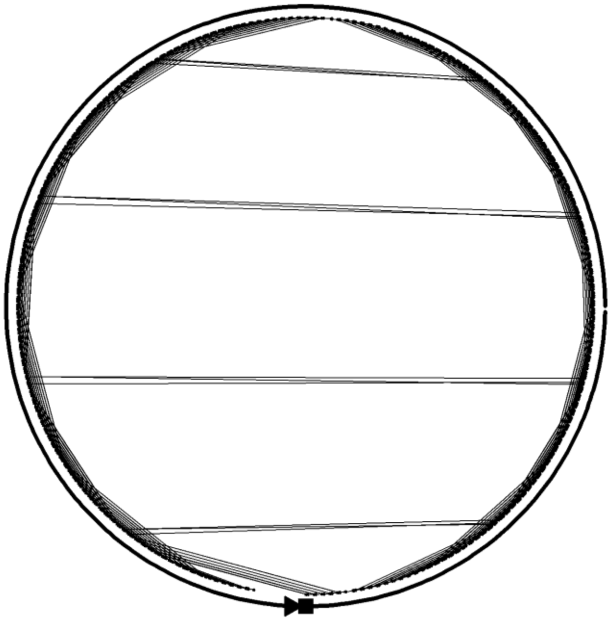

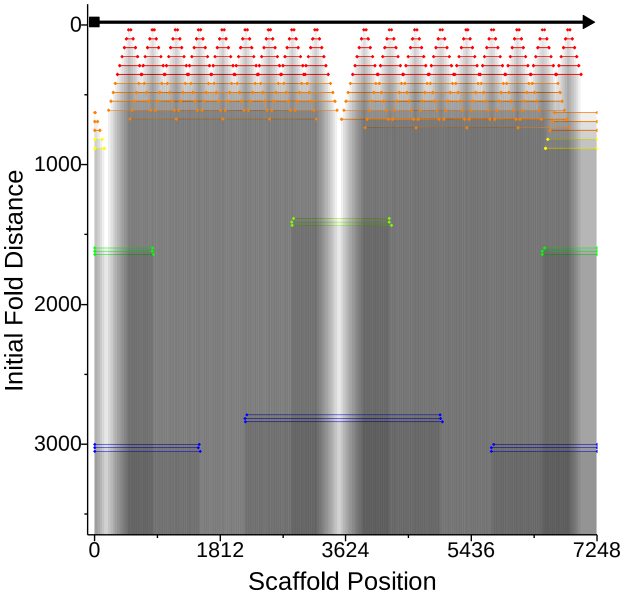

### Design # 5 Horizontal pos. 184 nucleotides

**ϕ – 8.6 ± 3.3**

**W=33±3 %; D= 34±3 %; R= 28±3 %, C= 5 ±1 %**

8-16-8

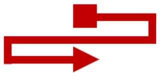

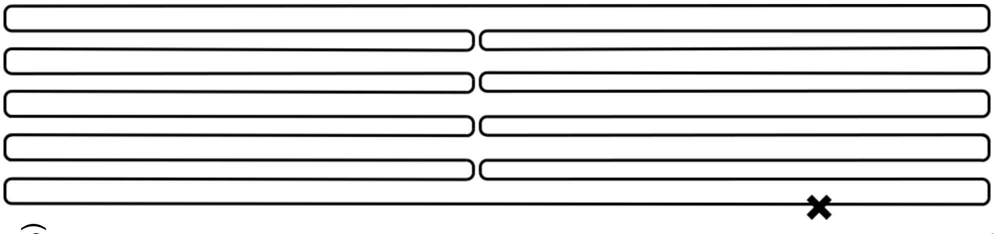

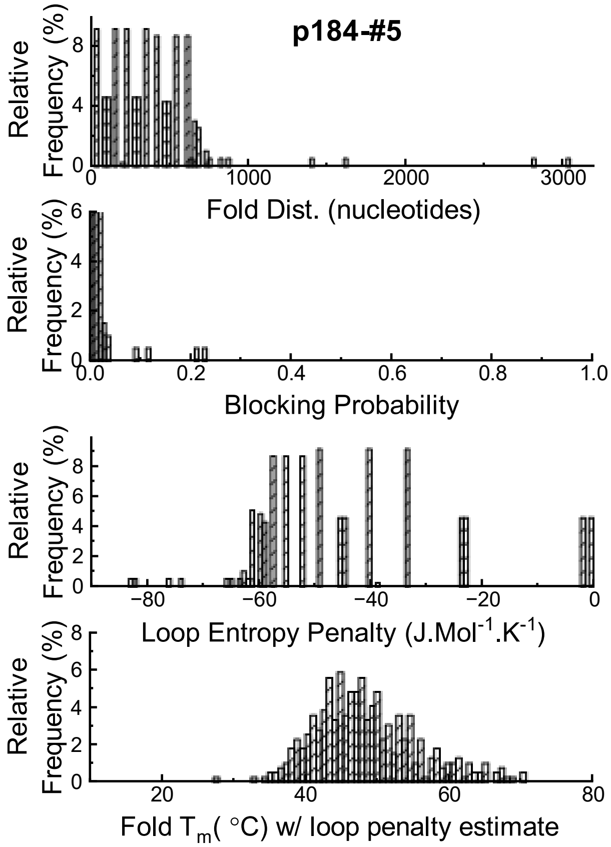

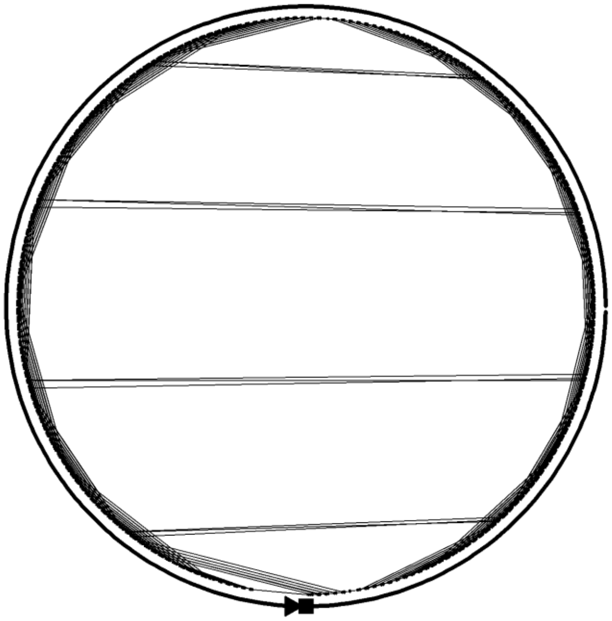

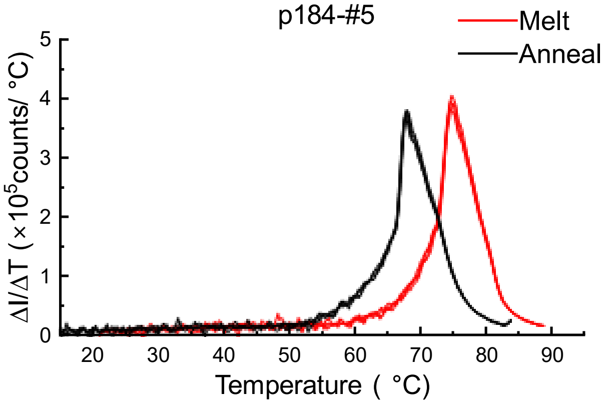

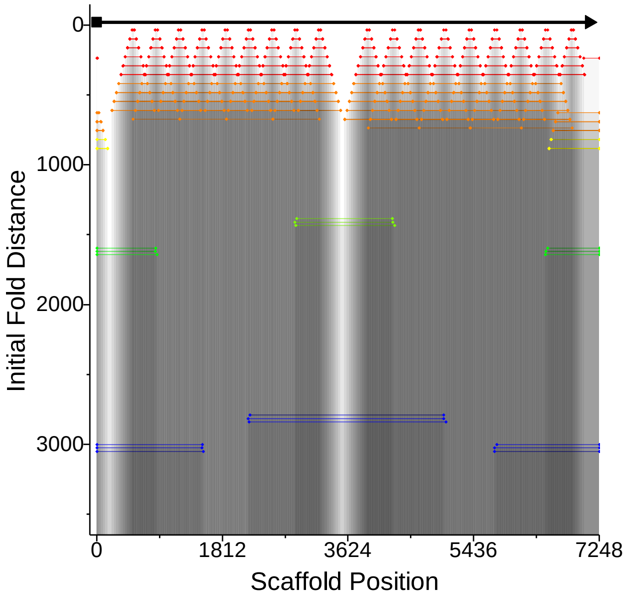

### Design #6 Horizontal pos. 368 nucleotides

**ϕ – 27.6 ± 3.6**

**W=2±2 %; D = 13±3 %; R= 45±5 %, C= 40 ±4 %**

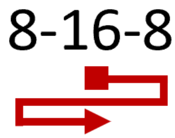

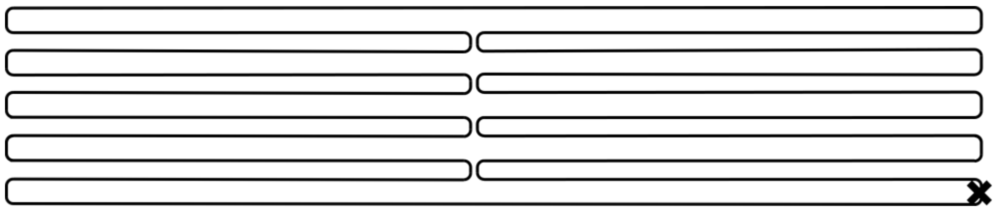

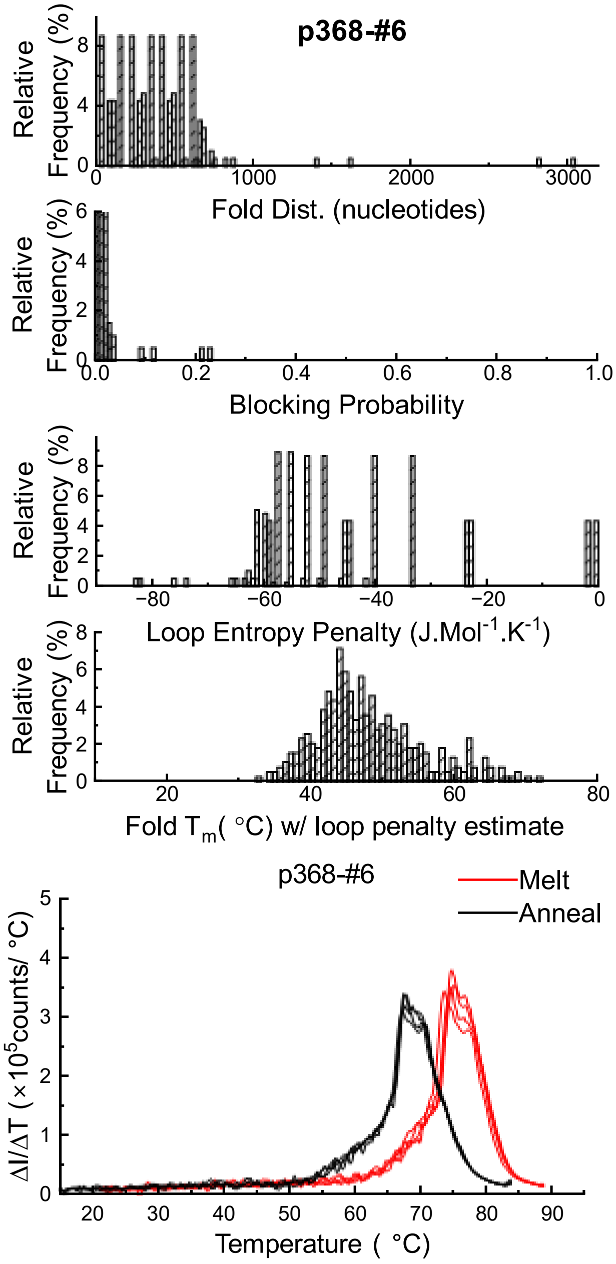

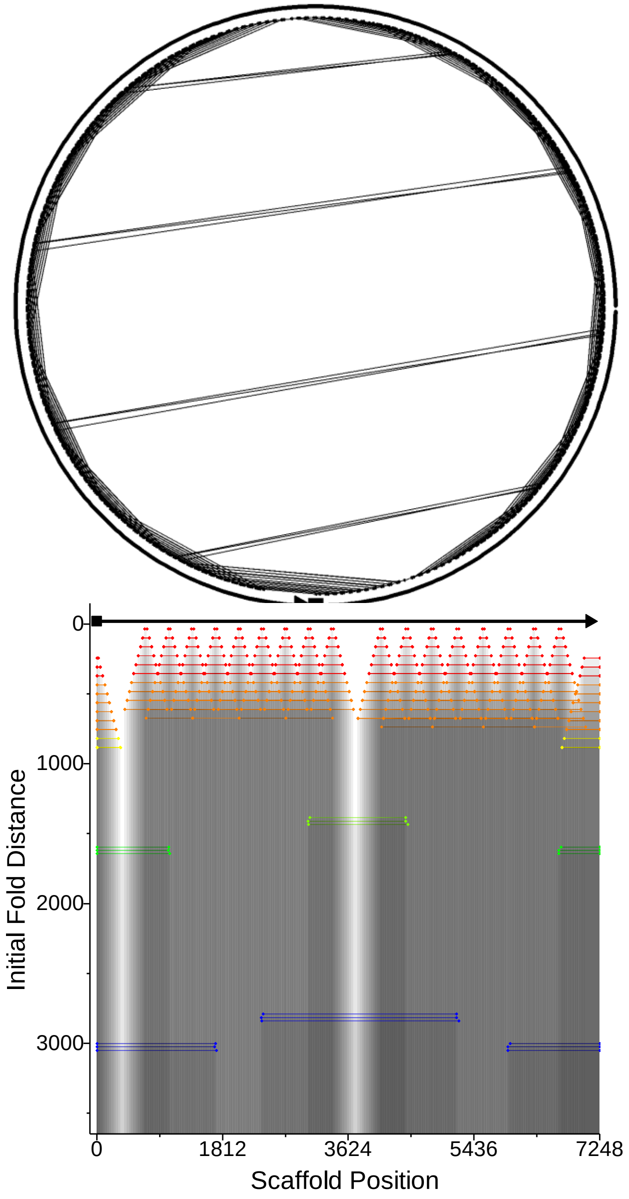

### Design #7 Horizontal pos. 736 nucleotides

**ϕ – 40.4 ± 3.5**

**W=10±2 %; D = 3±2 %; R= 9±52 %, C= 78 ±3 %**

8-16-8

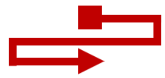

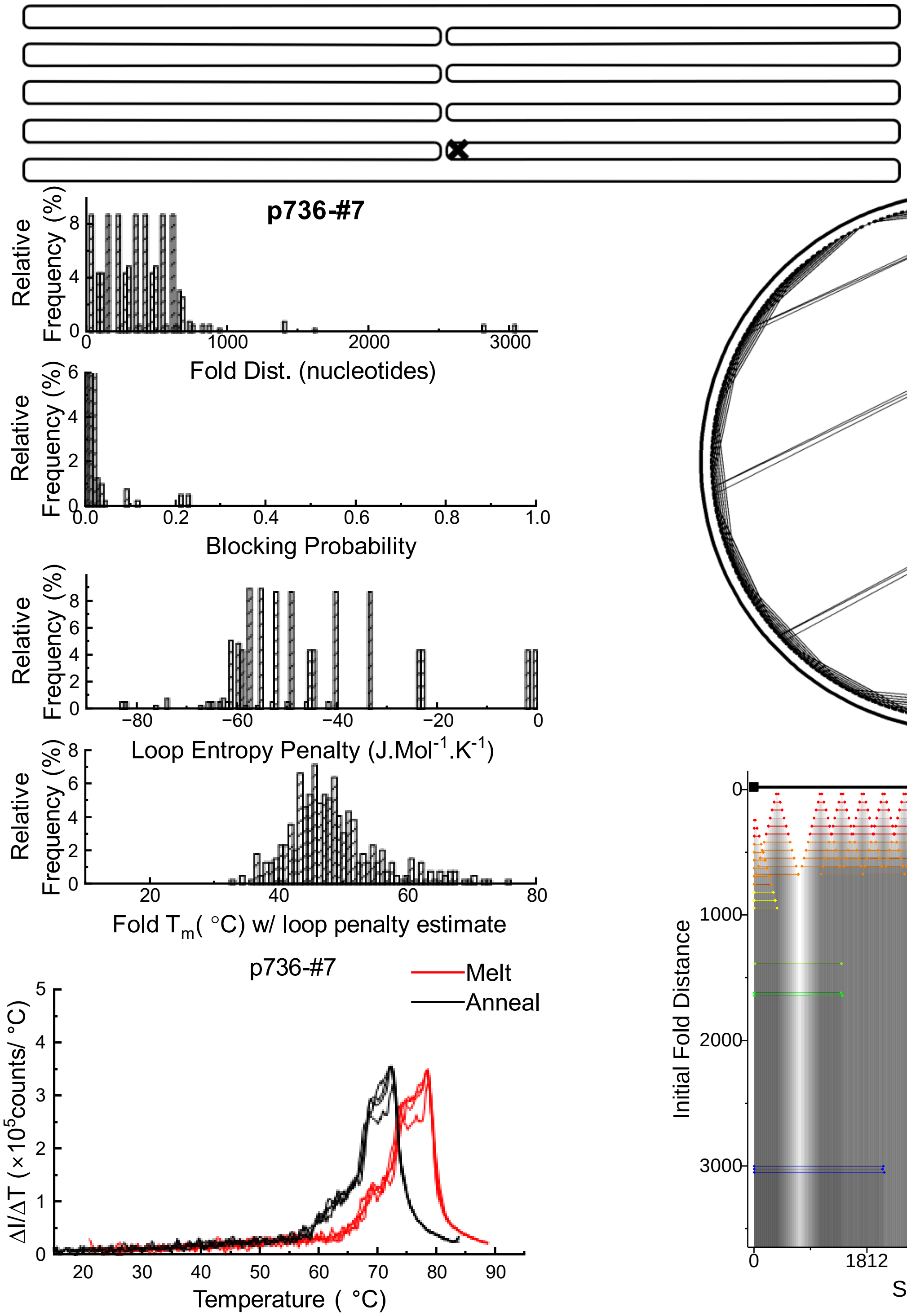

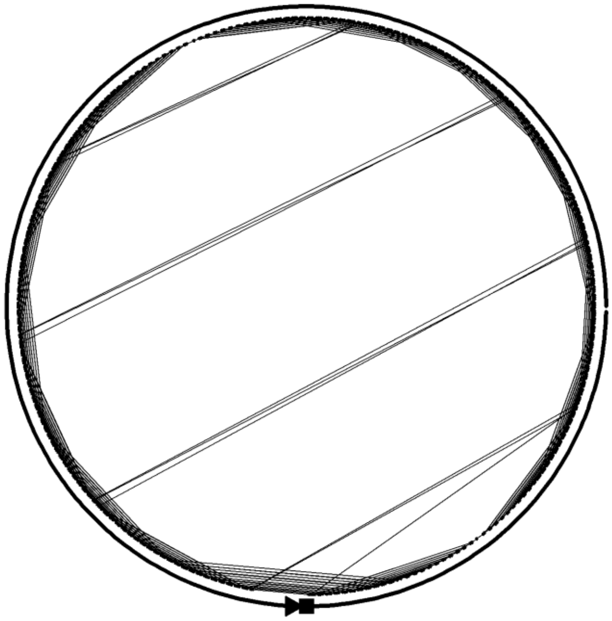

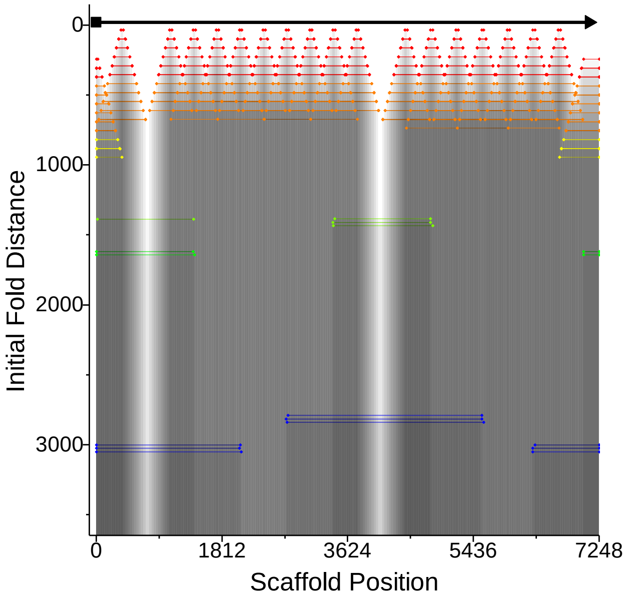

### Design #8 Horizontal pos. 2368 nucleotides

**ϕ −28.4 ± 3.4**

**W=1±1 %; D = 5±2 %; R= 54±3 %, C= 40 ±3 %**

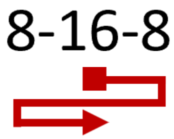

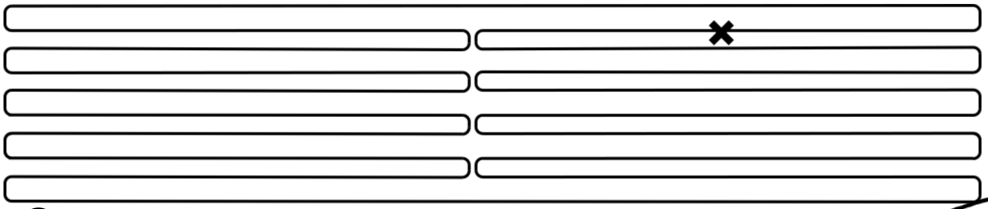

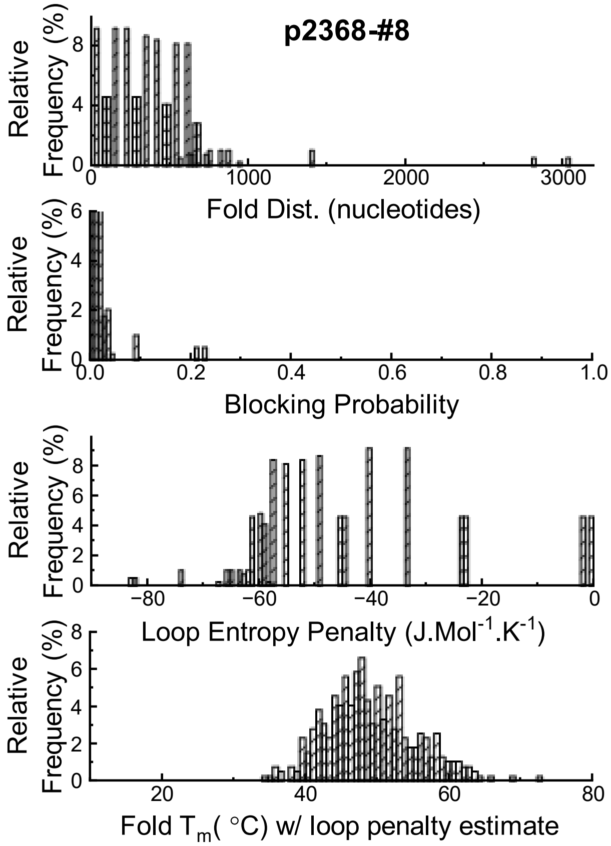

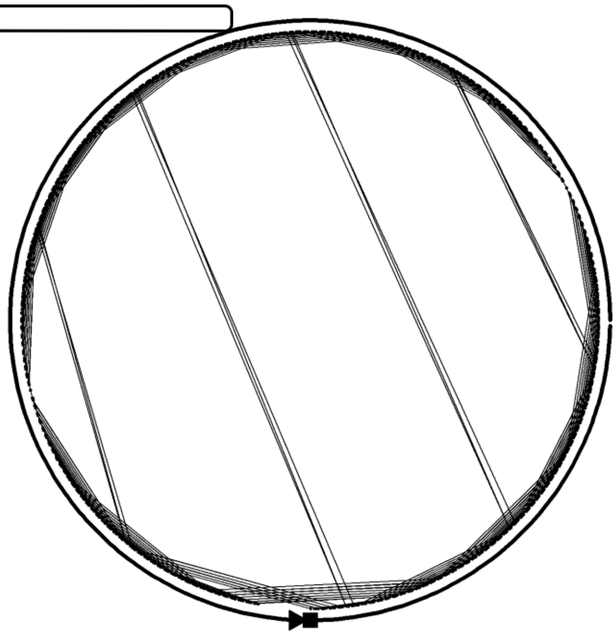

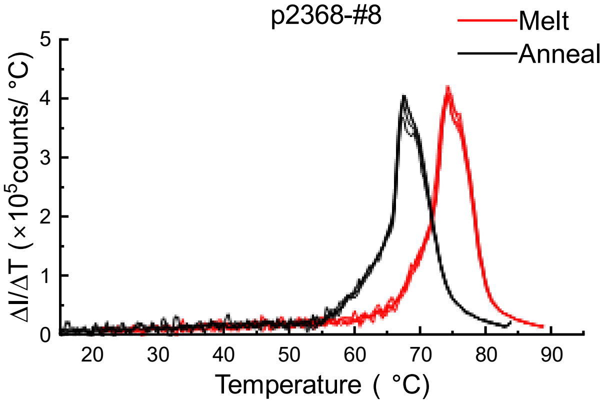

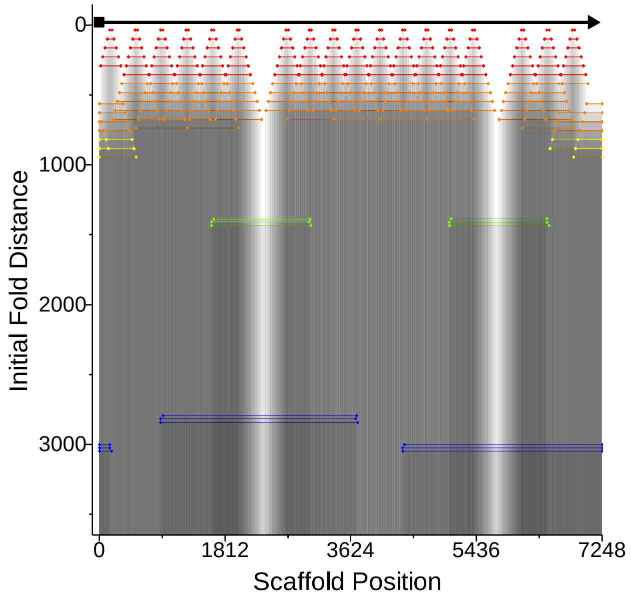

### Design # 9 Horizontal pos. 3680 nucleotides

**(all pools)**

**ϕ – 16.7 ± 3.2 for each (± 1.6 between them)**

**W=9±1 %; D = 17±1 %; R= 61±2 %, C= 13 ±1 %**

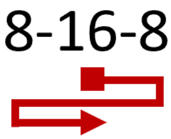

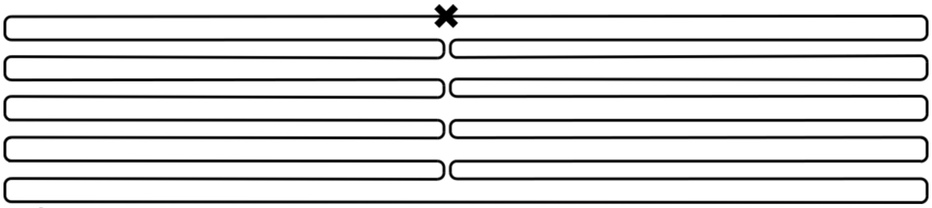

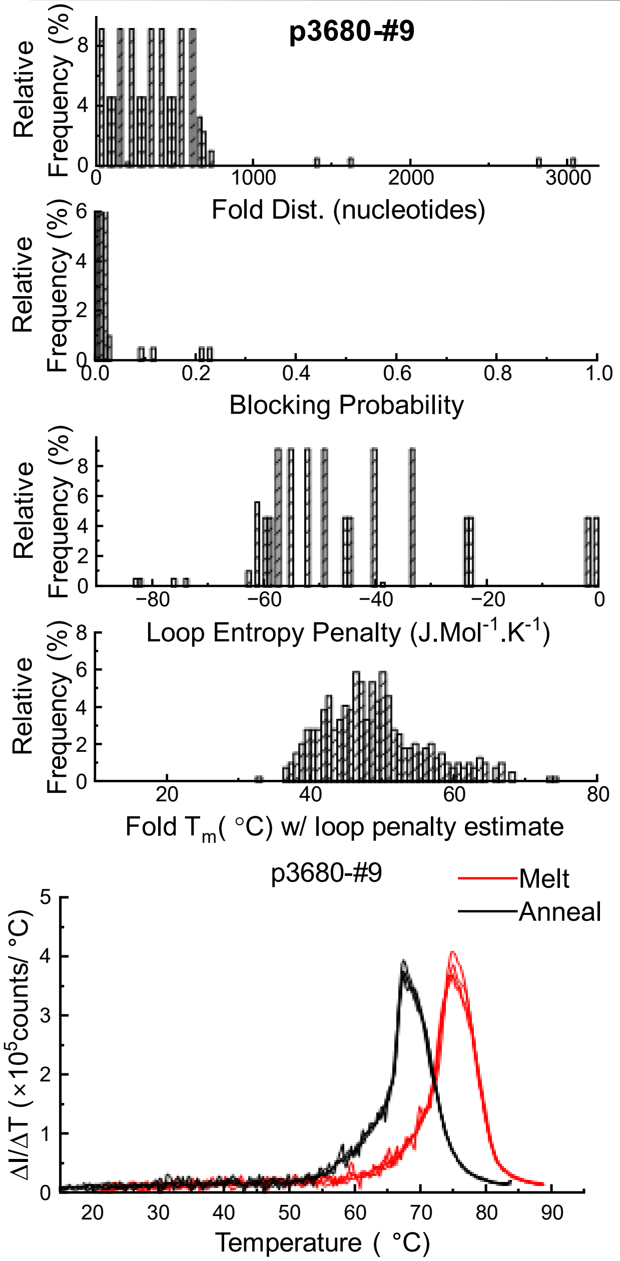

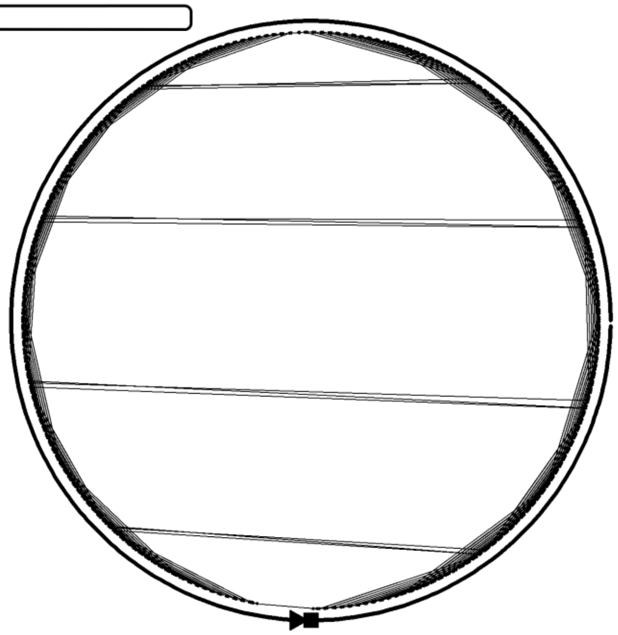

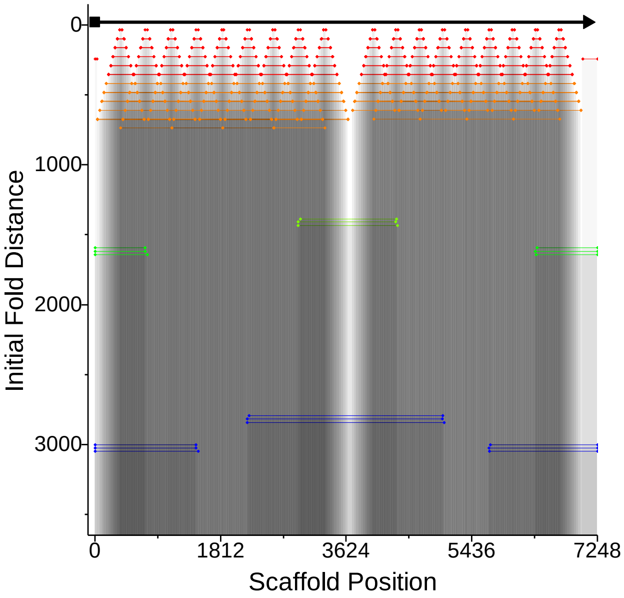

### Design # 10 Horizontal pos. 5630 nucleotides

**ϕ – 9.7 ± 3.2**

**W=32±5 %; D = 27±4 %; R= 36±5 %, C= 5 ±2 %**

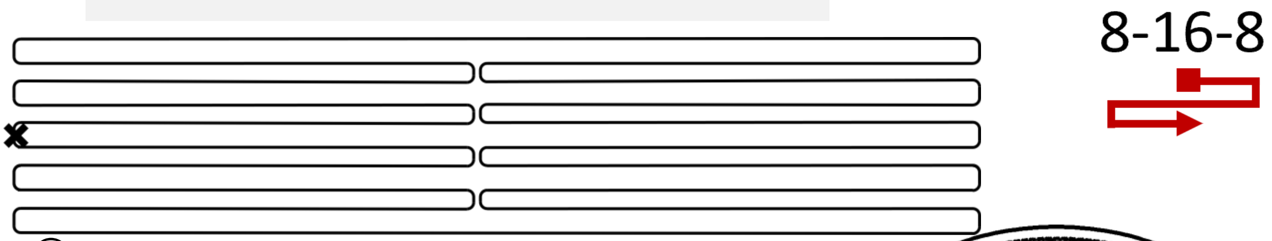

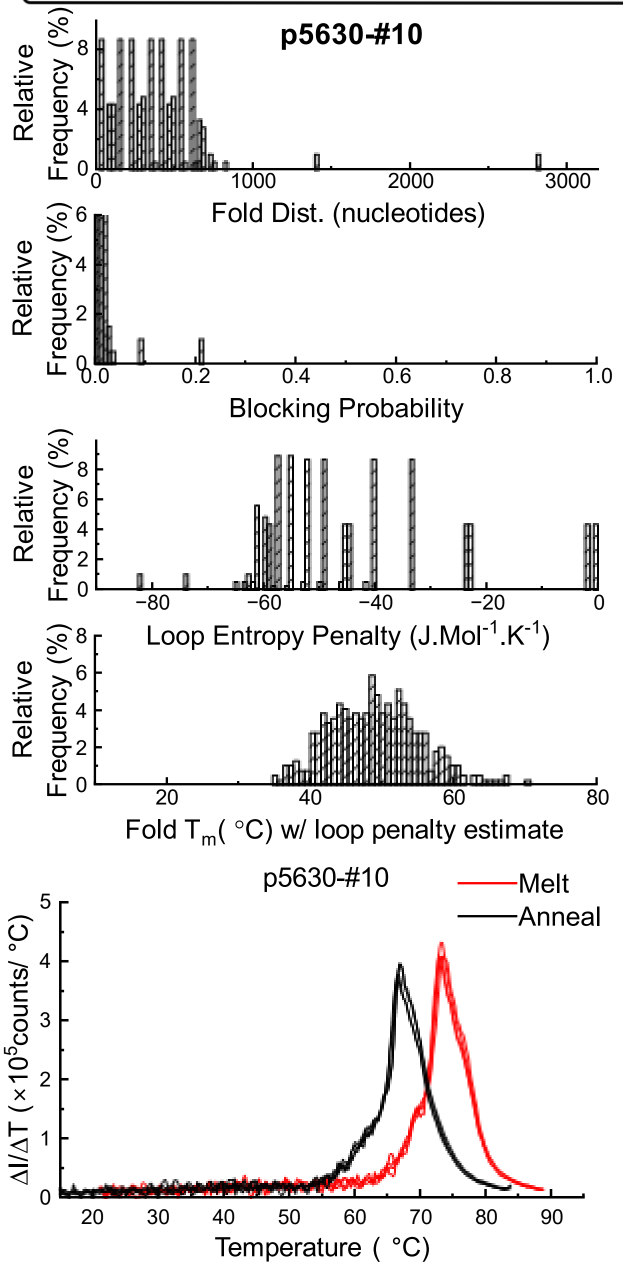

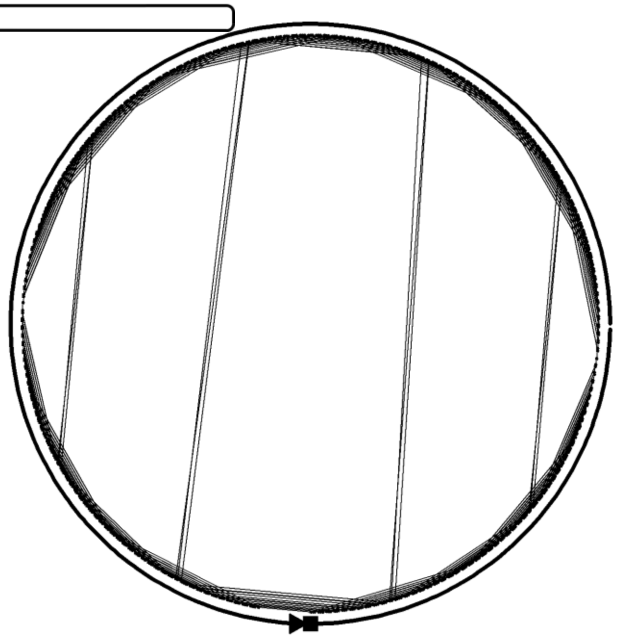

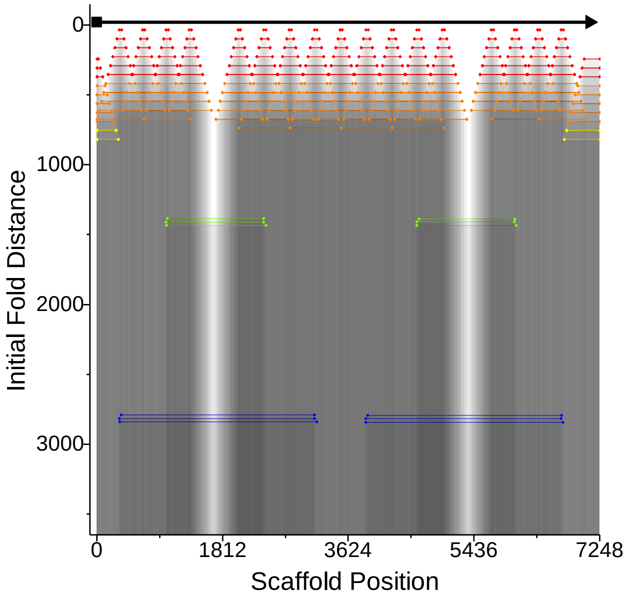

### Design #11 - 2Goatee (pos 736 & 2630)

**ϕ – 32.2 ± 3.4**

**W= 2±1 %; D = 1±2 %; R= 47±4 %, C= 50±3 %**

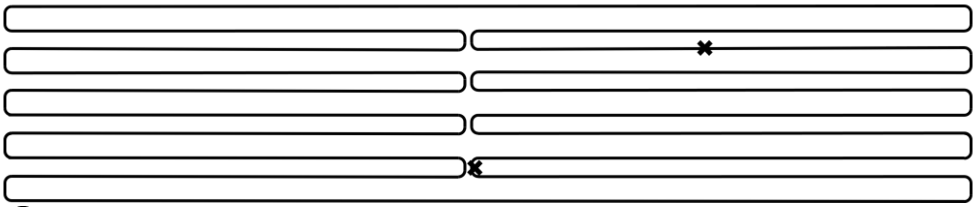

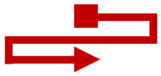

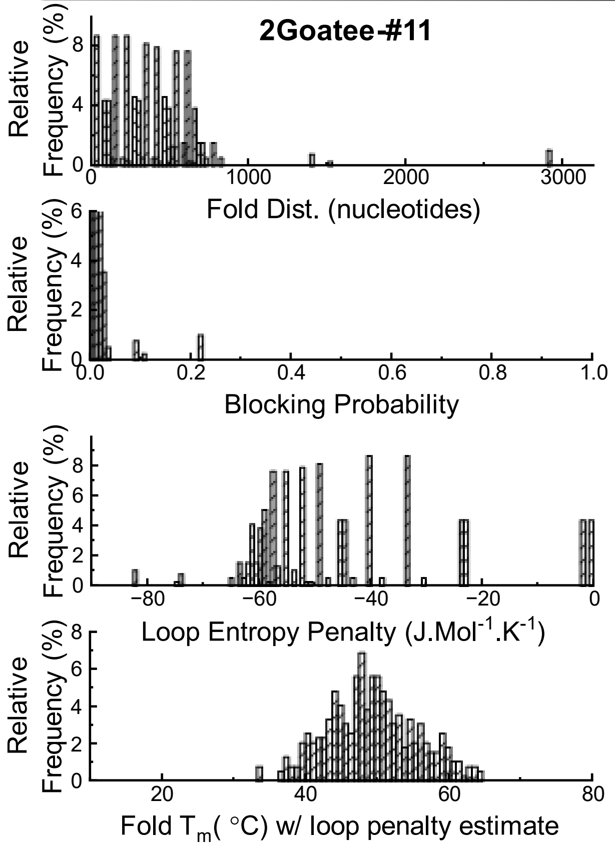

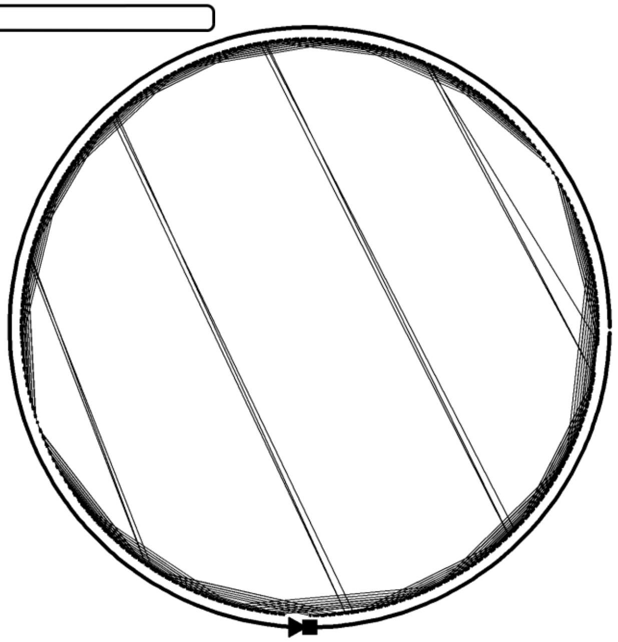

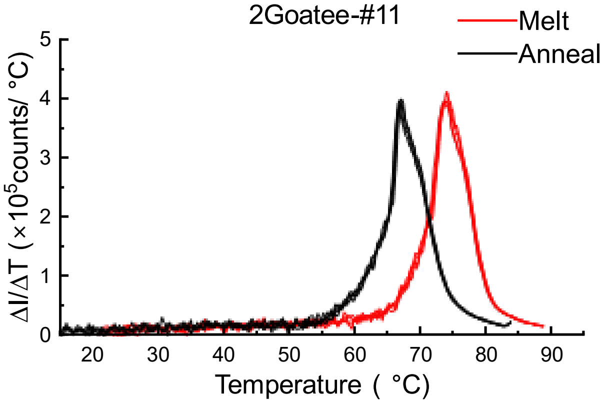

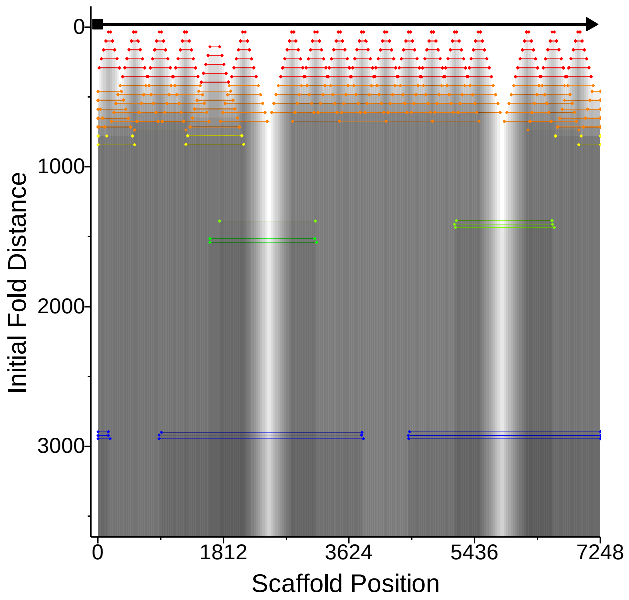

### Design #12 - 3Goatee (neg. 672, pos 736 & 2630)

**ϕ – 30.2 ± 3.3**

**W= 0±0.1 %; D= 1±2 %; R= 56±3 %, C= 43±3 %**

8-16-8

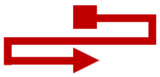

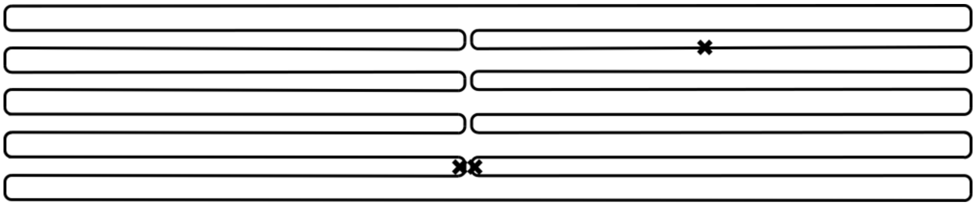

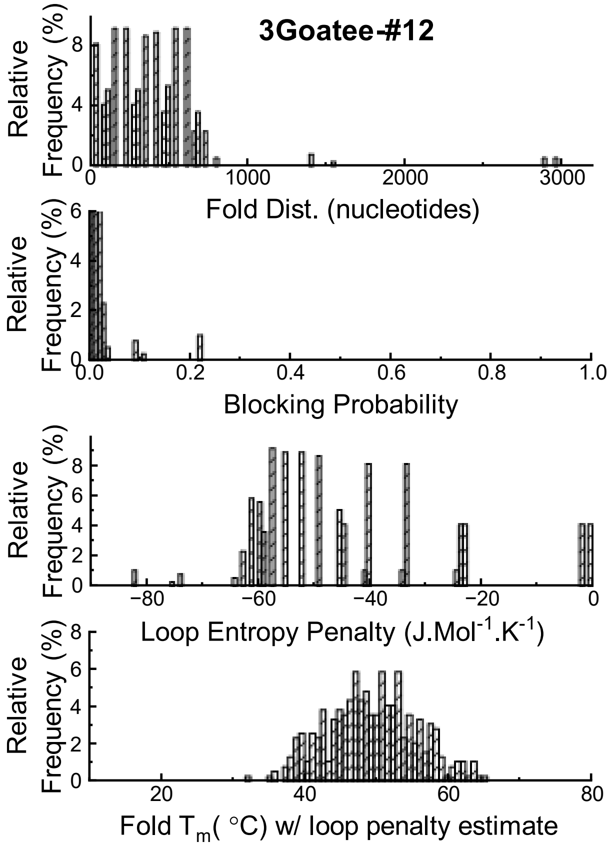

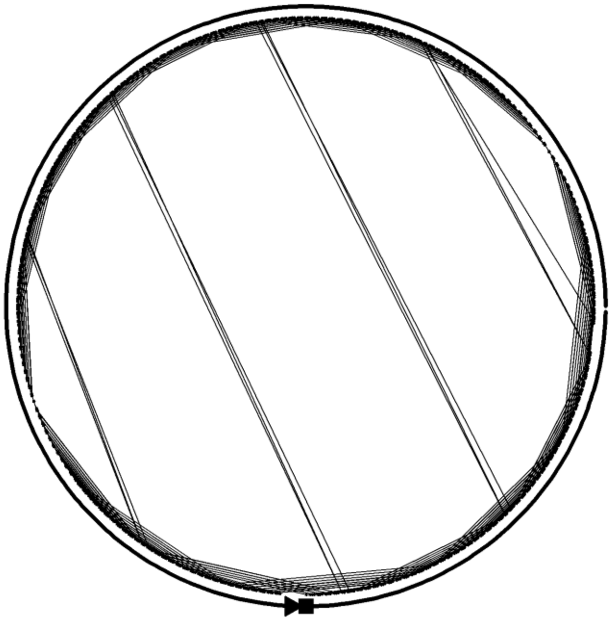

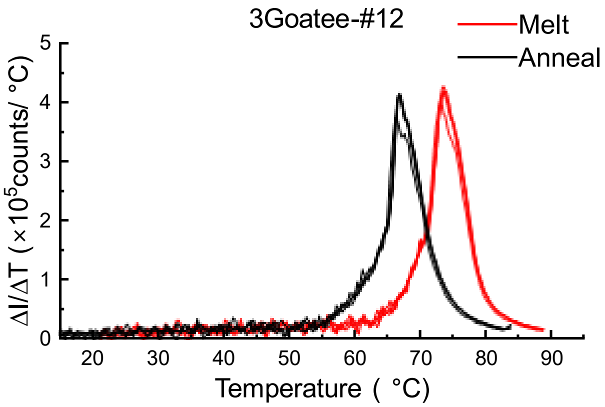

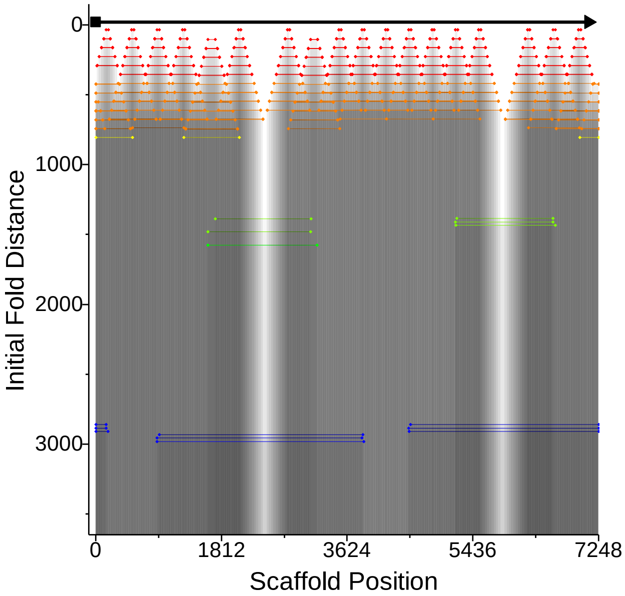

### Design #13 Horizontal – shifted (16 nucleotides)

**ϕ – 10.8 ± 3.3**

**W= 28±3 %; D= 27±3 %; R= 39±3 %, C= 7±1 %**

8-16-8

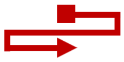

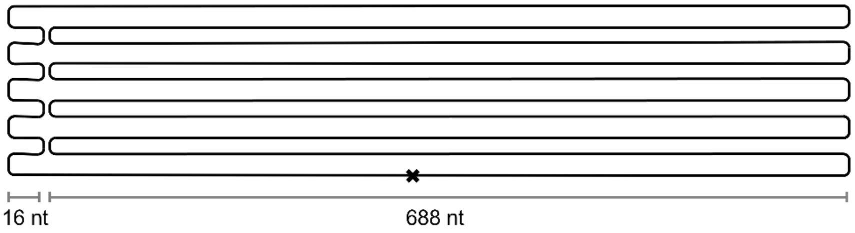

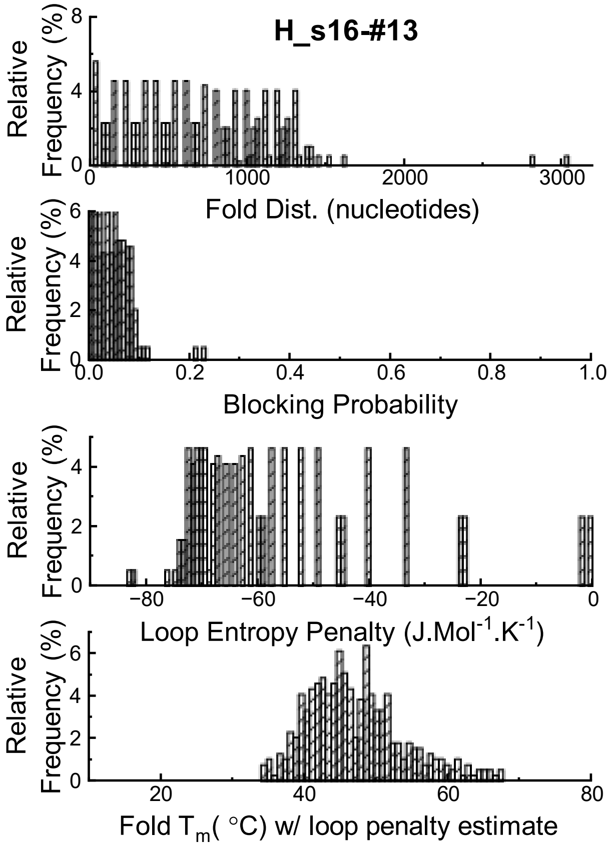

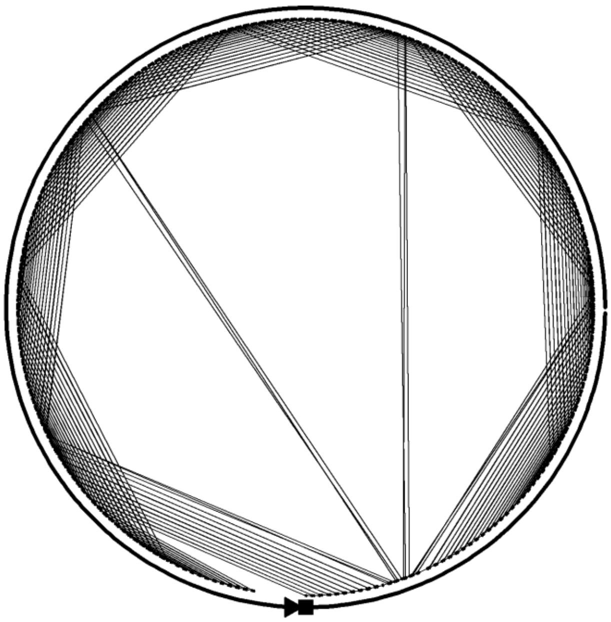

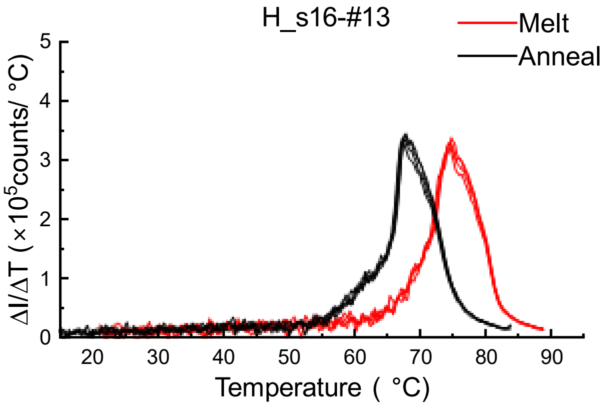

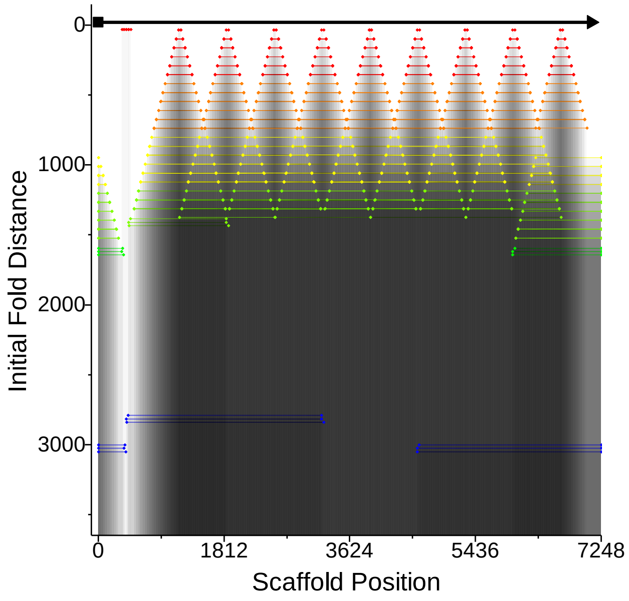

### Design #14 Horizontal – Shifted (144 nucleotides)

**ϕ – 14.8 ± 3.2**

**W= 3±1 %; D = 15±3 %; R= 77±3 %, C= 5±1 %**

8-16-8

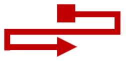

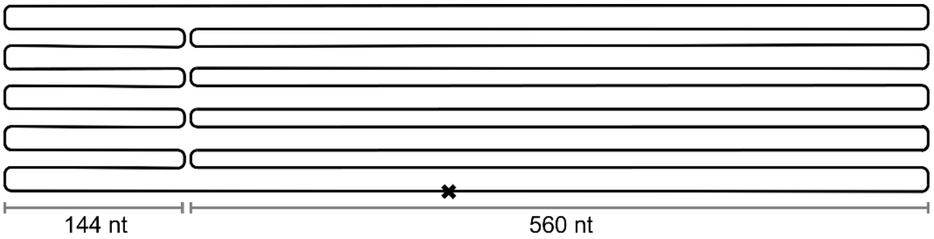

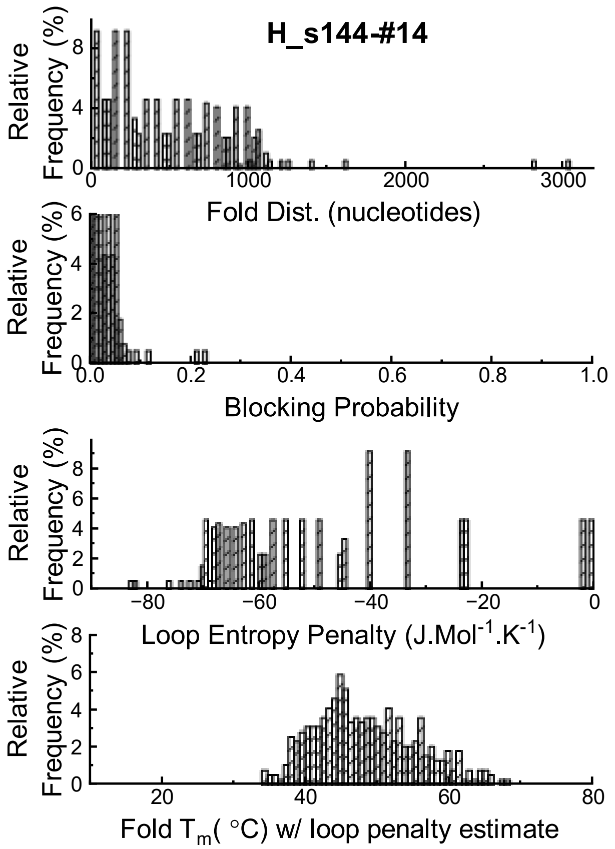

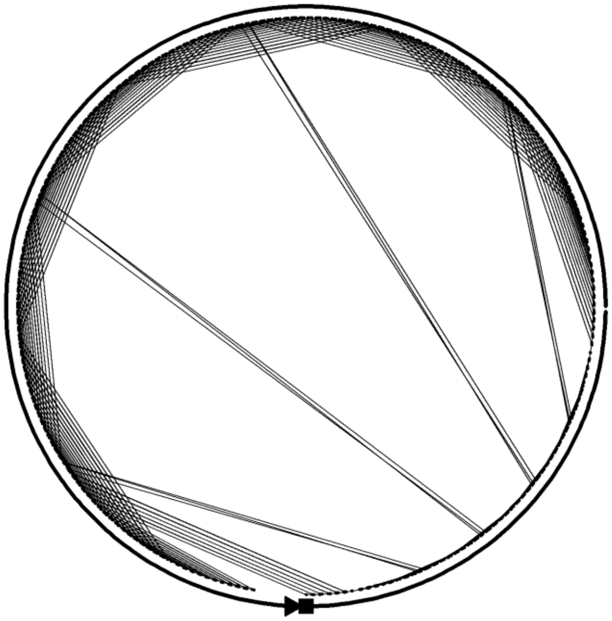

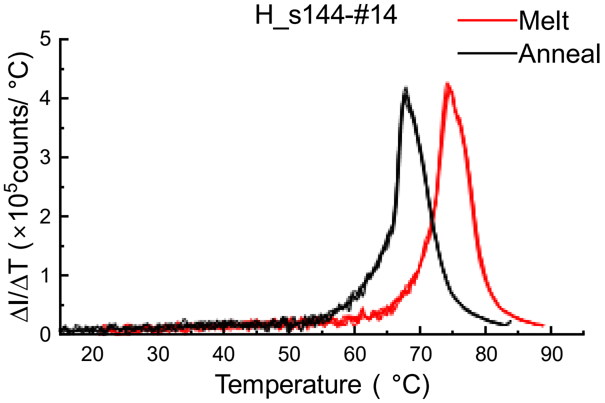

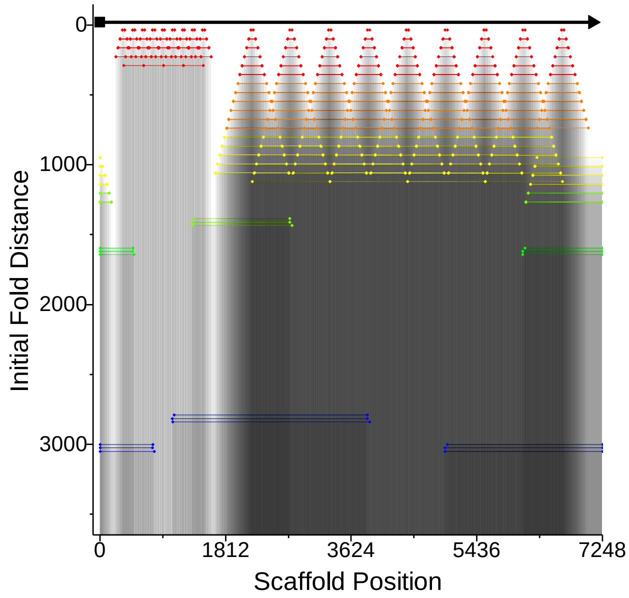

### Design #15 Horizontal – Shifted (260 nucleotides)

**ϕ – 9.9 ± 3.2**

**W= 22±3 %; D = 33±3 %; R= 42±3 %, C= 4±1 %**

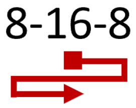

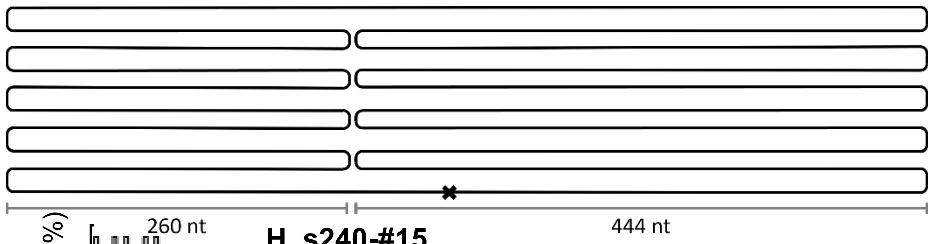

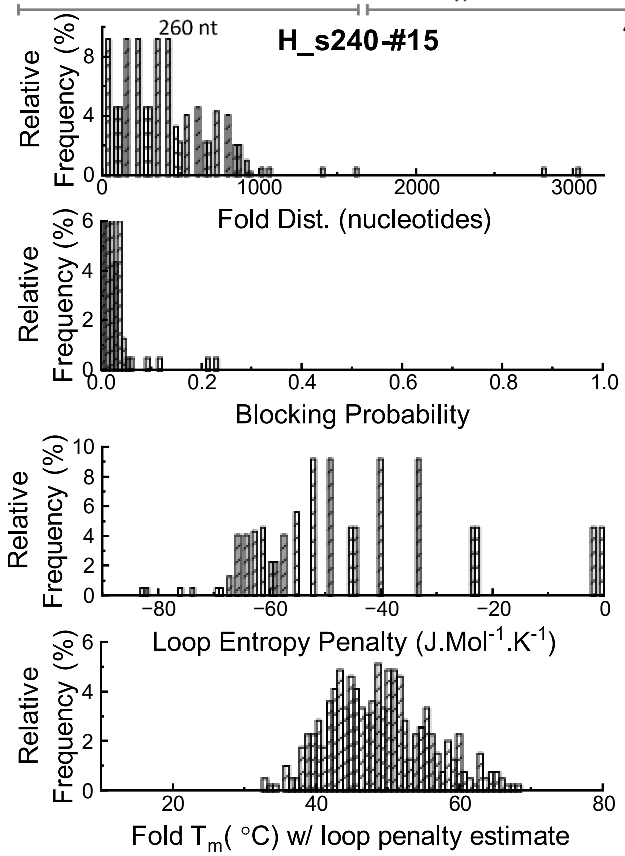

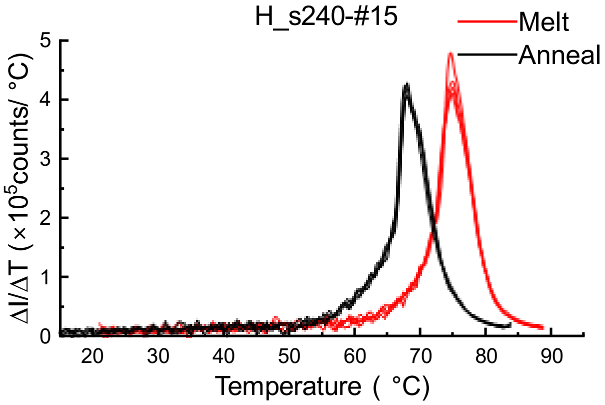

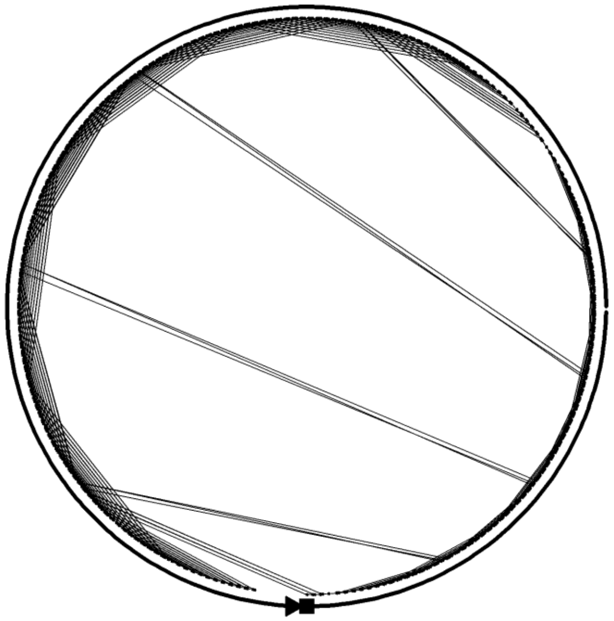

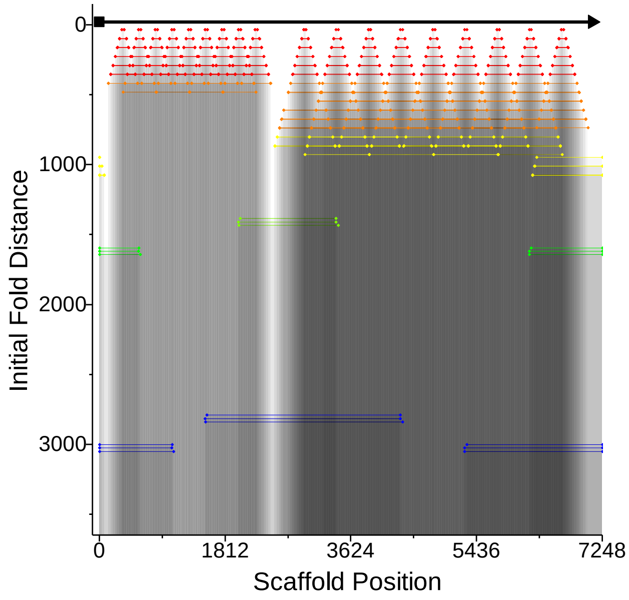

### Design #16 Horizontal – shift (165 nucleotides)

**w/ design flaw -*staples do not bridge seam correctly***

**ϕ – 31.9 ± 3.5**

**W= 5±3 %; D = 33±3 %; R= 42±3 %, C= 4±1 %**

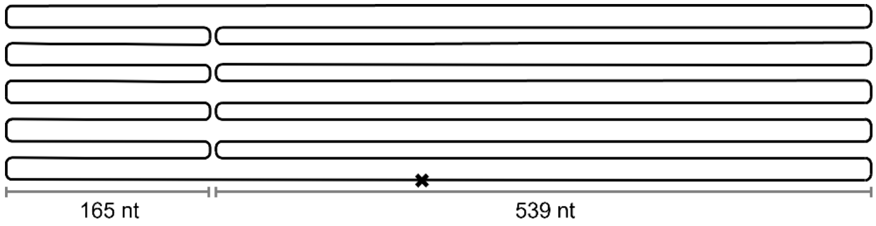

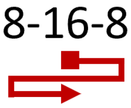

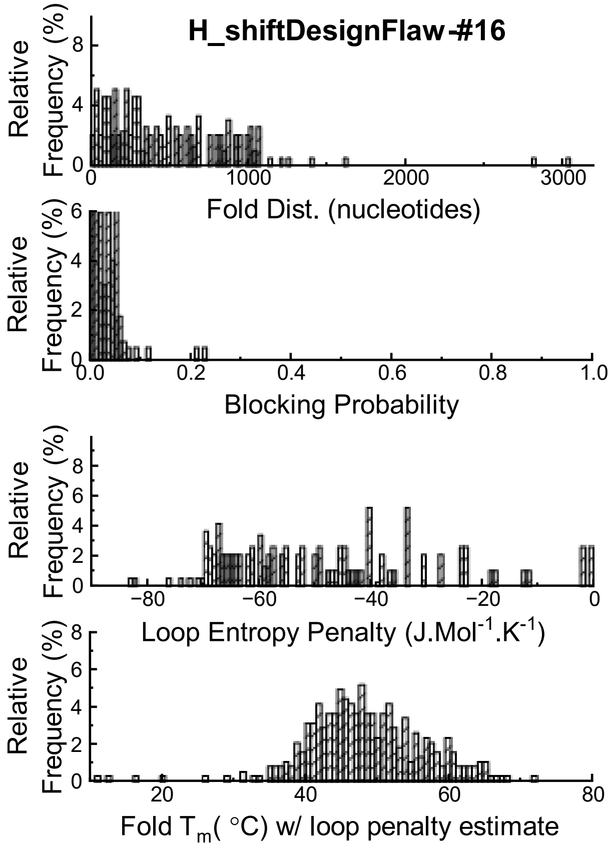

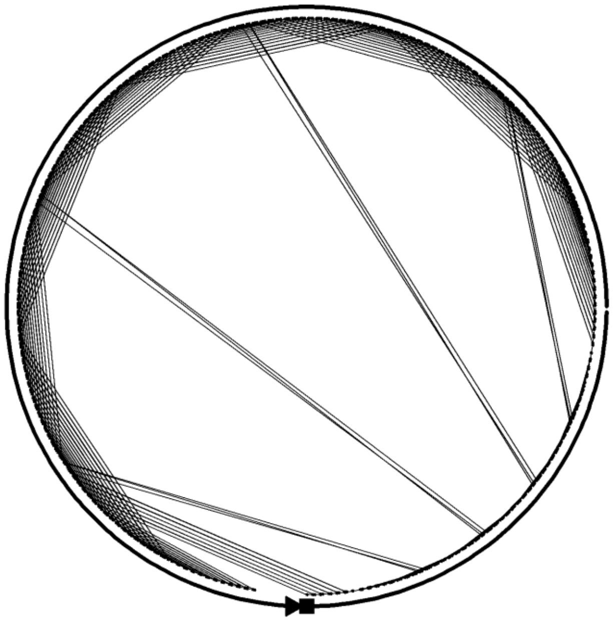

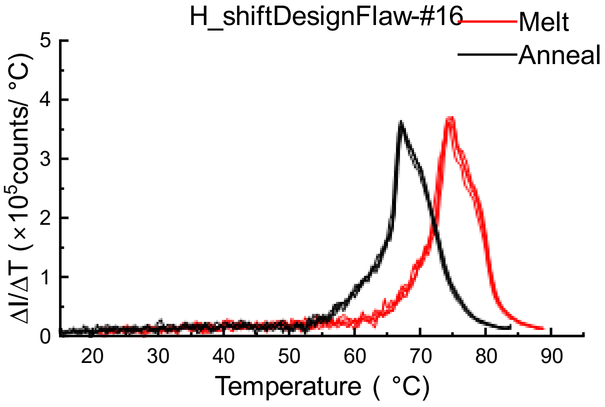

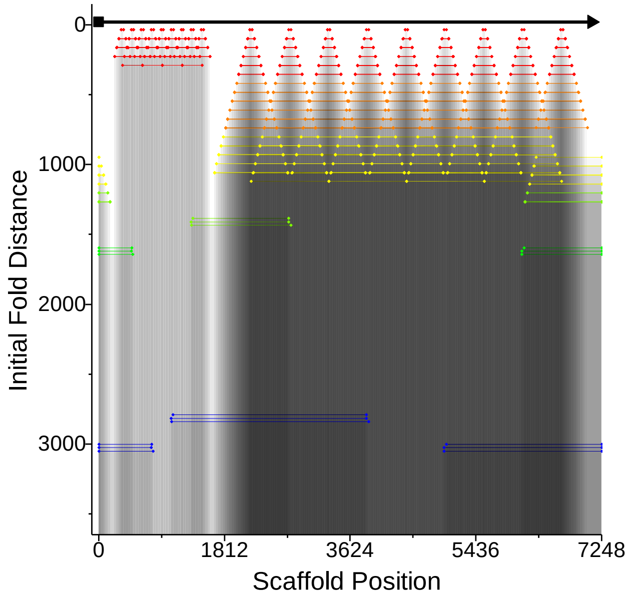

### Design # 17 Horizontal – inter-digitated (16 nucleotides)

**ϕ – 8.4 ± 3.3**

**W= 7±2 %; D= 20±3 %; R= 52±3 %, C= 21±2 %**

8-16-8

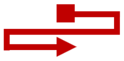

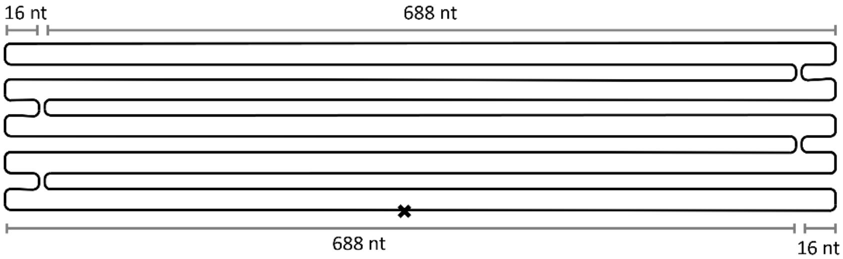

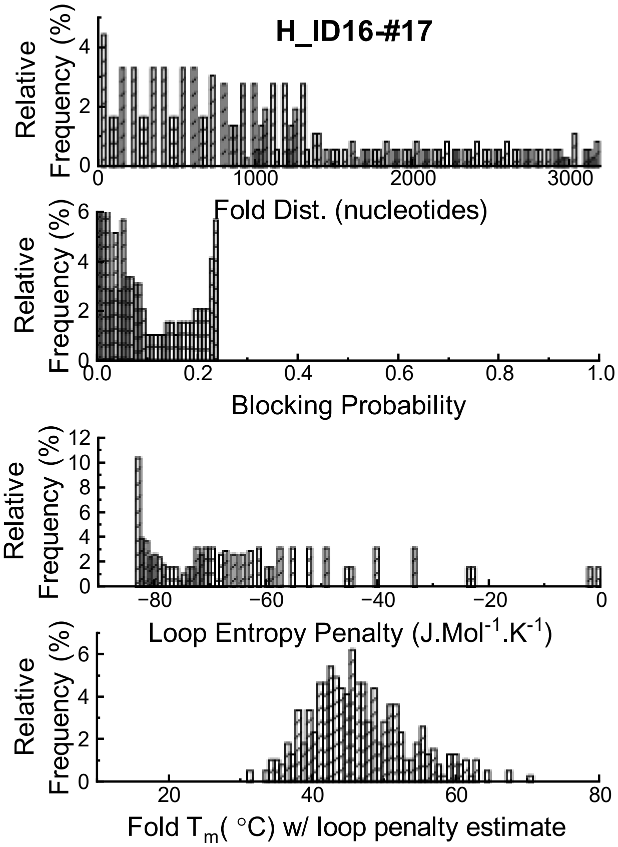

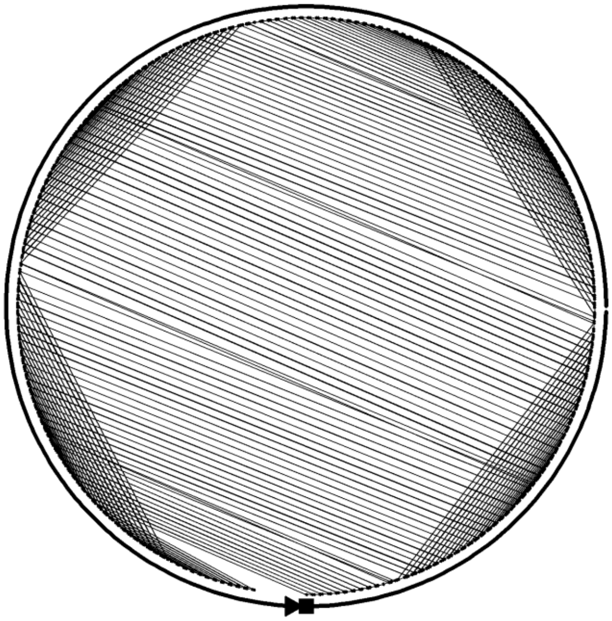

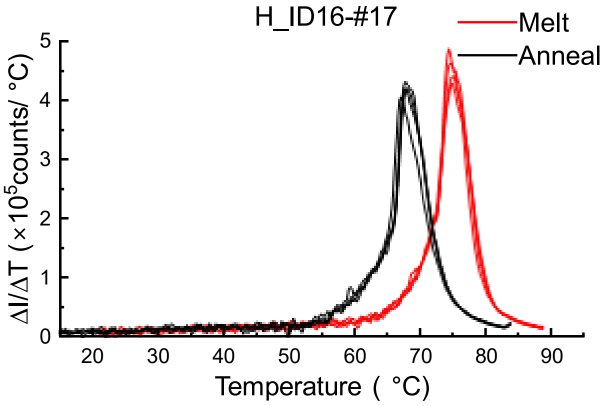

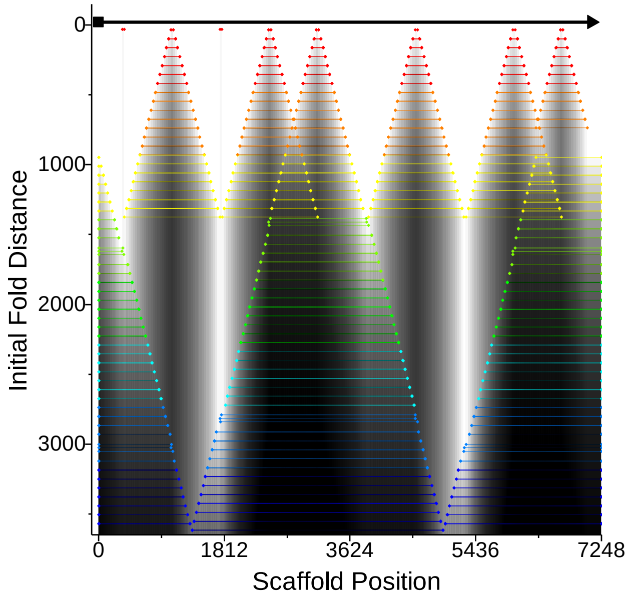

### Design #18 Horizontal – inter-digitated (144 nucleotides)

**ϕ – 16.7 ± 3.3**

**W= 4±2 %; D= 24±3 %; R= 58±3 %, C= 14±2 %**

8-16-8

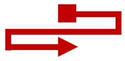

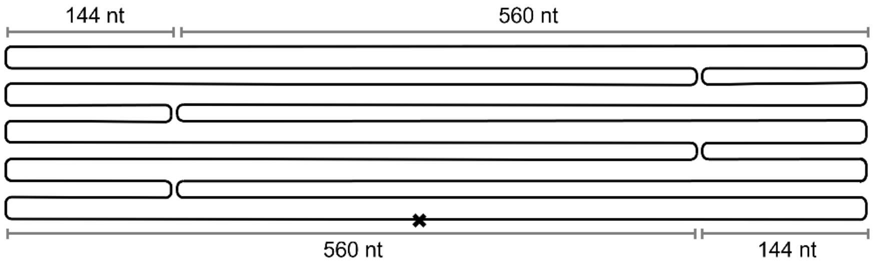

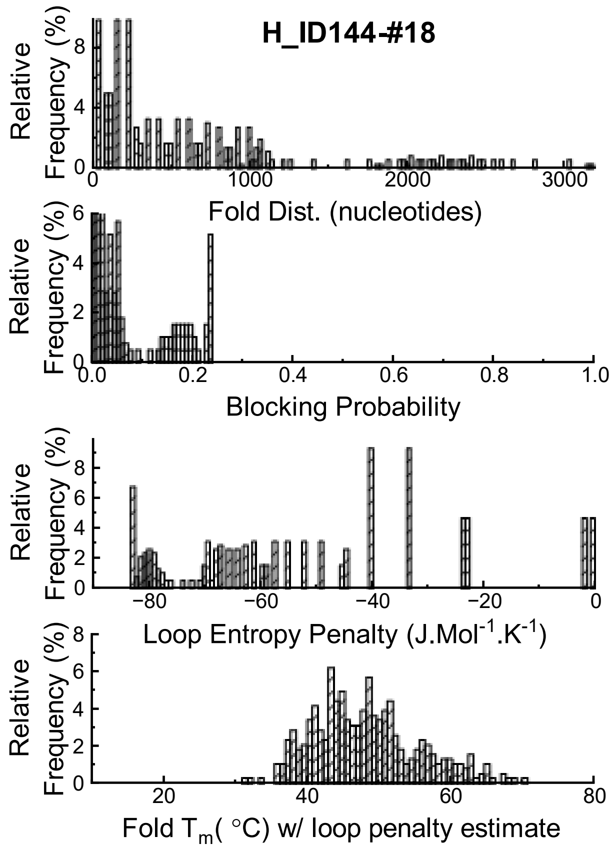

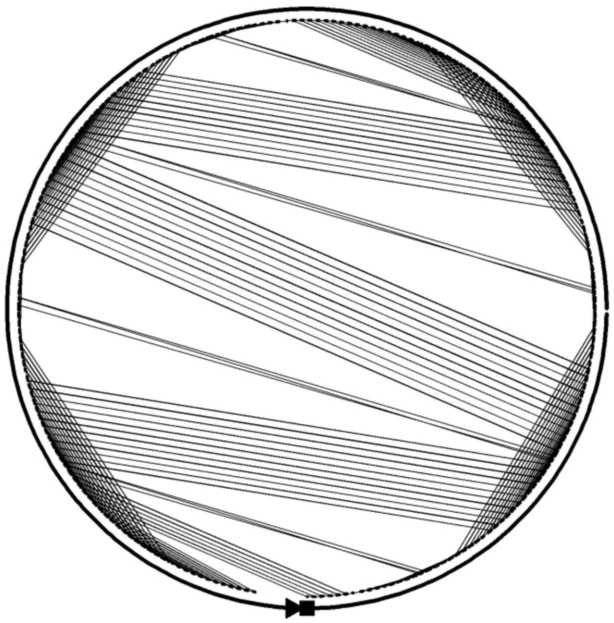

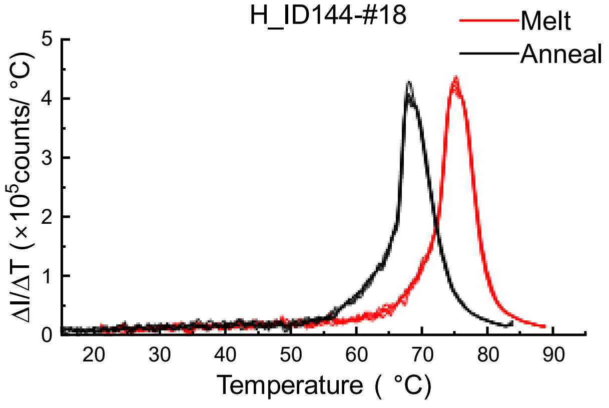

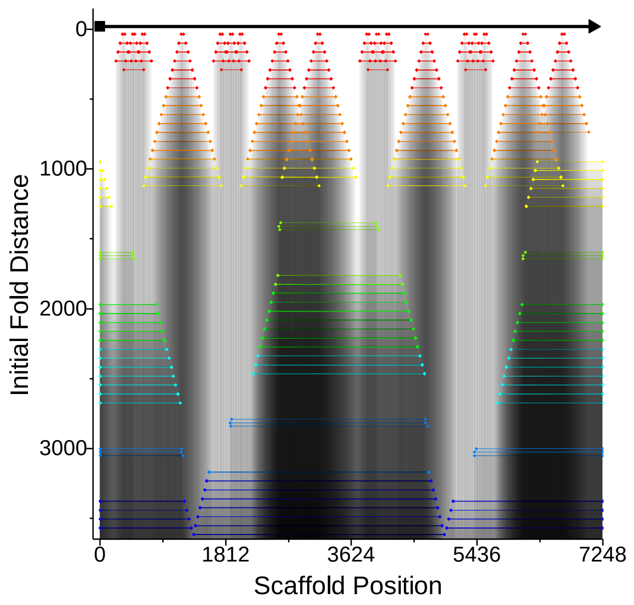

### Design #19 Horizontal – inter-digitated (260 nucleotides)

**ϕ – 12.7 ± 3.3**

**W= 12±3 %; D= 25±3 %; R= 58±3 %, C= 5±1 %**

8-16-8

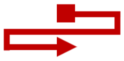

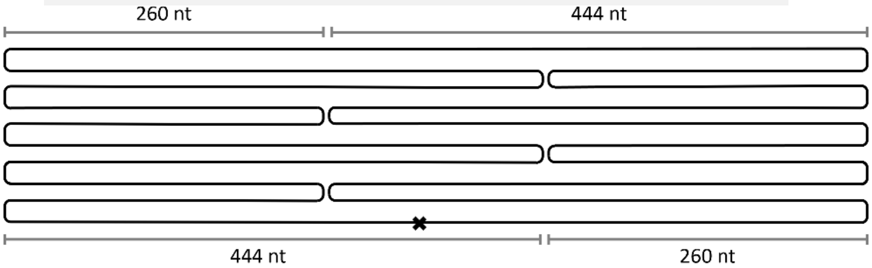

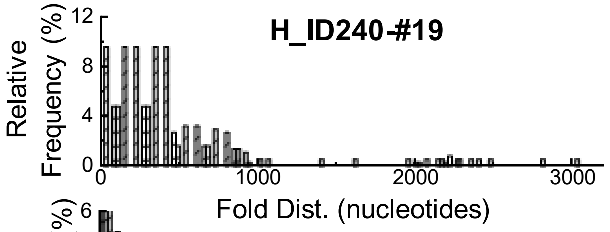

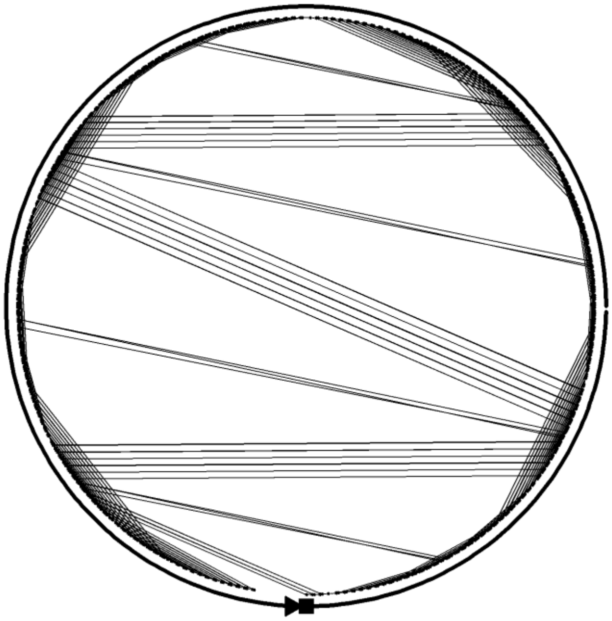

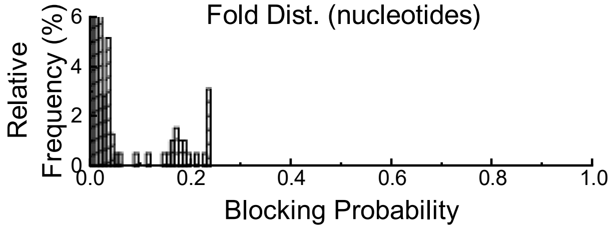

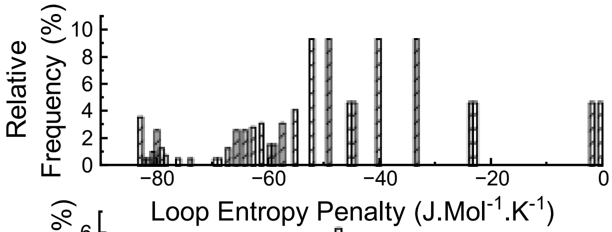

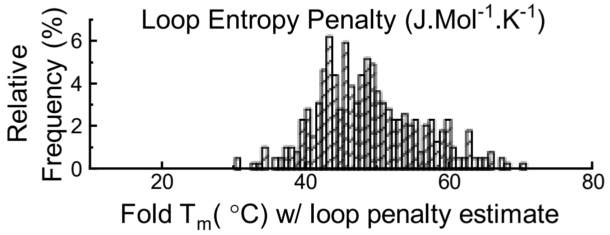

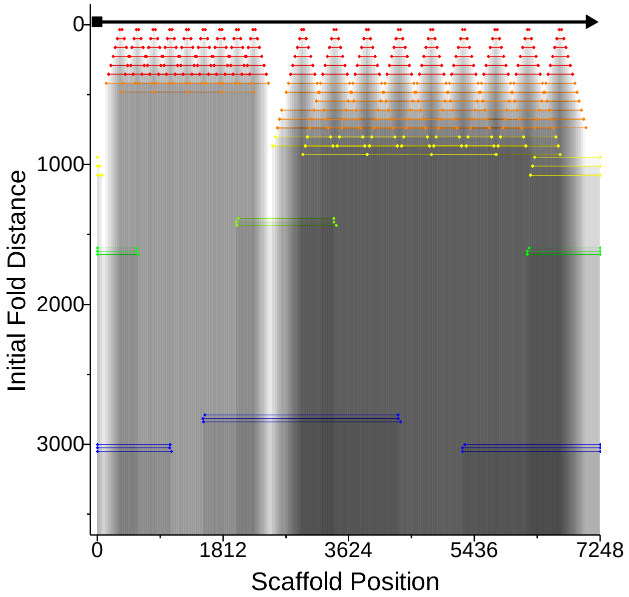

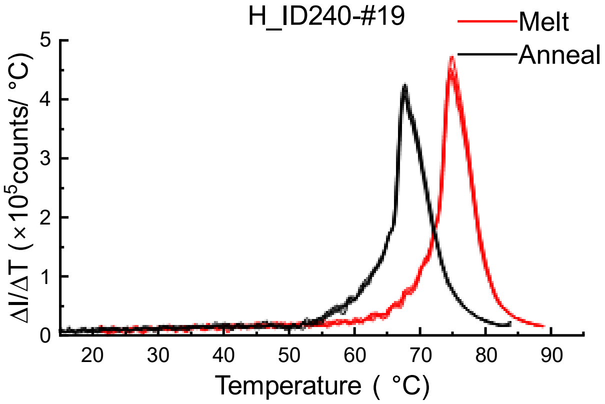

### Design #20 Horizontal – inter-digitated (272 nucleotides)

**ϕ – 8.9 ± 3.3**

**W= 27±3 %; D = 37±3 %; R= 32±3 %, C= 4±1 %**

8-16-8

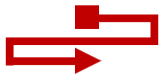

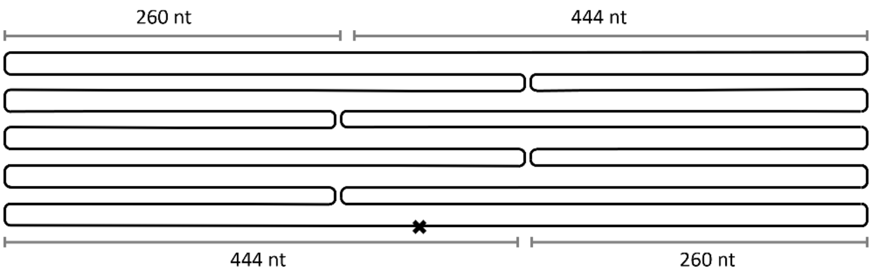

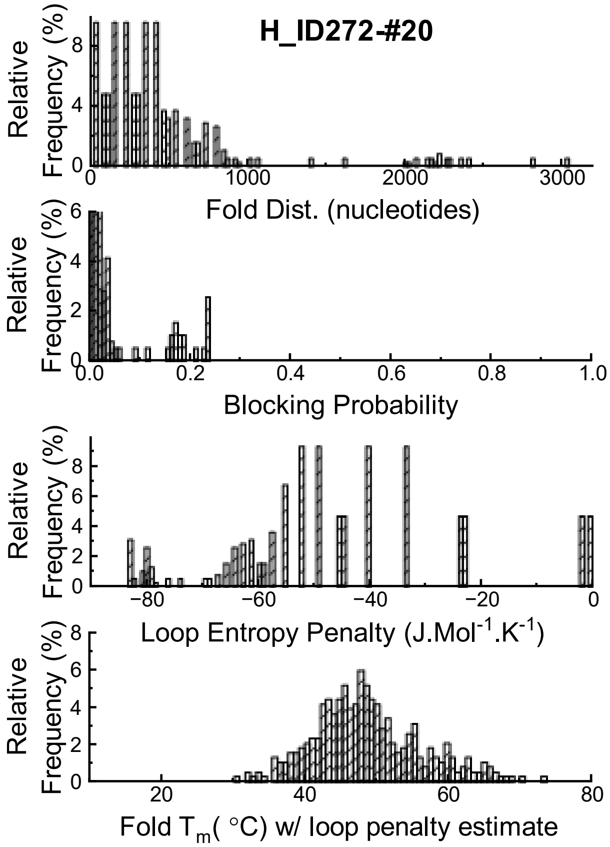

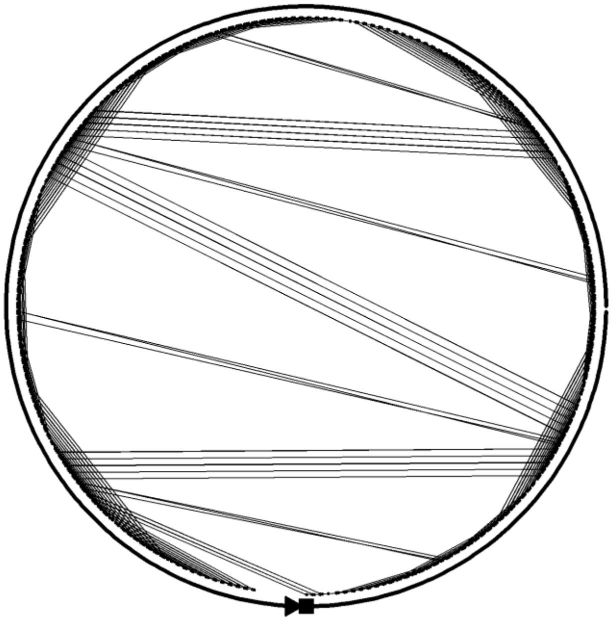

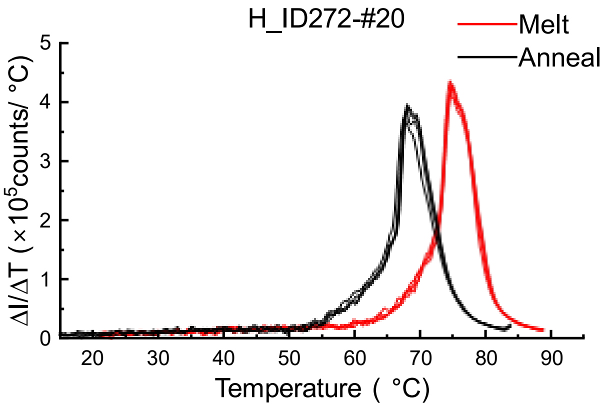

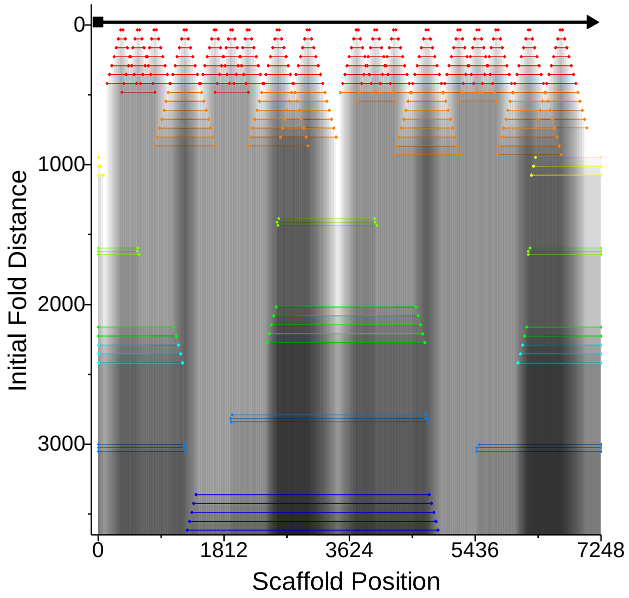

### Design #21 - 3 Seam horizontal

**ϕ – 11.0 ± 3.3**

**W= 15±2 %; D = 34±3 %; R= 44±3 %, C= 4±1 %**

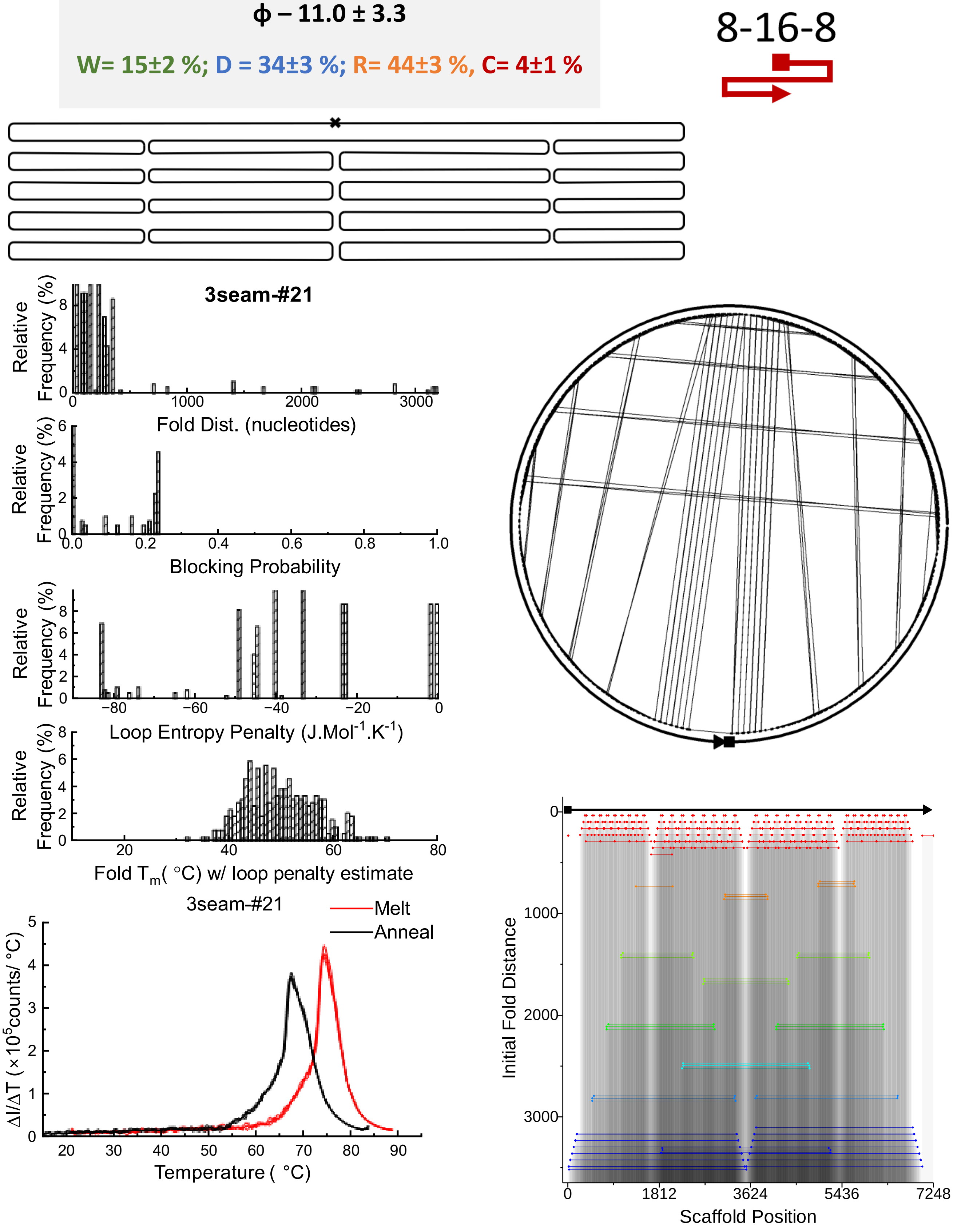

### Design #22 Notched Rectangle

**ϕ – 21.7 ± 3.3**

**W= 2±1 %; D = 22±2 %;**

**R= 50±3 %, C= 26±2 %**

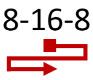

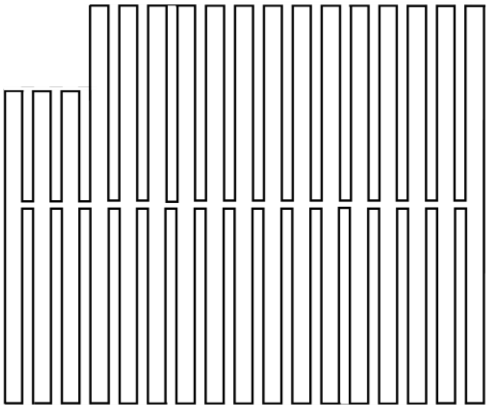

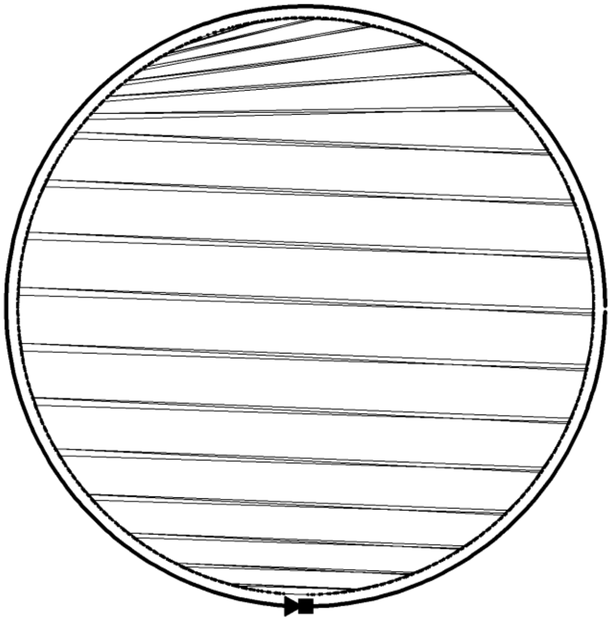

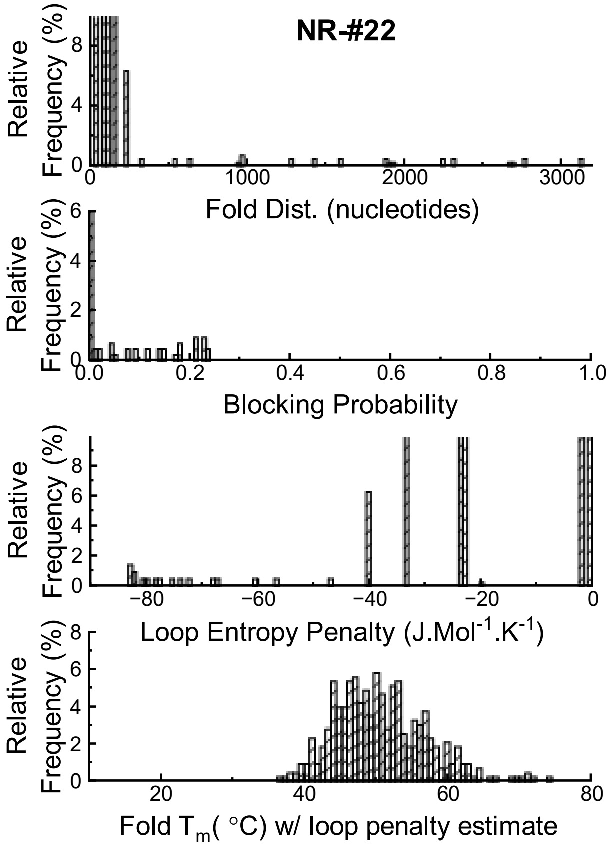

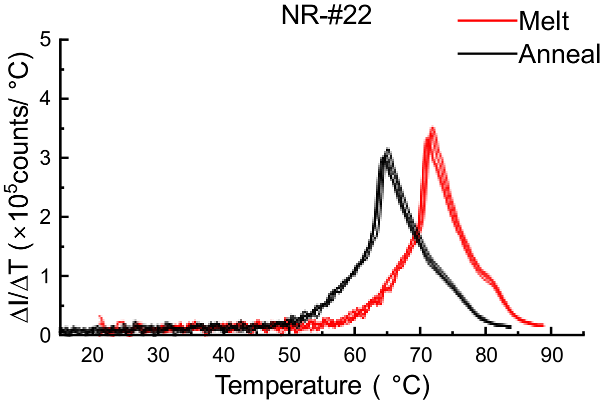

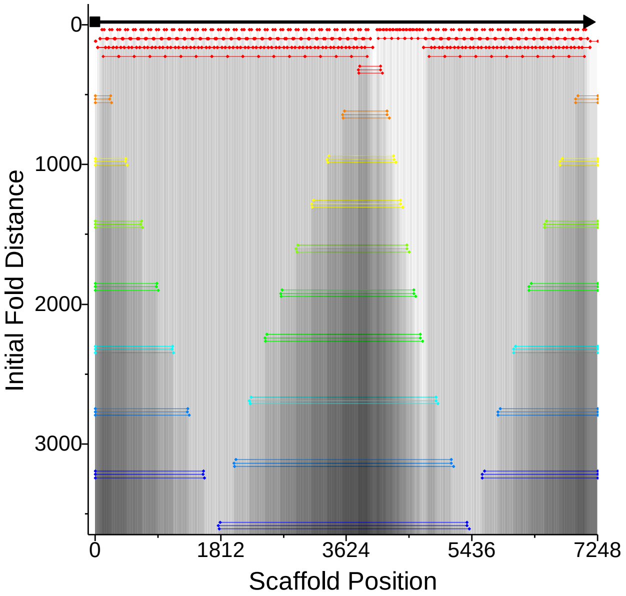

### Design #23 – Horizontal Weave, 16-16

**ϕ – 17.7 ± 3.3**

**W= 7±2 %; D = 21±3 %; R= 55±3 %, C= 17±2 %**

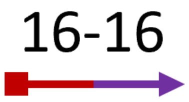

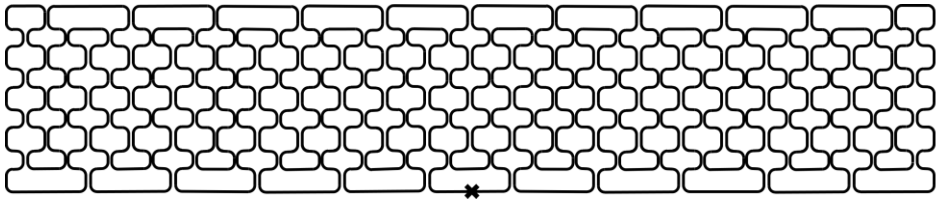

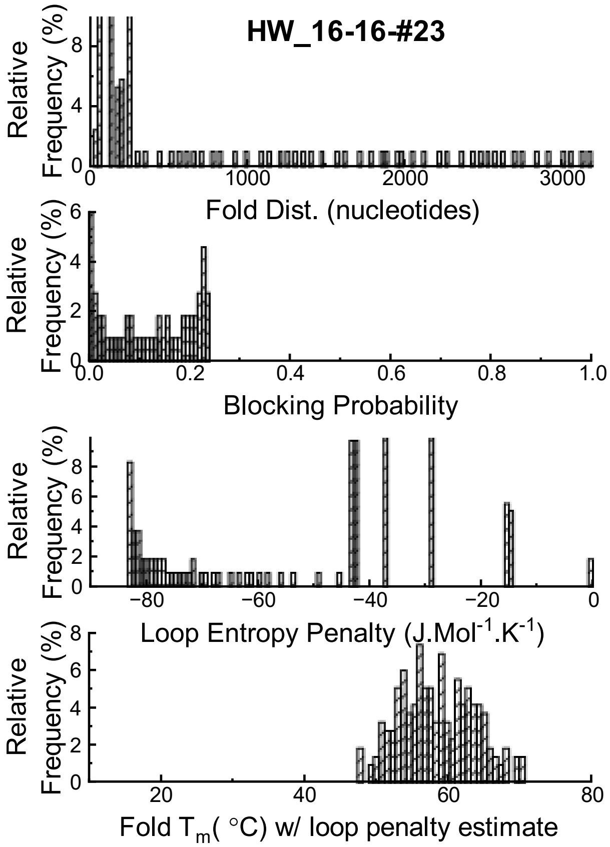

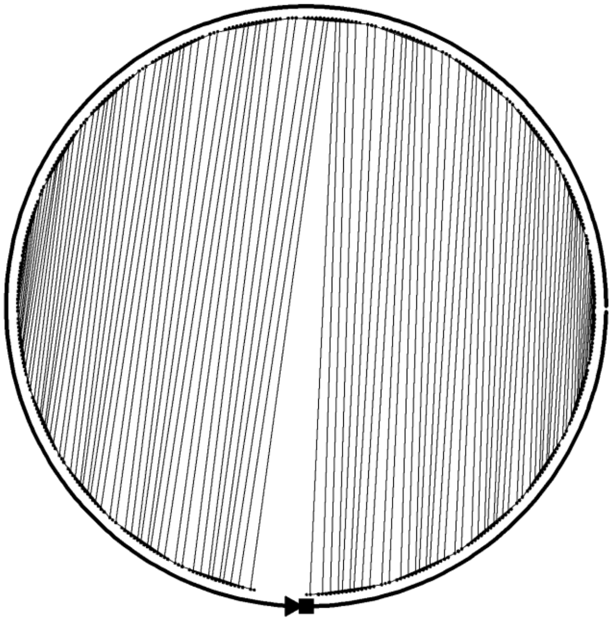

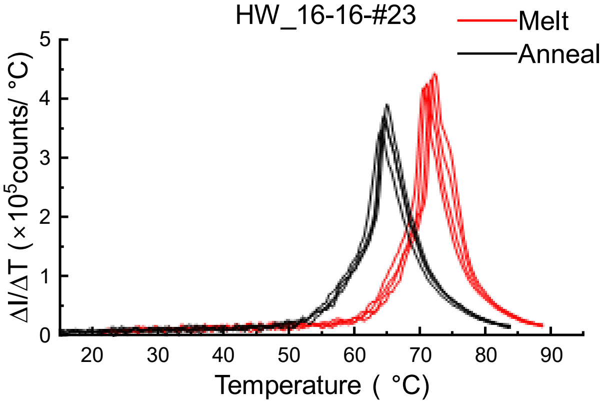

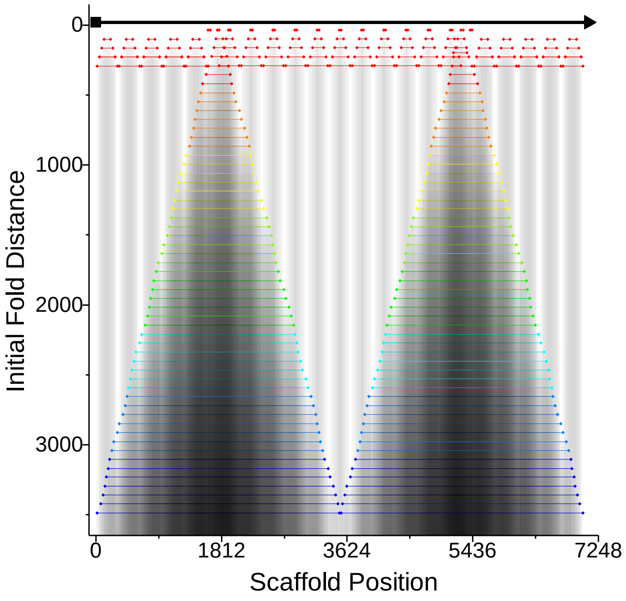

### Design #24 Vertical Origami 8-16-8

**ϕ – 25.4 ± 3.5**

**W= 0±1 %; D = 6±3 %; R= 62±4 %, C= 32±4 %**

8-16-8

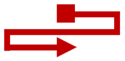

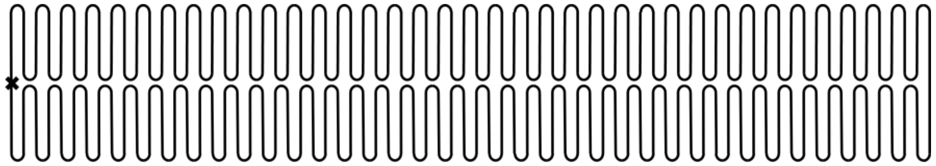

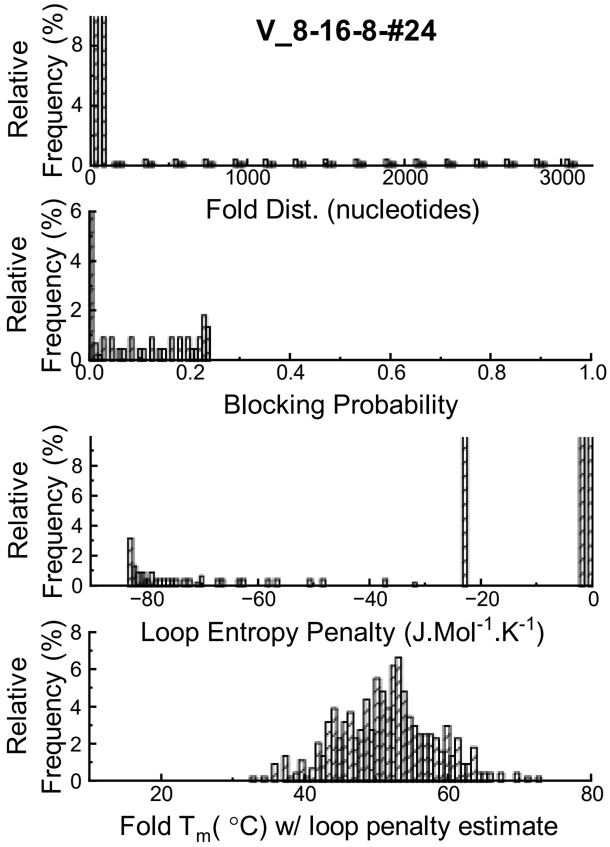

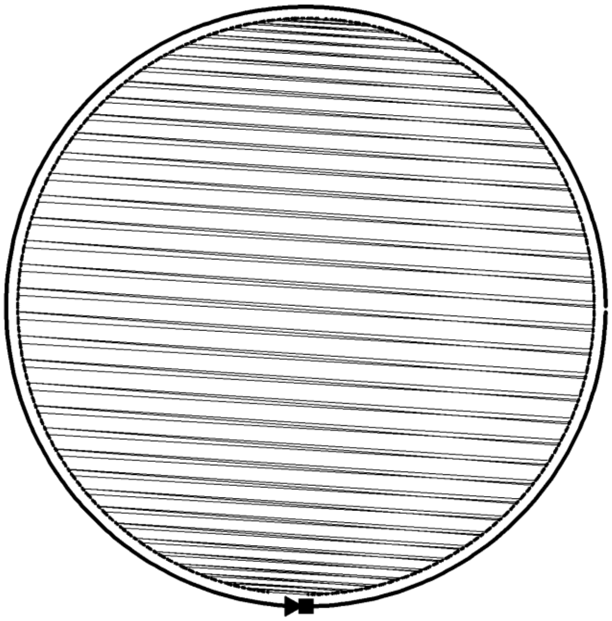

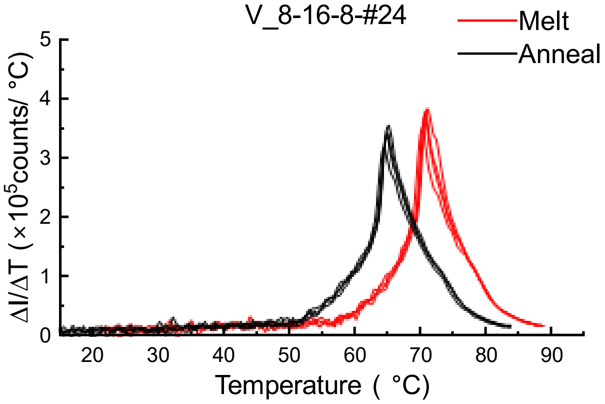

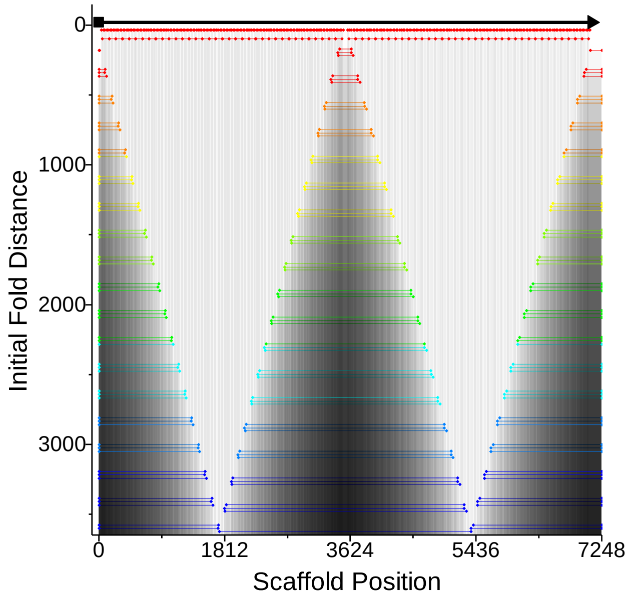

### Design #25 Vertical Origami – 16-16

**ϕ – 15.0 ± 3.3**

**W= 7±1 %; D = 20±3 %; R= 65±3 %, C= 8±1 %**

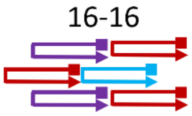

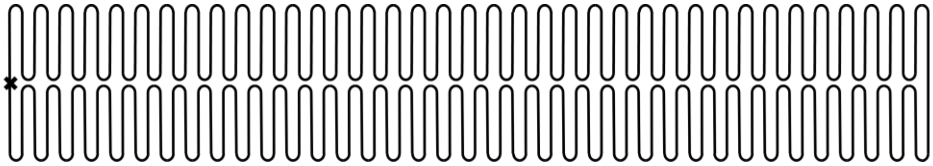

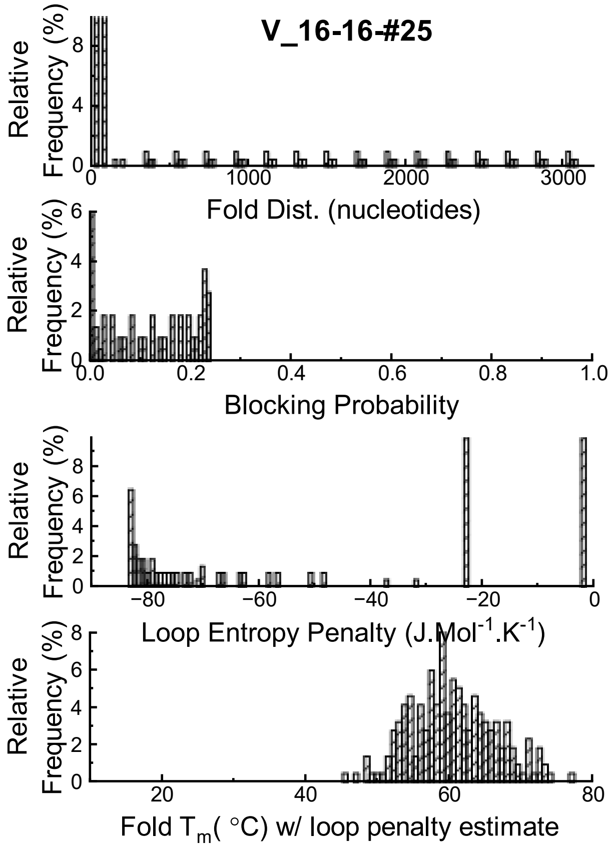

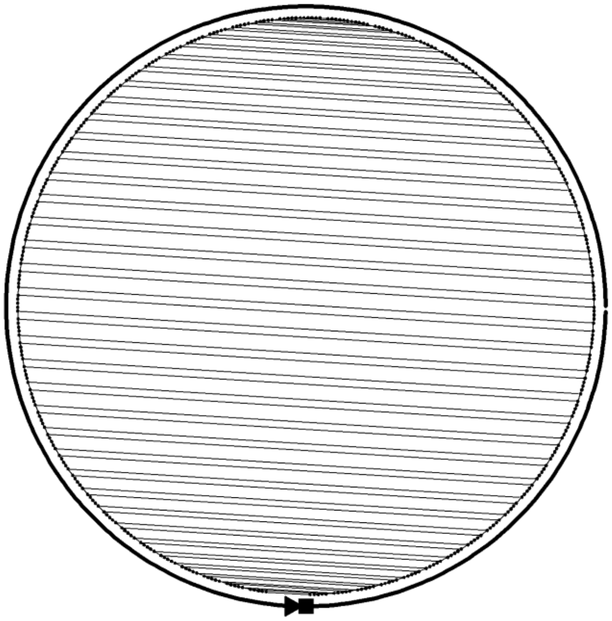

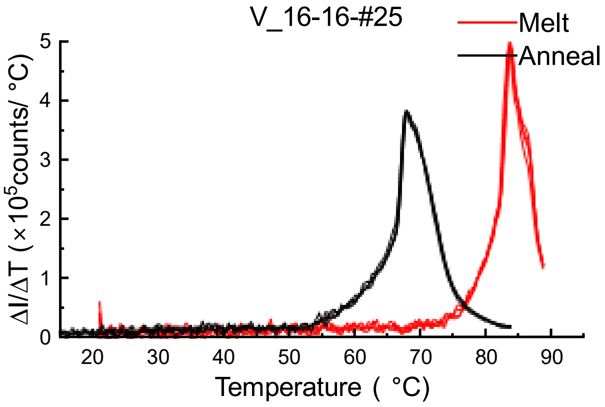

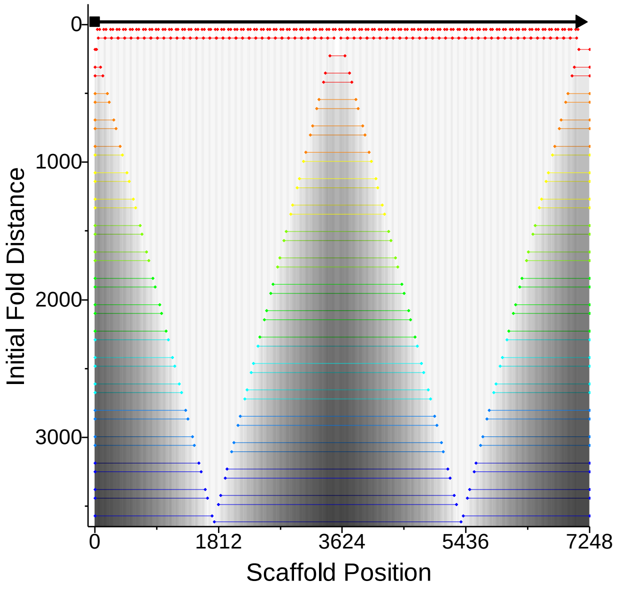

### Design #26 Vertical Origami – 8-16-16-16-8

**ϕ – 14.0 ± 3.3**

**W= 3±1 %; D = 26±3 %; R= 65±4 %, C= 6±4 %**

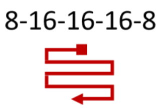

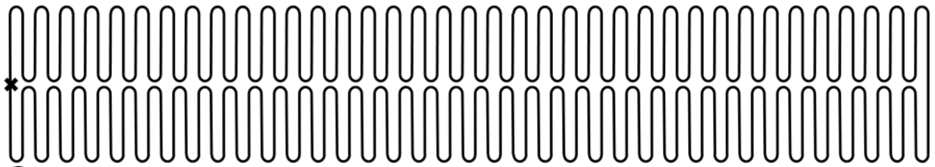

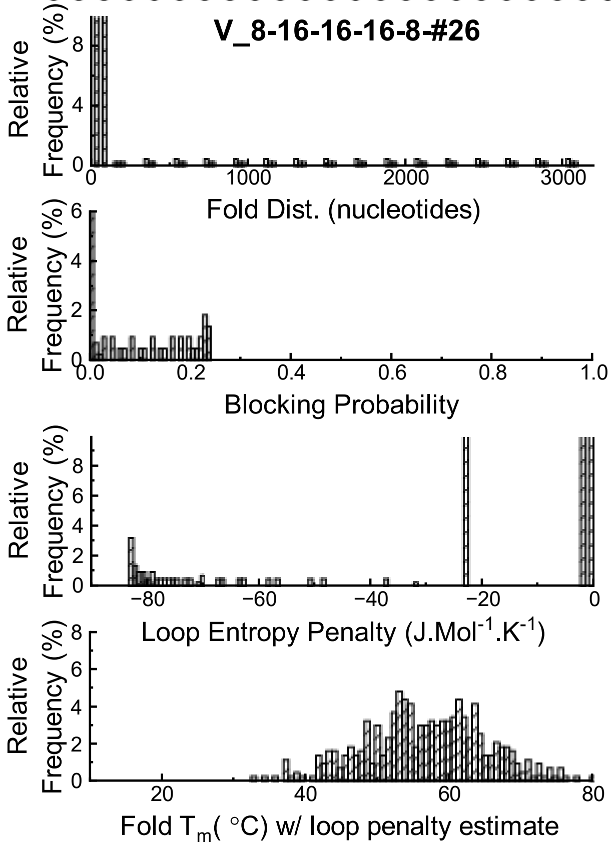

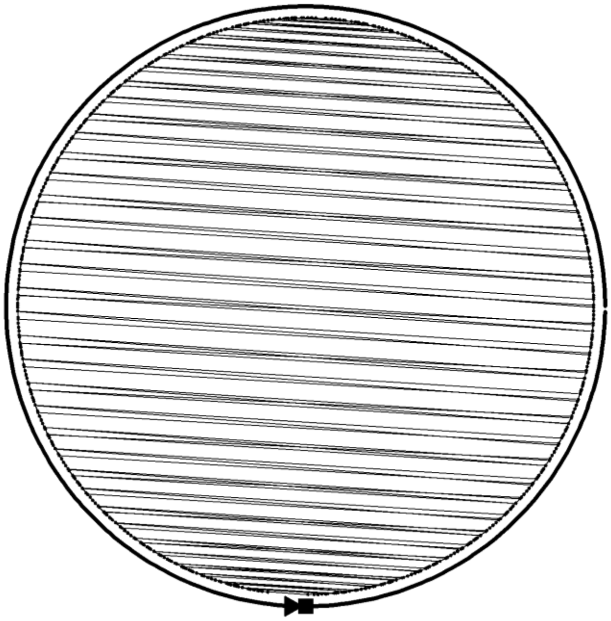

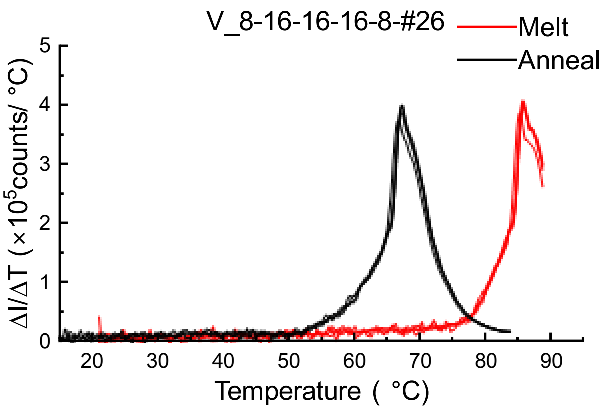

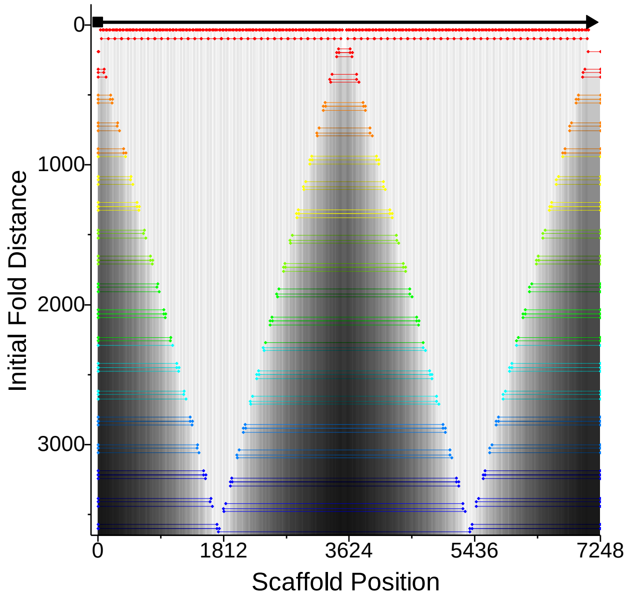

### Design #27,hex  Horizontal, homogenous domains

**ϕ – 26.1 ± 3.3**

**W= 8±1 %; D = 33±3 %; R= 56±4 %, C= 3±3 %**

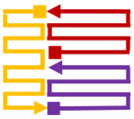

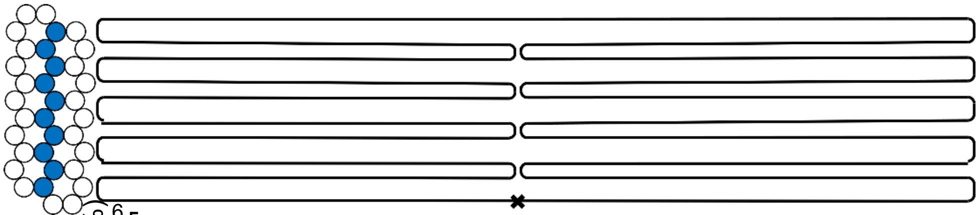

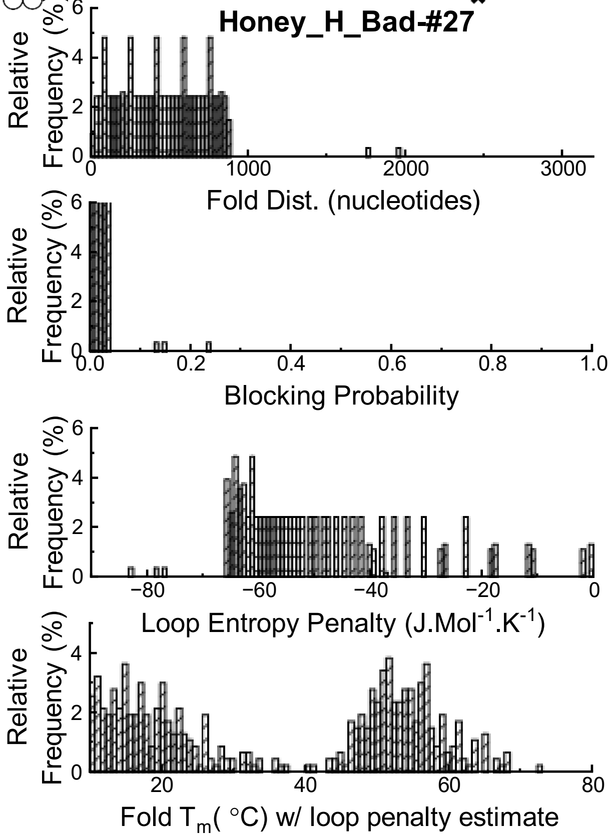

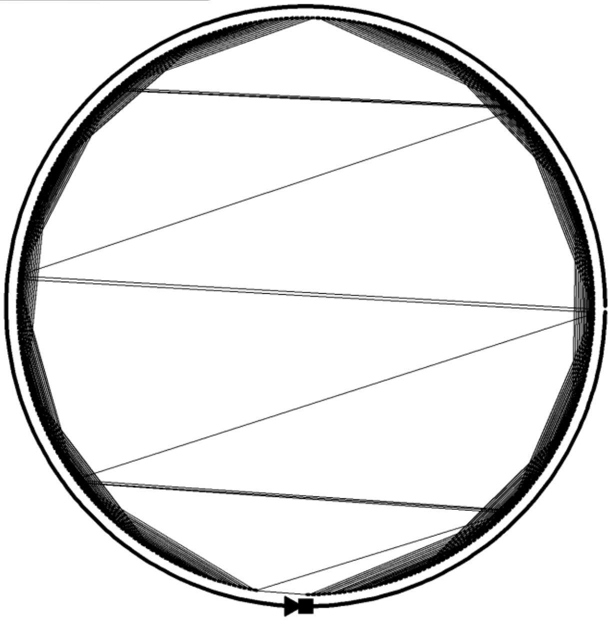

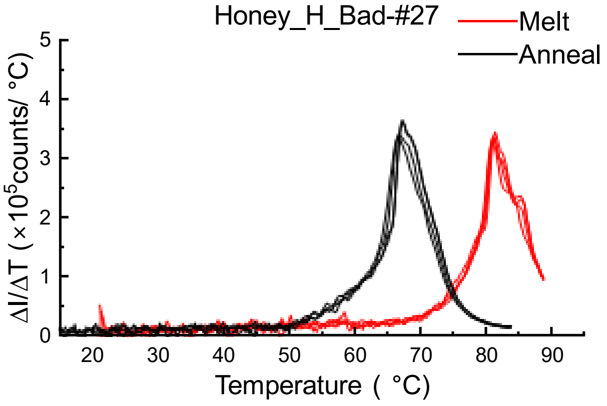

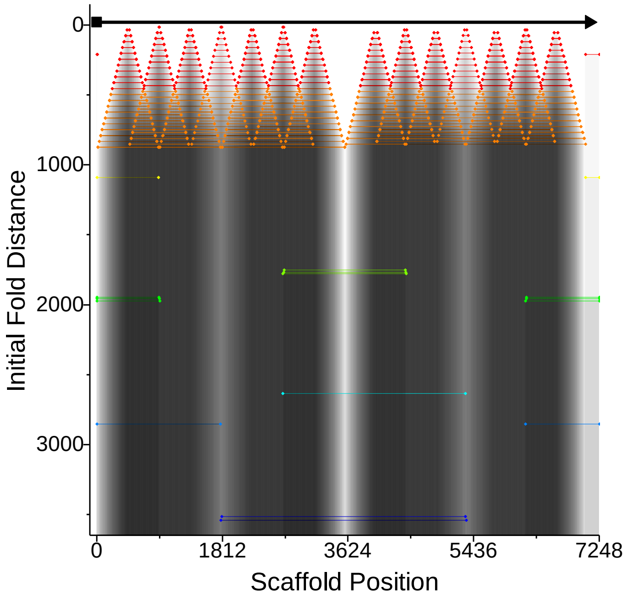

### Design #28 Hex, Vertical, homogeneous staples

**ϕ – 26.1 ± 3.5**

**W= 0±1 %; D = 9±3 %; R= 57±4 %, C= 34±3 %**

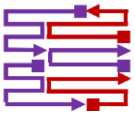

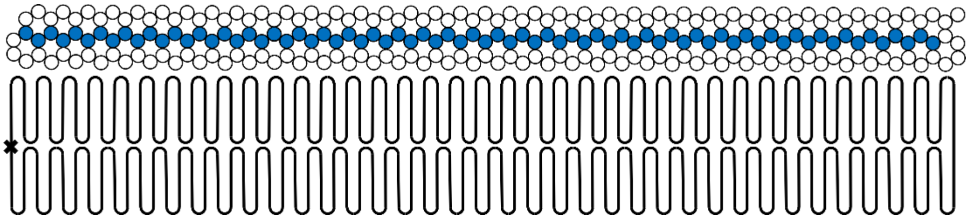

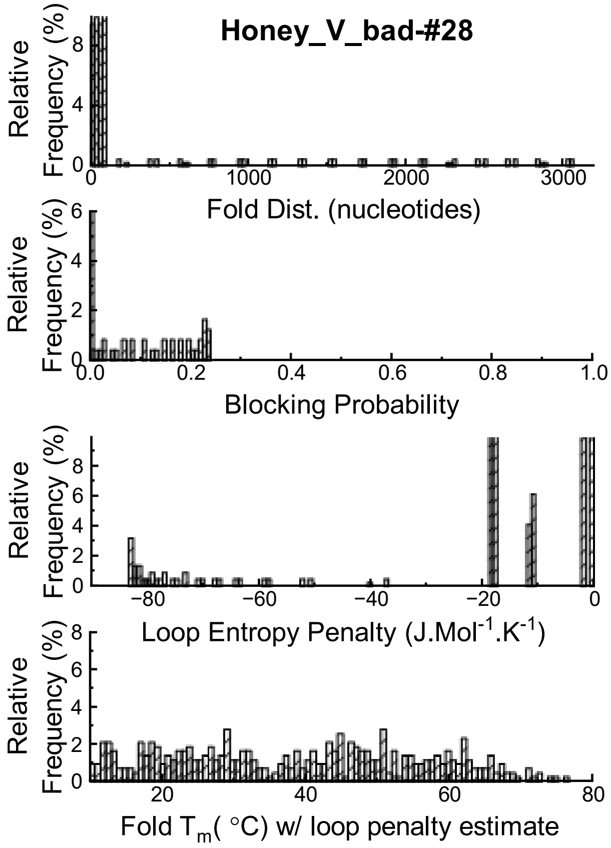

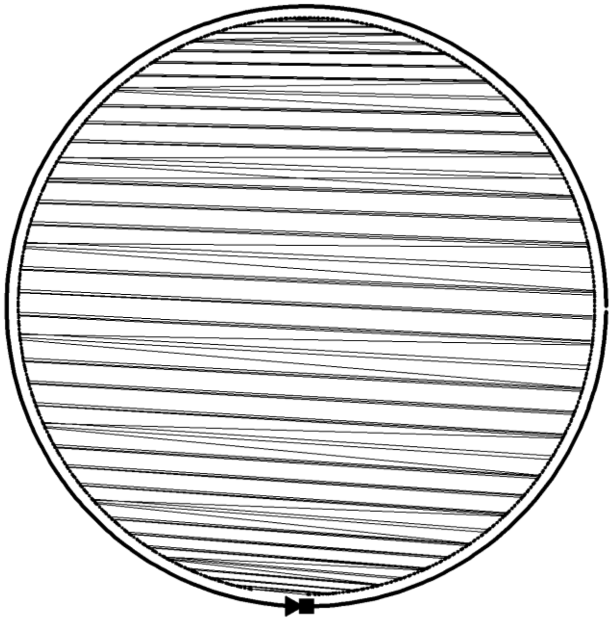

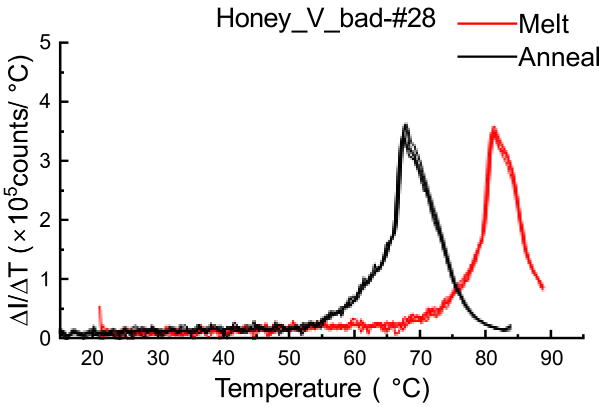

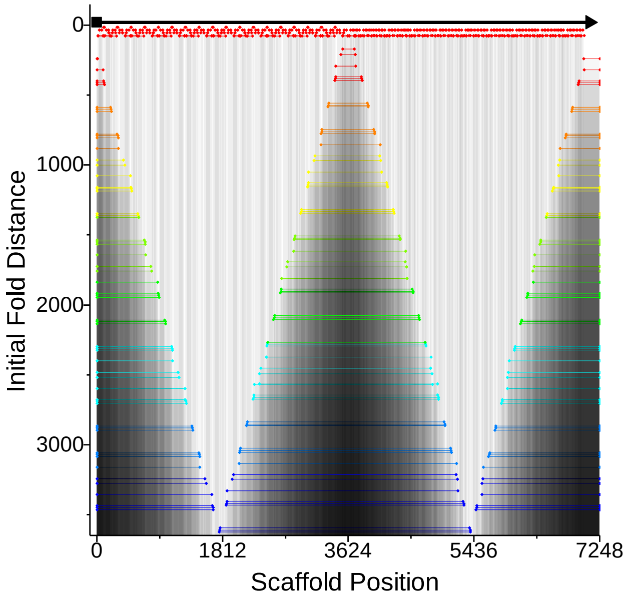

### Design #29, hex Horizontal, mix staples

**ϕ – 20.4 ± 3.4**

**W= 7±2 %; D = 12±3 %; R= 60±3 %, C= 21±3 %**

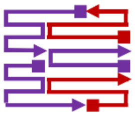

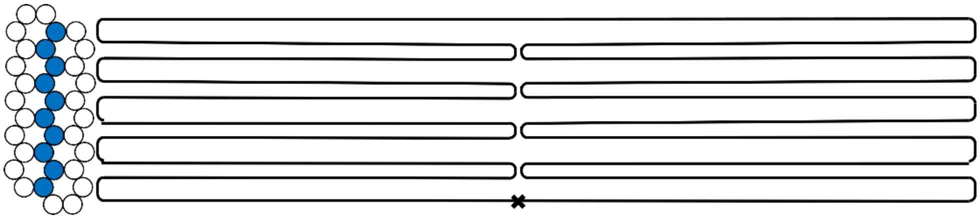

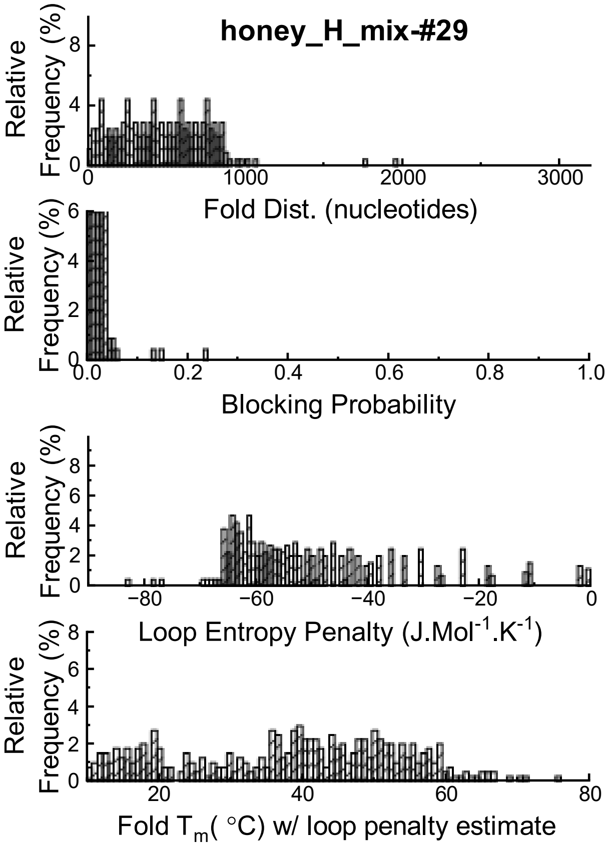

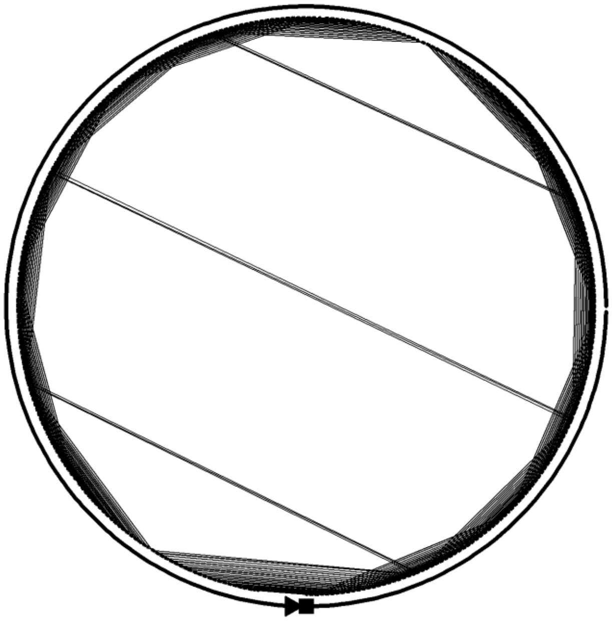

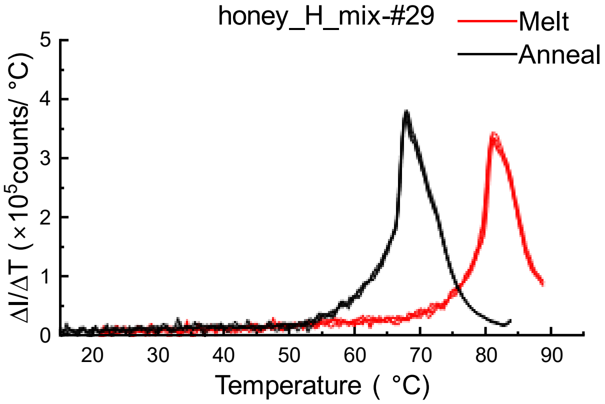

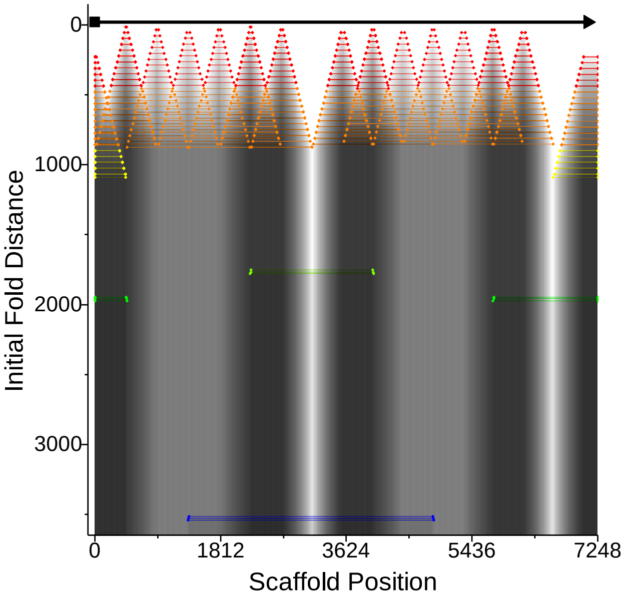

### Design #30 - Horizontal – 16-16 staple

**ϕ – 10.7 ± 3.3**

**W= 12±3 %; D = 42±4 %; R= 41±4 %, C= 5±2 %**

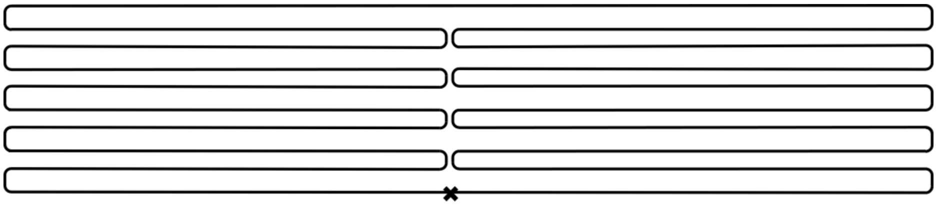

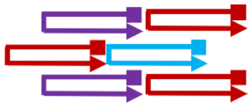

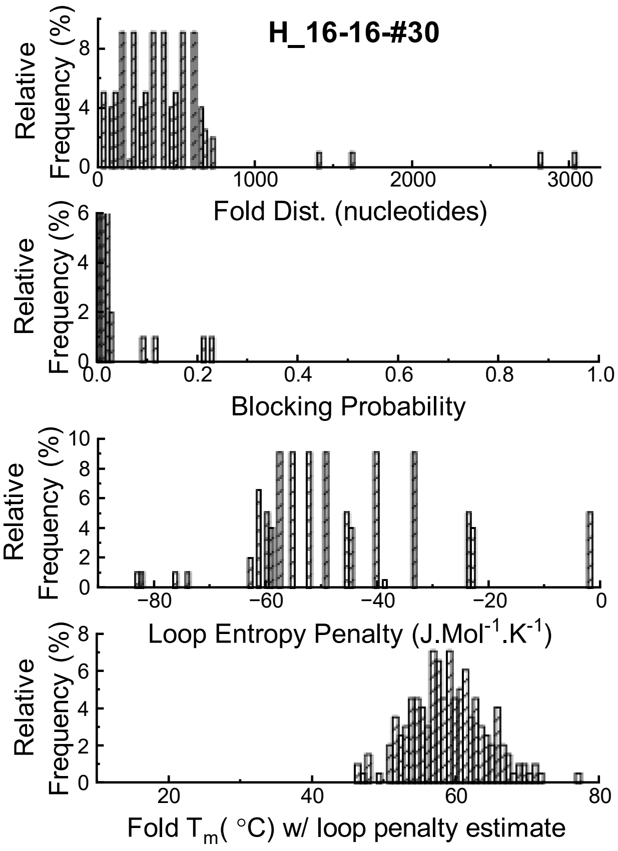

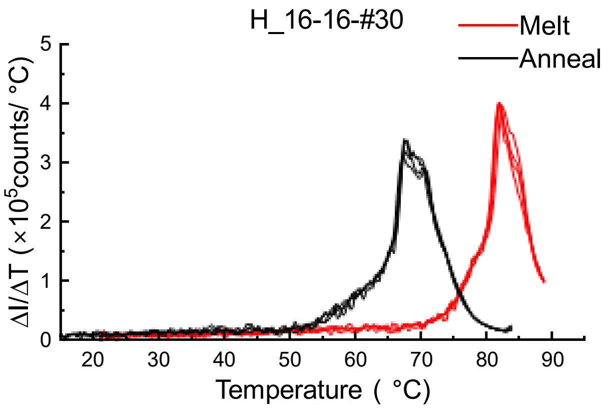

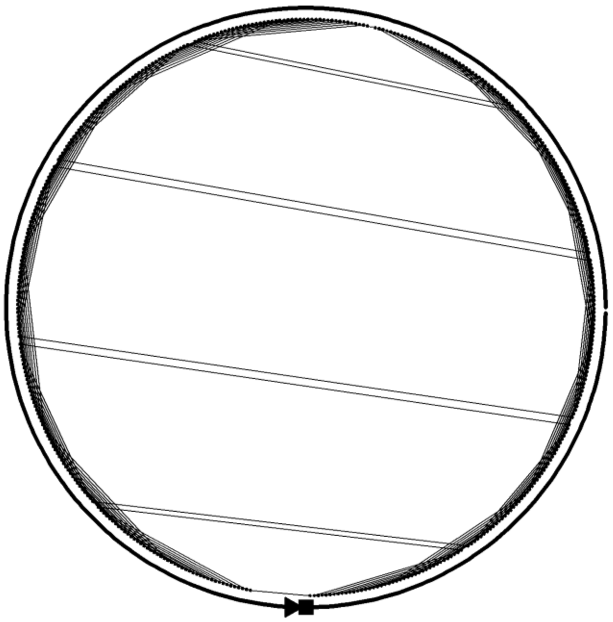

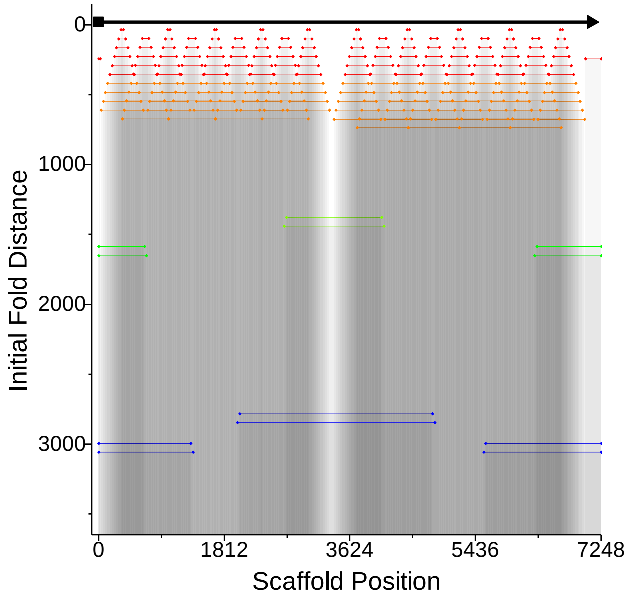

### Design # 31 - Horizontal – 16-16-16 staple

**ϕ – 8.4 ± 3.2**

**W= 31±3 %; D = 34±3 %; R= 33±3 %, C= 2±1 %**

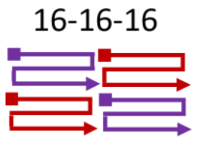

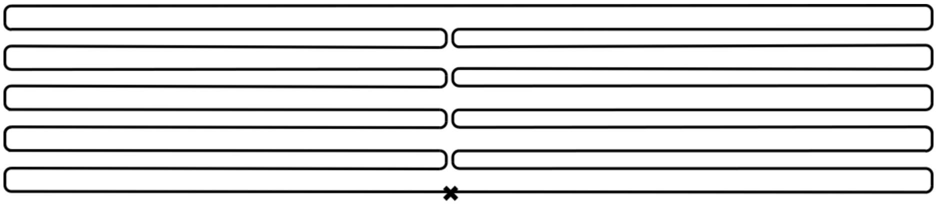

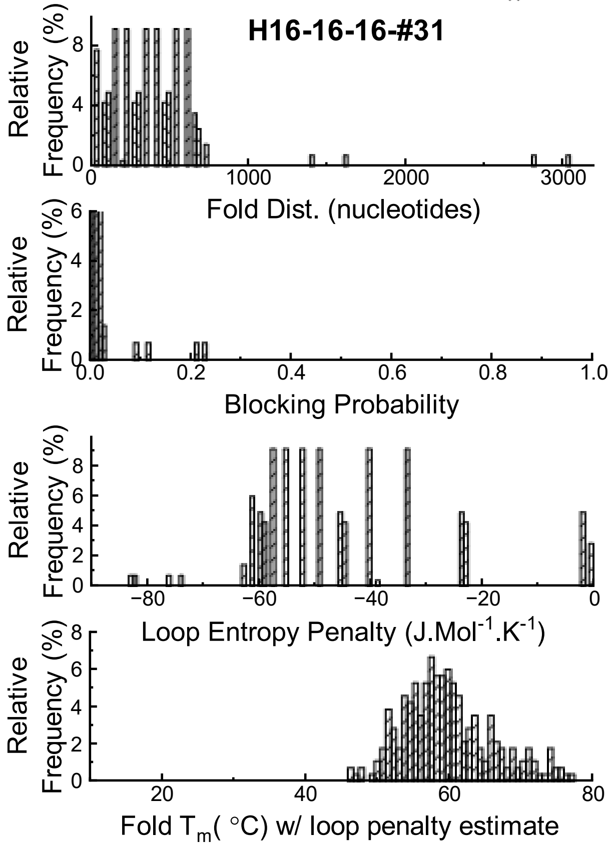

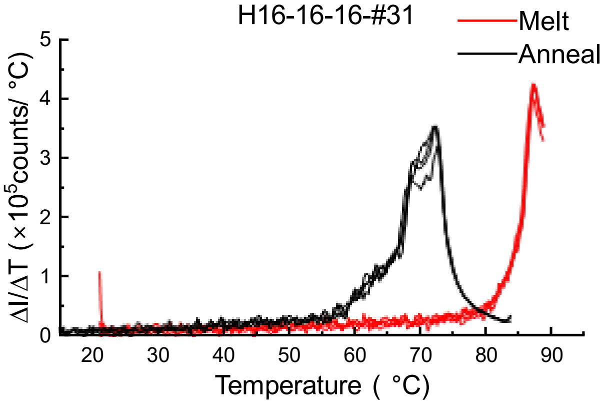

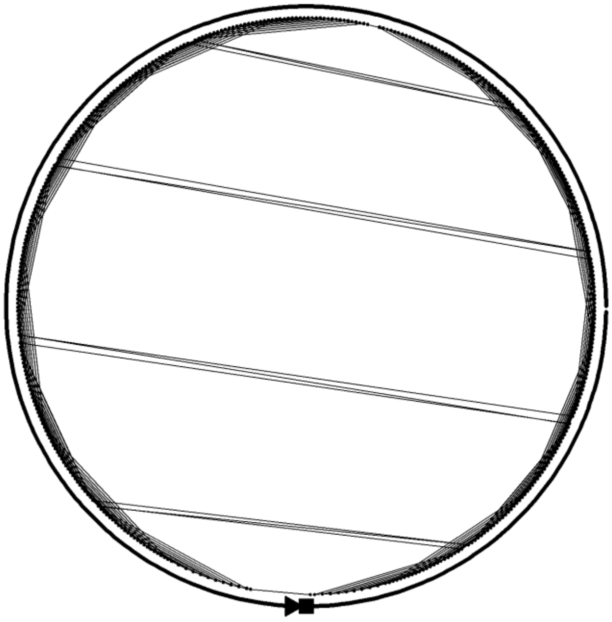

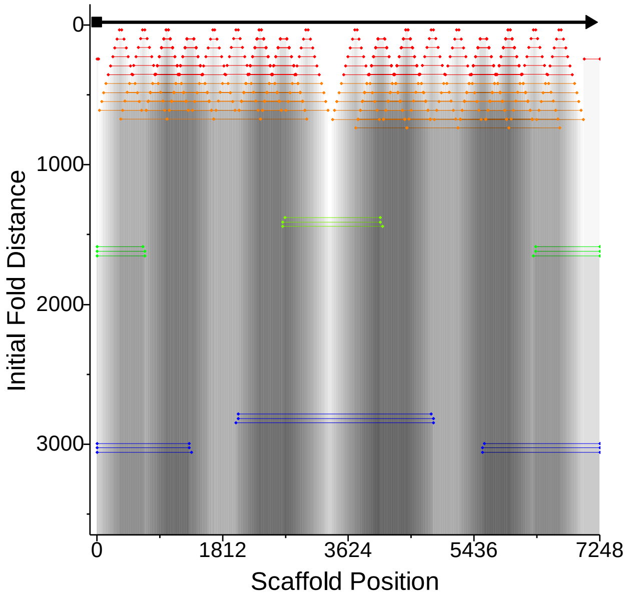

### Design # 32 - Horizontal – 8-16-16-8 staple

**ϕ – 12.0 ± 3.2**

**W= 17±2 %; D = 27±3 %; R= 50±3 %, C= 6±1 %**

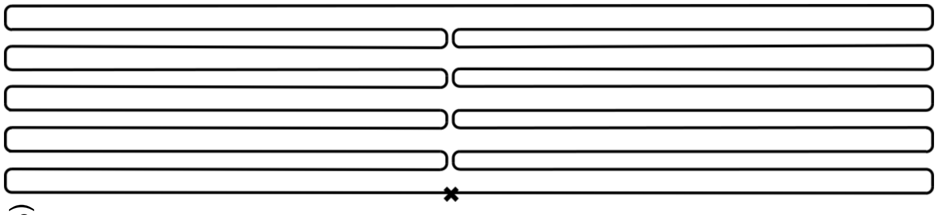

8-16-16-8

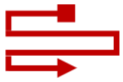

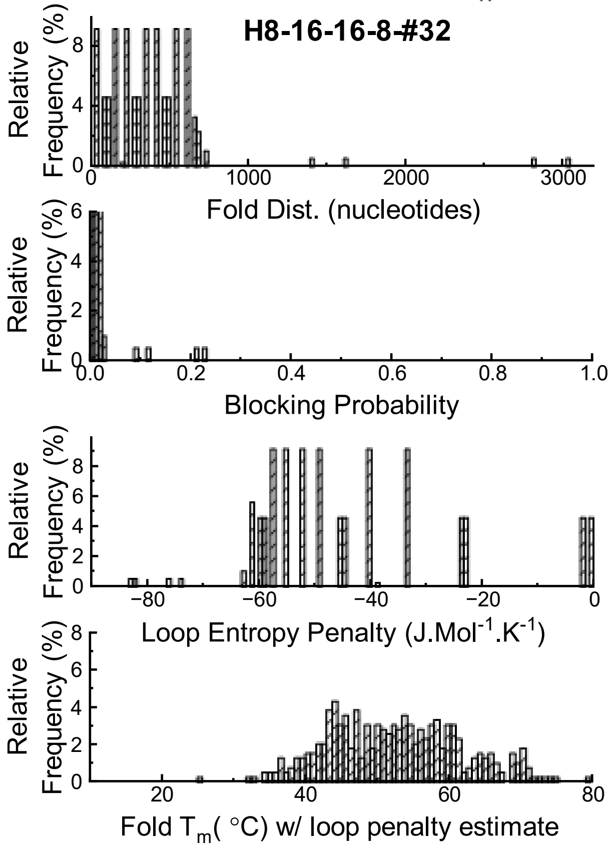

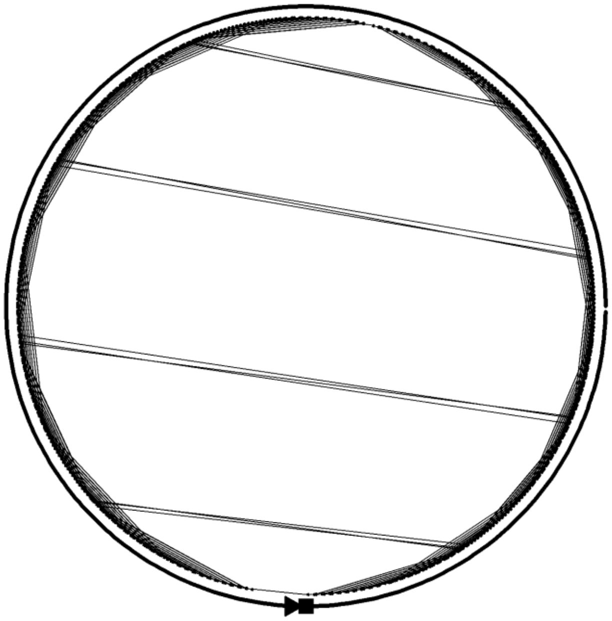

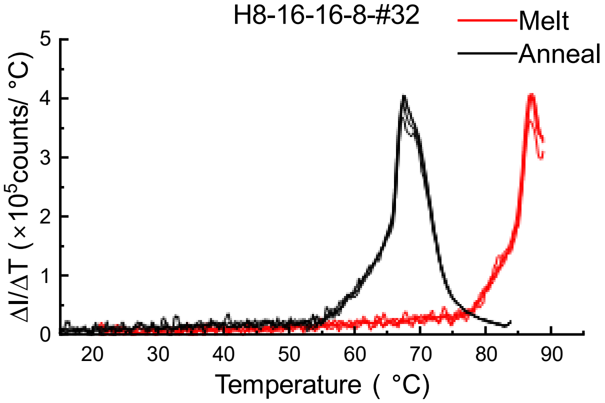

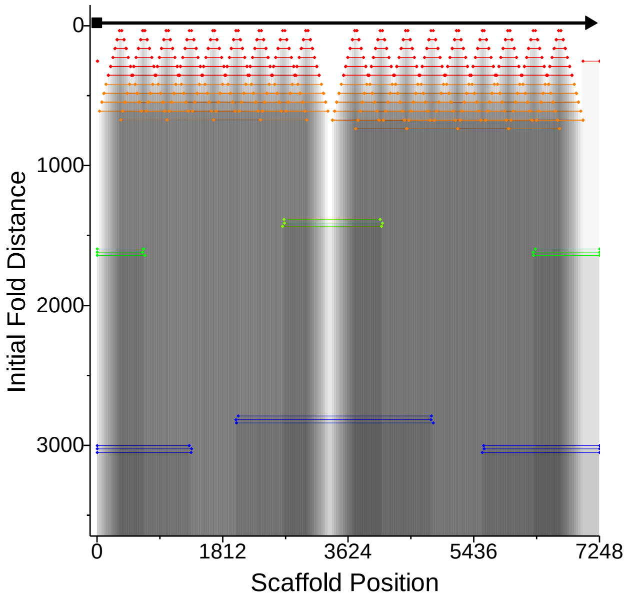

### Design #33 - Horizontal – 8-16-16-16-8 staple

**ϕ – 4.9 ± 3.3**

**W= 66±4 %; D = 18±4 %; R= 13±2 %, C= 3±1 %**

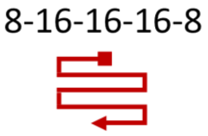

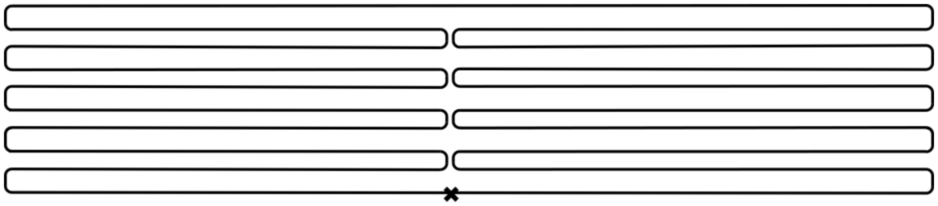

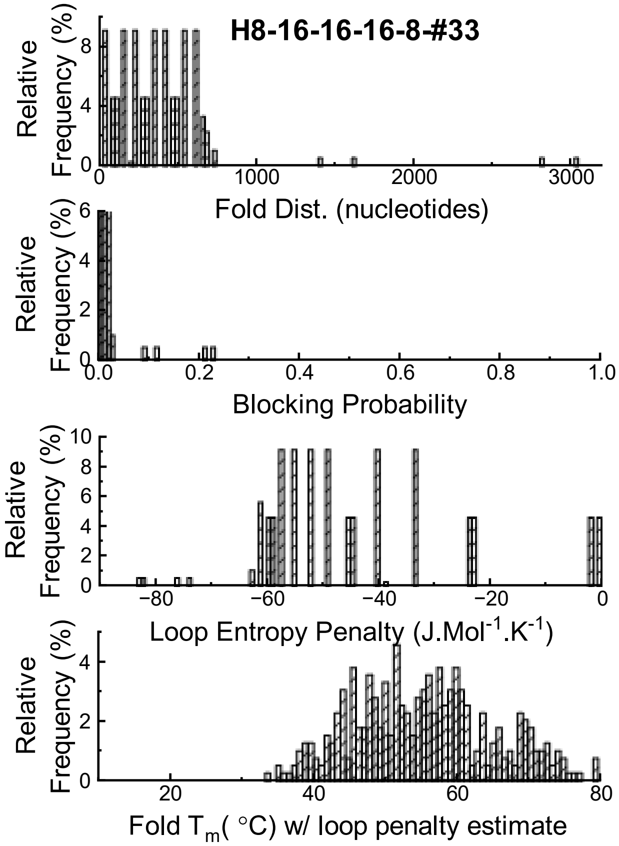

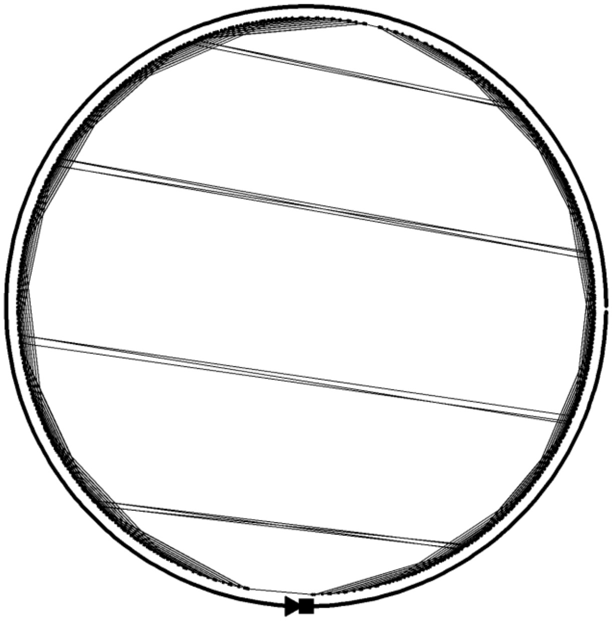

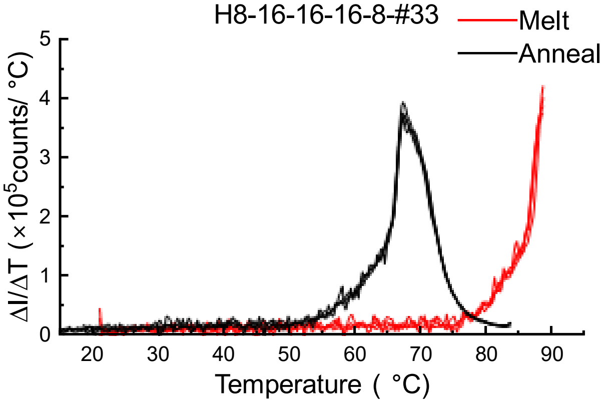

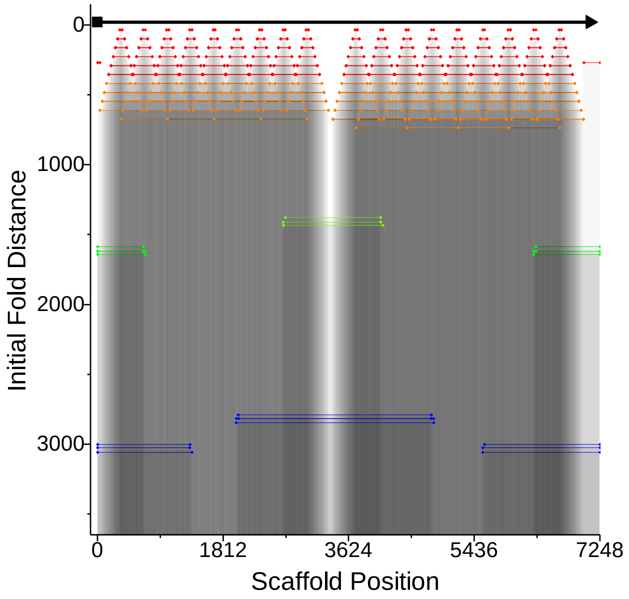

### Design #34 - Horizontal – 8-32-16-8 staple

**ϕ – 14.0 ± 3.3**

**W= 15±2 %; D= 26±2 %; R= 48±2 %, C= 11±1%**

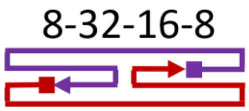

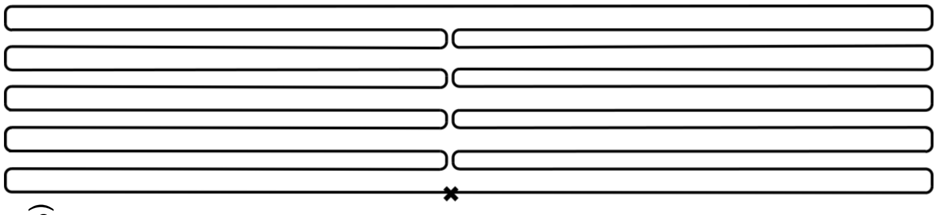

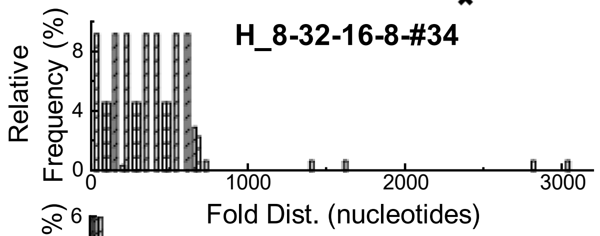

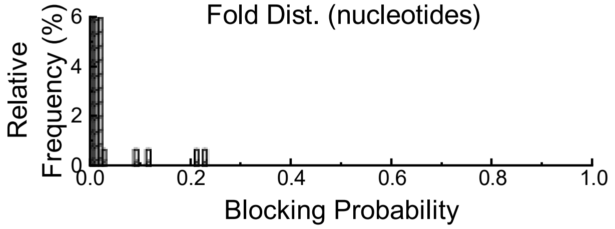

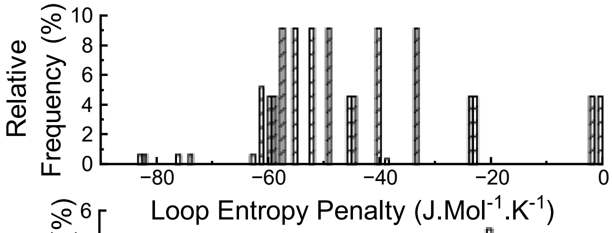

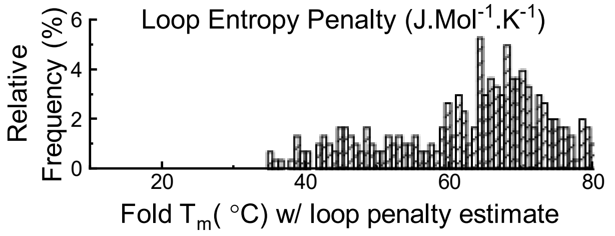

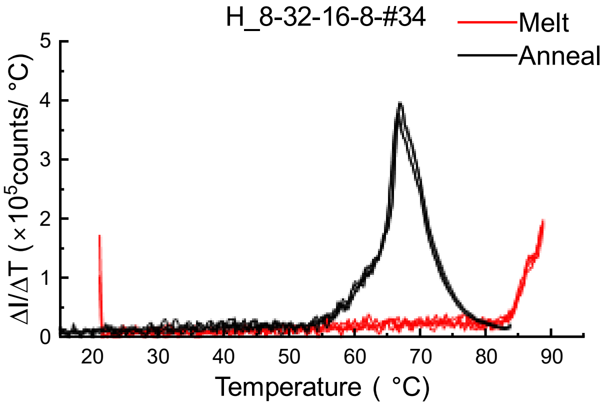

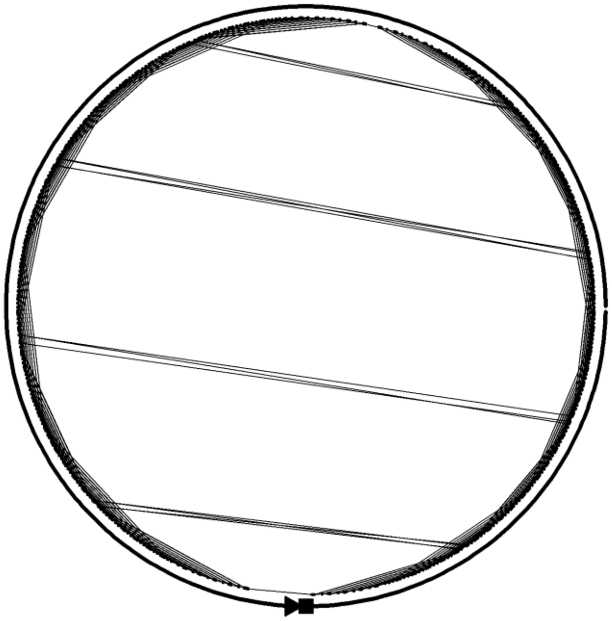

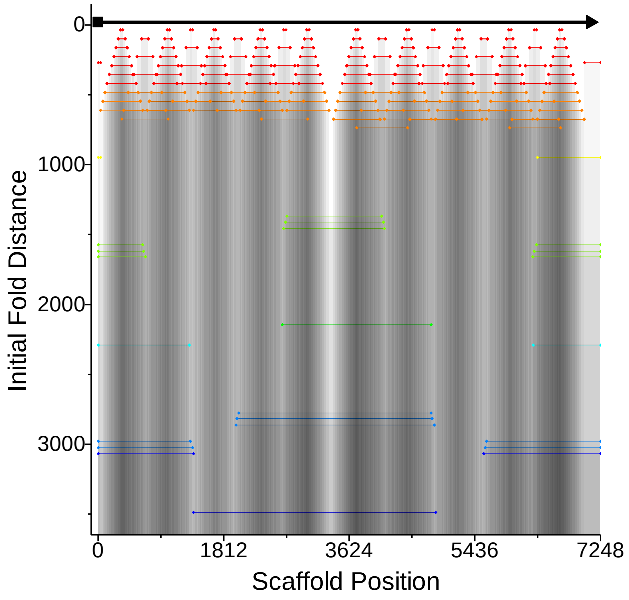
